# The diverse functions of viral microproteins

**DOI:** 10.64898/2026.08.23.746553

**Authors:** Yi Hua Chen-Neumann, Louisa Grauvogel, Adam S. Zhang, André C. Michaelis, Tim Heymann, Elisabeth Zollbrecht, Paulina Engel, Christian Pensl, Kathryn E. Yost, Wesley Suen, Sumita Sinha, Edwin N. Neumann, Stephen J. Elledge, Karl-Peter Hopfner, Benjamin E. Gewurz, Matthias Mann, Jonathan S. Weissman

**Affiliations:** Computational and Systems Biology, Massachusetts Institute of Technology; Cambridge, MA 02139, USA; Whitehead Institute for Biomedical Research; Cambridge, MA 02142, USA; Max-Planck Institute of Biochemistry; Martinsried, Germany; Gene Center, Department of Biochemistry, LMU Munich; Munich, Germany; Department of Biology, Massachusetts Institute of Technology; Cambridge, MA 02139, USA; Division of Genetics, Department of Medicine, Howard Hughes Medical Institute, Brigham and Women’s Hospital; Boston, MA 02115, USA; Cluster for Nucleic Acid Sciences and Technologies – NUCLEATE; Butenandtstraße 1, 81375 Munich, Germany; Division of Infectious Diseases, Mass General Brigham Hospital; Program in Virology, Harvard Medical School; Center for Integrated Solutions to Infectious Diseases, Broad Institute.; NNF Center for Protein Research, University of Copenhagen; Copenhagen, Denmark; Howard Hughes Medical Institute, Massachusetts Institute of Technology; Cambridge, MA 02142, USA; Koch Institute for Integrative Cancer Research, Massachusetts Institute of Technology; Cambridge, MA, USA

## Abstract

A large fraction of the uncharacterized proteome consists of microproteins. Viruses, rich in microproteins with potent activities, provide a unique system for exploring their functions and evolution. Here, we systematically annotate viral microproteins through a phenotype-first approach: construction of a >50,000-element small ORF library spanning the microprotein-coding potential of human-infecting viruses, followed by broad screens that enrich candidates for high-resolution profiling by Perturb-seq, mass spectrometry, AI-guided structure prediction, and mechanistic studies. This atlas identified >2,000 active microproteins that modulate host pathways through diverse mechanisms, including molecular mimicry, hijacking of key regulators, and altering subcellular localization. We further show the oncogenic potential of BNLF2b, an uncharacterized Epstein-Barr virus microprotein linked to nasopharyngeal carcinoma. Three principles emerge: viral microproteins are diverse and often multifunctional, show widespread convergence toward shared functions, and serve as reservoirs of evolutionary innovation. These findings elucidate how viral microproteins occupy a large functional landscape, establish them as versatile tools for host manipulation, and provide a framework for functional annotation of uncharacterized proteins, contributing toward predictive systems virology.

## Main Text

Genome sequencing has outpaced our ability to identify and annotate the functional elements it encodes. This annotation gap is especially acute in viruses, where gene-finding is confounded by dense information content, rapid mutation, and overlapping or discontinuous reading frames.

Recent high-throughput genomics and proteomics methods have further exposed the limits of standard annotation pipelines, uncovering pervasive translation of unannotated sequences across viral (*1–3*) and human genomes (*4–7*). Beyond gene-finding, the difficulty of establishing safe, tractable model systems has hindered the experimental characterization of both annotated and newly discovered viral coding sequences. Consequently, the viral dark proteome, encompassing both translated-but-unannotated and annotated-but-uncharacterized sequences, remains largely unexplored.

The scale of the viral dark proteome has been revealed by ribosome profiling (Ribo-Seq), which captures actively translated sequences at codon-level resolution (*8*). Across many viruses—including herpesviruses (*1*, *9–11*), influenza (*12*), SARS-CoV-2 (*2*), HIV-1 (*13*), and most recently a synthetic pan-viral library (*3*)—ribosome profiling has identified thousands of non-canonical open reading frames (ncORFs) mapping outside annotated protein-coding genes.

These ncORFs are translated through non-canonical mechanisms such as non-ATG initiation, ribosomal frameshifting, leaky scanning, stop-codon readthrough, and cap-independent translation, which viruses depend on to control gene expression and tune protein stoichiometry (*14–20*). ncORFs from viruses and cancer are a rich source of immunogenic peptides (*21–24*). However, how broadly the thousands of small ORFs (smORFs) translated through these mechanisms represent genuine coding sequences that yield functional microproteins is an open question.

Despite their small size, microproteins (<120aa) are not mere translational noise or functionless byproducts of viral evolution. Studies of a few well-characterized viruses reveal that microproteins can play a central role in host-pathogen biology. In HIV, nearly half (7/15) of annotated genes encode microproteins (*25*), many of which are non-canonically translated (*15*, *26*) or have poor conservation (*27*). Study of these microproteins has driven critical discoveries in host-pathogen biology, including the identification of tetherin, a host-restriction factor inhibited by Vpu (81aa) to enhance viral particle release and spread (*28*). Likewise, DCAF1 (VprBP) was identified through discovery that Vpr (98aa) hijacks this E3 ligase adapter to arrest cells in G2 and promote viral replication (*29*). Moreover, a substantial fraction (10/29) of annotated SARS-CoV-2 proteins are microproteins with critical roles across the entire viral life cycle (*30*). These insights are derived from these two intensively studied pandemic viruses, but the repertoire of functional microproteins in the broader virome including zoonotic viruses, remains unmapped.

The functional potential and pervasiveness of viral smORFs raise the possibility that they are reservoirs of newly emerged sequences that act as raw substrates for *de novo* gene evolution. While gene evolution via reuse-based mechanisms—duplication, fusion, and horizontal gene transfer— is well documented (*31*, *32*), *de novo* emergence from non-coding sequences remains less understood, owing to the difficulty of identifying such events (*33*). Although dozens have been cataloged (*34*, *35*), large-scale exploration of how *de novo* genes emerge and evolve activities has been out of reach. High mutation rates, unconventional expression, and intense selection pressures make viruses uniquely suited for studying *de novo* genes. In this context, viral smORFs offer a powerful model, especially since known viral microproteins, much like *de novo* genes, are often lineage-restricted and bear signatures of rapid evolution such as divergent functions among homologs (*27*). Pairing systematic functional characterization with phylogenetic analysis of viral smORFs could therefore capture the full cycle of gene birth and death to identify the key parameters driving *de novo* gene evolution.

Together, these considerations suggest viral microproteins are a rich landscape for exploring ncORFs, host-virus biology, and gene evolution. Multiple technological advances now enable systematic, high-throughput annotation of viral microproteins. First, high-fidelity synthesis of <400nt oligonucleotides permits construction of libraries containing tens of thousands of smORFs, bypassing the need to work directly with viruses and allows rapid characterization of the host phenotypes they impact (*3*, *36–38*). Second, high-dimensional functional genomics can now be implemented at a scale capable of capturing the full range of possible host pathways that viruses co-opt. Particularly enabling is Perturb-seq, which couples genetic perturbations, including ORF overexpression, with single-cell RNA sequencing (scRNA-seq). By eliminating the need to select predefined phenotypes prior to screening, Perturb-seq provides an unbiased platform for discovering host-virus interfaces. Beyond capturing a broad range of phenotypes, its high-resolution output can nominate specific molecular mechanisms that explain the cellular changes detected by genetic screens for growth, surface-marker expression, or reporter activity (*39*). In parallel, increased sensitivity and throughput of proteomics have transformed affinity enrichment–mass spectrometry (AE-MS) from a labor-intensive, single-bait technique into a screening platform for mapping the molecular interactions underlying a phenotype (*40*). Together, these complementary methods provide a systematic framework for annotating the pan-viral dark proteome with unprecedented breadth, depth, and scale.

In this study, we generate a systematic functional atlas of viral microproteins that spans the full spectrum of human-infecting viruses. Our phenotype-first approach uncovers a large functional landscape that emerges from the diverse, modular arrangement of compact building blocks that include small structural domains and short linear motifs (SLiMs), which we refer to as “molecular keys”. We also document widespread convergence of microproteins with shared functions throughout the virome, some arising through independent evolution of similar molecular keys. These observations suggest that viruses exploit their vast dark proteomes to generate new functional sequences, fueling rapid innovation in the host–virus arms race. These themes are well illustrated in our identification and mechanistic dissection of the oncogenic properties of BNLF2b, a lineage restricted and previously uncharacterized Epstein–Barr virus (EBV) microprotein linked to nasopharyngeal carcinoma (*41*). Altogether, this work elucidates the principles that give rise to microprotein functions, provides a framework for systematic functional annotation, and uncovers vulnerable host nodes susceptible to viral targeting.

## Results

### A funnel strategy for the identification of functional microproteins

Identifying functional microproteins is a “needle in a haystack” problem: any long DNA sequence contains many ORFs by chance (*42*), yielding large numbers of potentially functional sequences. To balance trade-offs between cost, tractability, and scale, we adopted a funnel strategy that begins with permissive curation of a pan-viral library of 52,979 smORFs and progressively filters to smaller, enriched libraries. First, all smORFs, based solely on the presence of initiation and stop codons, were extracted from the genomes of viruses documented to infect human cells (Fig. 1A). For the initial screens, we chose growth as a phenotypic readout, reasoning that this approach would capture hits spanning a wide variety of functions. Two cell lines, K562 leukemia cells and non-transformed hTERT-immortalized RPE-1 cells, were selected for their robustness in high-throughput screens and to broaden the spectrum of functional candidates, including growth-promoting genes detectable only in a non-transformed background. Top-scoring candidates from these primary screens were resynthesized into the Enriched Library comprising 4,107 elements, for which additional genetic screens and Perturb-seq were performed. Finally, a subset of hits was selected for arrayed AP-MS and specialized follow-up assays. This permissive-to-stringent approach was designed to achieve functional characterization with both breadth and depth.

**Fig. 1.**
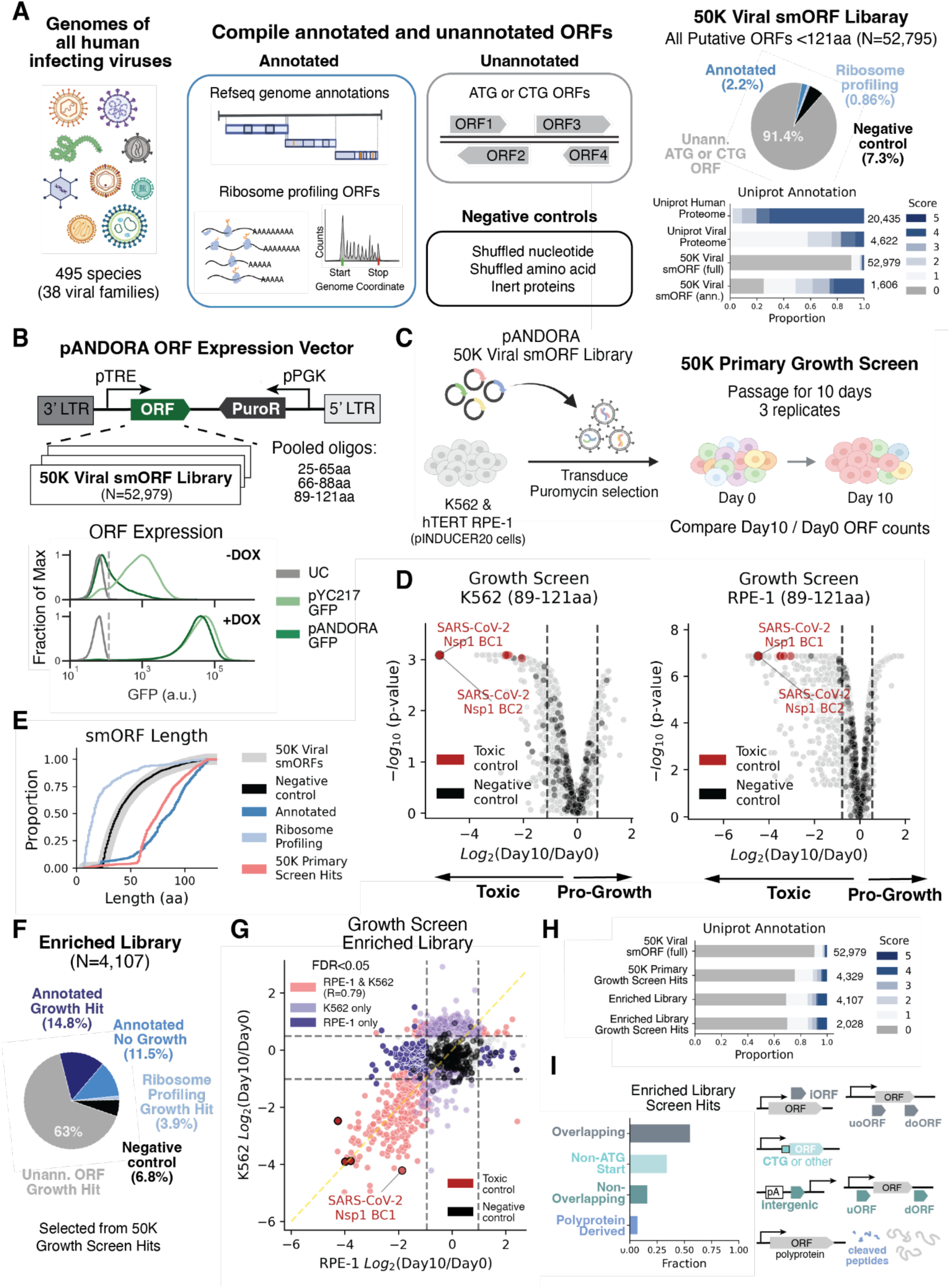
A pan-viral microprotein library for the identification of functional viral genes. (**A**) Schematic of 50K Viral smORF library design pipeline. Candidates were sourced from annotated genes, ribosome-profiling data, and unannotated ORFs with ATG or CTG start codons. Shuffled sequences and inert proteins were included as negative controls. Fractions of library elements in each subpool are shown as a pie chart. UniProt annotation scores were retrieved for all library elements shown alongside the annotated human and viral proteomes. Elements absent from the database are assigned a score of zero. (**B**) The pANDORA lentiviral vector was used to inducibly express a pooled oligo library of 52,979 putative smORFs with minimal leakiness. Oligos encoding smORFs were synthesized in three oligo pools distinguished by length (25-65aa, 66-88aa, and 89-121aa). (**C**) Experimental workflow for primary viral smORFeome screens. 10-day growth screens were performed in cancer-derived K562 and hTERT-immortalized RPE-1 cells that express the pINDUCER20 system. (**D**) Volcano plots of primary growth screens. Dashed lines are mean +/- two standard deviations of negative control elements. (**E**) Cumulative distribution of amino acid length. (**F**) Validation library design. (**G**) Correlation of validation growth screen results between K562 and RPE-1. Pearson R=0.78 for elements with negative log2 fold changes at Day 10 compared to Day 0. (**H**) UniProt annotation scores of growth screen hits (FDR<0.05). (**I**) Fraction of smORFs with significant growth phenotypes for indicated ORF type.

### Design and curation of the pan-viral smORF library

The initial pan-viral library of 52,979 smORFs was derived from 502 genome sequences of all human-infecting viruses cataloged in VirusHostDB (*43*), which spans 38 viral families across all seven Baltimore classifications (fig. S1; data S1). To accommodate the 400nt oligo synthesis limit at the time of this study, we restricted our curation to smORFs <121aa. The library consists of three categories: (i) annotated sequences (N = 1,606); (ii) unannotated smORFs encoding peptides ≥24aa and selected solely by the presence of an ATG or CTG start codon (N = 47,525), which made up the largest fraction of the library (89.7%); and (iii) negative controls (N = 3,848), ∼7% of the library. To assess annotation status, we asked whether each element was present in or homologous to a UniProt entry and retrieved its annotation score, which is a five-point heuristic reflecting how much annotation an entry carries (names, functional comments, sequence features, GO terms, cross-references). Scores of 3 or less mark sparsely annotated entries, with little known beyond the sequence itself (*44*). Of the 1,606 annotated smORFs, a large majority (74%) had scores of 3 or less (Fig. 1A), and 272 were polyprotein cleavage products, a common viral strategy for expressing multiple functional proteins from a single precursor (*26*). Four types of negative control sequences were designed: (i) shuffled amino-acid sequences (N = 1,446), (ii) shuffled nucleotide sequences (N = 1,428), (iii) antisense-derived smORFs from genomes with defined directionality (N = 934), and (iv) 40 inert peptides composed of common linkers and epitope tags. To minimize oligo costs, sequences were synthesized as three pools grouped by length (24-65aa, 66-88aa, and 89-121aa).

### Optimization of the pANDORA system for inducible expression of toxic ORFs

A major challenge of high-throughput ORF screens is that, unlike gRNAs in CRISPR-based approaches, ORFs have inherent activities that can disrupt the screening pipeline, including during lentiviral production. This is particularly acute for highly toxic ORFs that can drop out before assay measurement. Indeed, standard constitutive (pConstitutive) and inducible (pInducible) vectors failed to yield viable titers for highly toxic genes (fig. S2), artificially shrinking phenotypic effect sizes in pooled screens with benchmarks such as SARS-CoV-2 Nsp1 (fig. S3), a host-shutoff factor that potently inhibits translation (*45*).

To overcome this technical limitation, we developed a lentiviral vector that enables tightly controlled doxycycline-inducible ORF expression using the Tet-On system in cell lines expressing the rtTA transactivator (*46*). This yielded the pANDORA vector, which shows minimal basal leakiness throughout the screening pipeline (Fig. 1B, fig. S4). In pooled screens, pANDORA prevented premature dropout of toxic elements prior to induction and captured significantly larger phenotypic effect sizes for Nsp1 and other toxic ORFs (fig. S3). Although its antisense configuration reduced lentiviral titers by ∼50-fold (fig. S4D), pANDORA maintained robust and uniform representation of library elements. We therefore cloned the synthesized oligo pools to generate the final pANDORA 50K library.

### Caveats and benchmarks for ORF overexpression screens

Because our pipeline relies on transgenic expression to interrogate gene function, ectopic expression can be below or above physiologically relevant levels and confound phenotypic characterization. Although the pTRE promoter drives high expression (*47*), it may still fail to accumulate particularly short or highly unstable microproteins, potentially yielding false negatives. Conversely, non-physiological overexpression can produce artifacts, although this risk may be less pronounced for viral genes, which are often the most highly expressed genes during acute infection (*2*, *48*).

To address these caveats, we built several benchmarks into our experimental design. First, to control for overexpression artifacts, we compared phenotypic effect sizes against negative controls spanning the length and biophysical properties of the pANDORA 50K library. These controls establish a baseline to distinguish specific functional activities from stress phenotypes caused by misfolding or protein aggregation. Second, we leveraged well-characterized positive controls to validate our approach’s ability to capture established biological functions. For instance, if our screens recapitulated the pro-proliferative impact of the 97aa human papillomavirus (HPV) E7 oncoprotein (*49*), this would demonstrate that pANDORA drives expression in a regime relevant to its native biological context. Finally, integrating multiple orthogonal assays generates complementary evidence to support functional assignments.

### Growth screens identify toxic and pro-growth smORFs

To enrich for functional viral smORFs, we conducted 10-day pooled growth screens with the pANDORA 50K library in K562 and RPE-1 cells in triplicate (Fig. 1C). The assay detected known toxic elements, including the positive control SARS-CoV-2 Nsp1, while maintaining a low background rate from negative controls (Fig. 1D; fig. S5). Known oncogenes including HPV E7 homologs were among the top-scoring pro-growth elements in RPE-1 screens (data S2).

These benchmarks gave confidence to our ability to detect active smORFs. Implementing an effect-size cutoff of at least two standard deviations from negative controls, 4,329 primary hits (8.2% of the library) were identified, with over 90% longer than 50aa versus only 25% of elements in the overall library (Fig. 1E).

Based on these primary screens, we designed a secondary library (the Enriched Library) comprising 4,107 smORFs (Fig. 1F), selected based on the following criteria: (i) toxic hits scoring in both K562 and RPE-1 screens; (ii) pro-growth hits scoring in either screen; and (iii) all annotated genes, regardless of growth phenotype scores. New negative controls were redesigned and redundant sequences sharing >95% amino-acid identity were removed. We then repeated the 10-day growth screen in K562 and RPE-1 cells with four replicates, identifying 2,028 hits (FDR < 0.05) in either screen.

Consistent with cell-type-specific growth requirements, few pro-growth hits overlapped between K562 and RPE-1 cells, whereas toxic hits (LFC<0 in either screen) correlated (Pearson R = 0.78) in both screens (Fig. 1G). Although this secondary screen enriched for annotated elements relative to the library baseline, 20% had annotation scores of only 1 or 2, and 70% were unannotated (Fig. 1H). Furthermore, many of these smORFs exhibited features of non-canonical translation: 55% overlap with annotated genes, and 34% have non-ATG start codons (Fig. 1I).

### Additional microprotein functions captured by targeted screens and Perturb-seq

Although growth screens identified thousands of putative viral microproteins, the majority (58%) of annotated smORFs induced no significant growth phenotype (939/1,606), suggesting many modulate host pathways independent of cell proliferation (data S2, 3). To identify smORFs acting on growth-independent processes, we performed targeted screens for MHC class I (MHC-I) antigen presentation and survival under ER stress. We prioritized MHC-I presentation given its essential role in adaptive antiviral immunity, screening for modulators of surface expression via FACS-based β2m staining in RPE-1 cells transduced with the Enriched Library, with or without IFN-γ stimulation. We also screened A375 cells expressing the 66-88aa smORF library (N=6,267), as it did not undergo the growth-phenotype filter of the funnel. Because the ER is central to the MHC-I pathway—which involves heavy chain and β2m light chain assembly, peptide cargo loading, and maturation in the ER followed by trafficking through the ER and Golgi secretory pathway for presentation to CD8+ T cells (*50*)—we reasoned that survival under ER stress would offer a complementary readout, separating smORFs that impact different steps of the pathway. In parallel, we performed Perturb-seq on the entire Enriched Library for high-dimensional transcriptional profiling (Fig. 2A) using a newly optimized, more scalable probe-based method built on the 10x Genomics Gene Expression Flex platform for scRNA-seq (fig. S6), also used in other recent studies (*51*, *52*). Combining specialized reporters with unbiased, high-resolution methods enables exploration of a wide variety of viral microprotein functions.

**Fig. 2.**
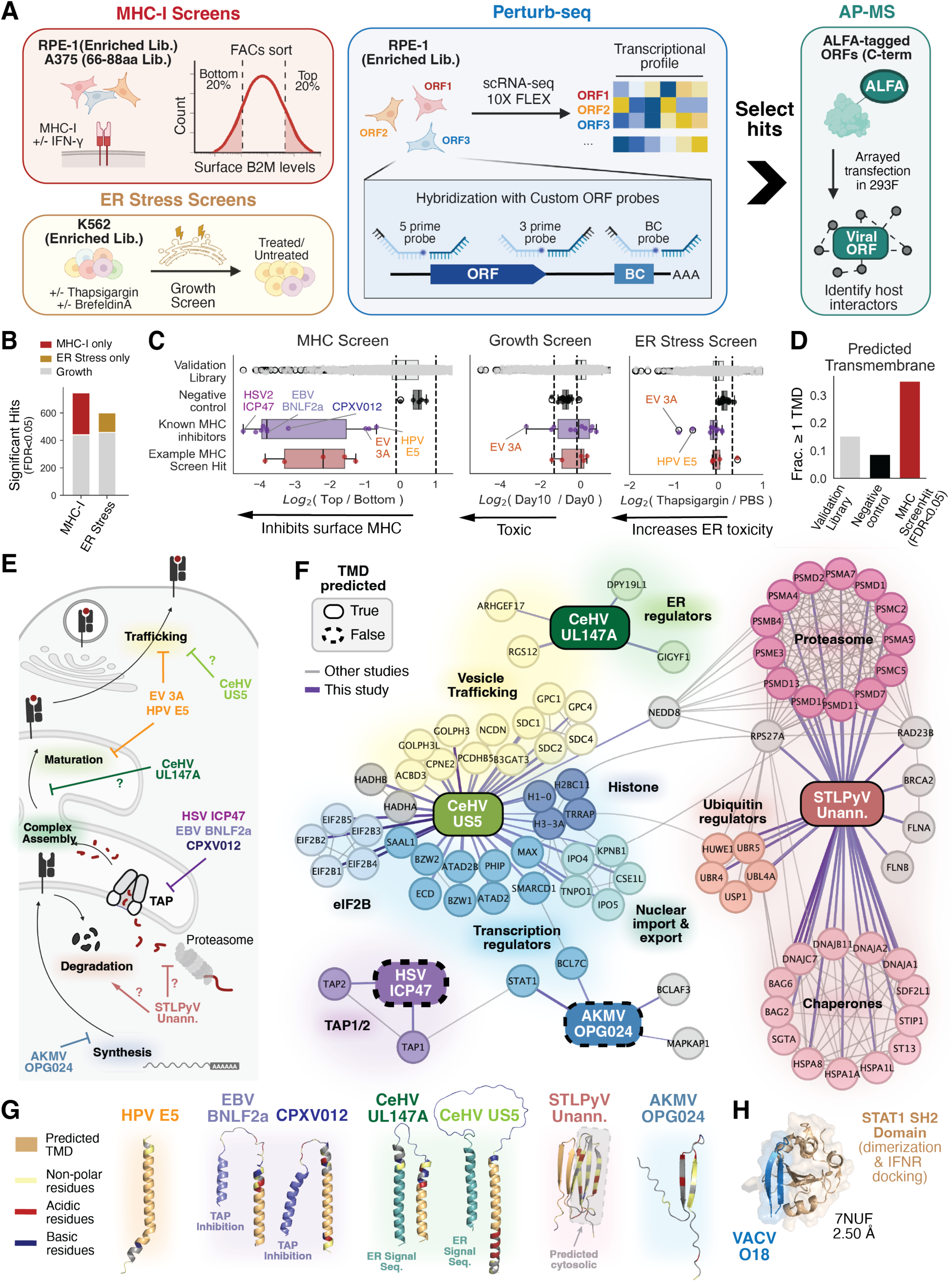
A multi-omic functional map of the viral smORFeome reveals diverse mechanisms of host cell perturbation. (**A**) Schematic of assays used to characterize smORF functions beyond growth. Screens for FACS-based MHC-I surface expression were performed with or without IFN-γ stimulation in RPE-1 cells with the enriched library or in A375 cells with the 66-88aa library. Screens for survival under ER stress were induced by thapsigargin in K562 cells. Perturb-seq was performed in RPE-1 cells using 10x Genomics Flex with custom probes targeting viral smORFs to link viral ORF overexpression to transcriptional profile. High-confidence hits were selected for high-throughput AE-MS identification of host protein interactors. (**B**) Number of hits identified from the MHC-I and ER-stress screens, with the number of smORFs inducing significant growth phenotypes in grey. (**C**) Effect sizes in the MHC-I, growth, and ER-stress screens for known and newly identified MHC-I inhibitors. (**D**) Fraction of MHC-I screen hits with at least one PHOBIUS-predicted TMD. (**E**) Schematic of the MHC-I presentation pathway, with the steps targeted by viral MHC-I inhibitors shown. (**F**) Network of significant interactors identified by anti-ALFA AE-MS in 293F cells 48 h after transfection with plasmids encoding ALFA-tagged MHC-I inhibitors. Purple edges are significant interactions from this study; edge width corresponds to significance and edge darkness to fold change. Grey edges are high-confidence interactions (>0.7) from the STRING database. (**G**) AF3 structure predictions of viral microproteins with TMDs, highlighted by residue property. (**H**) X-ray structure of VACV O18, a homolog of AKMV OPG024, bound to STAT1 (PDB: 7NUF).

### Identification of uncharacterized viral MHC inhibitors

Of 8,806 smORFs screened, 748 viral smORFs passed the significance threshold (FDR < 0.05) for altered surface MHC-I levels. Many hits for the MHC-I screen did not score in growth screens (Fig. 2B), supporting that functional smORFs may have activities that do not impact cell growth. The screens recovered well known inhibitors, including the TAP inhibitors HSV ICP47, EBV BNLF2a, and CPXV012 (*53*), as well as the ER disruptors HPV E5 (*54*) and enterovirus 3A microproteins (*55*, *56*) (Fig. 2C; fig. S7).

While most MHC-I hits also scored in growth screens, a significant portion (40%) did not, suggesting the screens detected both direct and indirect modulators of antigen presentation. We therefore hypothesized that specific immunomodulators would score in the MHC-I screens, but not in the ER stress or growth screens. Indeed, TAP inhibitors were among the strongest surface MHC-I down-regulators that showed no phenotype in the other two screens. By contrast, broad ER disruptors ranked among the top smORFs sensitizing cells to ER stress and exhibited modest growth phenotypes (Fig. 2C; fig. S8). Consistent with these trends, 37% of MHC-I hits were highly hydrophobic with predicted signal sequences or transmembrane domains (TMDs) and also scored in ER stress screens, potentially by localizing and disrupting MHC-I processing in the ER (Fig. 2D; fig. S9). Prioritizing specific antigen-presentation regulators , we focused on candidates scoring exclusively in the MHC-I screens for further analysis.

### AE-MS reveals that MHC-I inhibitors interact with diverse host proteins

Because the MHC-I presentation pathway is well characterized, we reasoned that host interactors can be used to pinpoint the antigen-processing step disrupted by each microprotein. This systematic approach permits exploration of whether microproteins are restricted to a narrow set of host nodes or instead engage multiple processes. The distinction is particularly pertinent in this pathway, as four of the five TAP inhibitors discovered to date are <150aa (*37*, *53*, *57*), raising the possibility that microproteins are restricted to select host nodes accessible within their biophysical constraints.

To explore these questions, we highlight five example hits that scored exclusively in the MHC-I screens for AE-MS analysis: three have predicted TMDs (CeHV US5, CeHV UL147A, and an unannotated STL polyomavirus smORF) and two do not (AKMV OPG024 and HSV ICP47).

Our pipeline robustly recovered the established ICP47–TAP1/2 interaction (*53*), lending confidence to the diverse host interactors detected for the remaining smORFs (Fig. 2E and F; fig. S10A).

Two microproteins, CeHV UL147A and CeHV US5, were found to interact with ER or Golgi components. Top-scoring host interactors of CeHV UL147A were the ER regulators DPY19L1 and GIGYF1, implicating UL147A as a disruptor of ER protein processing (Fig. 2E and F; fig. S10B). This has parallels with a recent study showing that the homologous HCMV UL147A localizes to the ER, promotes degradation of NK stress ligands, which results in evasion of NK-cell-mediated killing (*58*). Our findings suggest that UL147A homologs act as immunomodulators targeting surface-bound immune-activating ligands to escape innate or adaptive surveillance. Similarly, CeHV US5 interacted with ER and Golgi components involved in vesicle trafficking, suggesting it interferes with maturation or transport of MHC-I complexes to the plasma membrane (Fig. 2E and Fl; fig. S10D).

The remaining MHC-I modulators interacted with host proteins involved in other steps of MHC-I presentation. The unannotated STL polyomavirus ORF associated with protein-degradation machinery, including E3 ubiquitin ligases, their regulators, and proteasome components. These interactions suggest two non-exclusive mechanisms: it acts as a molecular adapter linking MHC-I assembly factors to the degradation machinery, or it impairs proteasome-mediated antigen processing, depleting the peptide pool required for stable MHC-I assembly (Fig. 2E and F; fig. S10E). Additionally, AKMV OPG024 bound the transcription factor STAT1, which upon activation upregulates MHC class I genes and their associated antigen-processing machinery (*59*), pointing to synthesis as the step of interference (Fig. 2E and F; fig. S10C).

Together, these findings illustrate that microproteins are not confined to a single vulnerable host node, as even the five hits examined here engage at least four distinct steps of the MHC-I pathway.

### Microproteins act through short “molecular keys”

To understand what biochemical features underlie the molecular activities of MHC-I inhibitors, we leveraged domain predictions and AlphaFold 3 (AF3) structure modeling (*60*), which revealed additional “molecular keys” beyond TMDs. Though structure prediction of host–virus complexes remains challenging due to lack of coevolutionary information and poor conservation, recent systematic benchmarking has shown that accurate models are recovered when ranked by prediction confidence (*61*). We therefore restricted interpretation to high-confidence models and used interface confidence (ipTM) as our criterion throughout (fig. S9). This analysis identified small structural elements and revealed that many microprotein hits adopt bipartite architectures, pairing two distinct elements that expand functional capabilities (Fig. 2G).

The TAP inhibitors BNLF2a and CPXV012 share this bipartite design, pairing a membrane-anchoring hydrophobic TMD with a specialized TAP-inhibitory domain. Combining these domains enhances inhibition of peptide transport, as the TMD anchors the microprotein to the ER, increasing the local effective concentration of the TAP-inhibitory domain to block peptide transport (*57*). Similarly, CeHV UL147A and CeHV US5 combine a hydrophobic TMD with a predicted ER signal peptide, ensuring robust ER or Golgi localization to disrupt MHC-I complex assembly or trafficking (Fig. 2G; fig. S9).

Beyond combinations of alpha-helical keys, we also observe microproteins with predicted beta-sheet topologies. For example, the unannotated STL polyomavirus smORF is predicted to form a rigid, six-stranded beta-sandwich fold (Fig. 2G; fig. S9), with two hydrophobic strands predicted as transmembrane anchors and the remaining strands predicted to localize to the cytosol, potentially recruiting protein-degradation machinery. Likewise, AKMV OPG024 is predicted to contain a beta-hairpin motif, which AF3 predicts to bind STAT1 (ipTM = 0.78), its top-scoring host interaction from AE-MS (Fig. 2F, fig. S9). This is corroborated by a recent study showing that the VACV O18 beta-hairpin binds the SH2 domain of STAT1 (PDB: 7NUF) to inhibit its activation (Fig. 2H). This motif also appears in the non-structural V protein of the phylogenetically distant Nipah virus, demonstrating convergent evolution of a shared molecular key (*62*).

These results point to molecular keys as a common feature underlying microprotein activity, a capacity attributable to their compact, diverse, and modular designs. These keys can target critical host nodes (e.g., TAP and STAT1) and be combined, shedding light on how such small proteins can profoundly alter host phenotypes. Furthermore, molecular keys with similar features have independently evolved multiple times across the virome, suggesting that large numbers of short sequences have been produced and selected throughout viral evolution. These observations raise the question of whether these themes generalize to the full repertoire of activities adopted by viral microproteins and what new ones may emerge in other contexts.

### Perturb-seq identifies hundreds of smORFs that induce strong transcriptional phenotypes

To chart the broader landscape of host pathways engaged by viral microproteins beyond MHC-I presentation, we performed Perturb-seq on RPE-1 cells transduced with the 4,107-element Enriched Library. As benchmarks for our newly optimized Perturb-seq method and as anchors for interpreting transcriptional responses, we designed 18 positive controls composed of well-characterized transcription factors and regulators that drive robust transcriptional phenotypes.

These controls activate a range of pathways, including the unfolded protein response (UPR; XBP1s, ATF6n, nSREBP1), the integrated stress response (ISR; ATF4, DDIT3, CEBPB), immune signaling (IκBα, STING/TMEM173), and RNA metabolism (HNRNPH2). The viral host-shutoff factor SARS-CoV-2 Nsp1 was also included to assess representation and detection of highly toxic library elements.

To evaluate the specificity of our method, we established a reference group of randomly selected cells expressing negative control ORFs and compared their transcriptional profiles against those of smORF perturbations. Two metrics were used to assess transcriptional phenotype strength: pseudobulk differential gene expression (DGE) and the energy-distance test, which determines whether a perturbation significantly alters cell states by calculating cell-to-cell distances in transcriptional space within and between experimental groups (*39*, *63*). Both behaved as expected: all negative controls had <5 differentially expressed genes (DEGs) and did not pass the energy-distance test (FDR < 0.05), whereas all 18 positive controls passed the energy-distance test and had >10 DEGs (Fig. 3A).

**Fig. 3.**
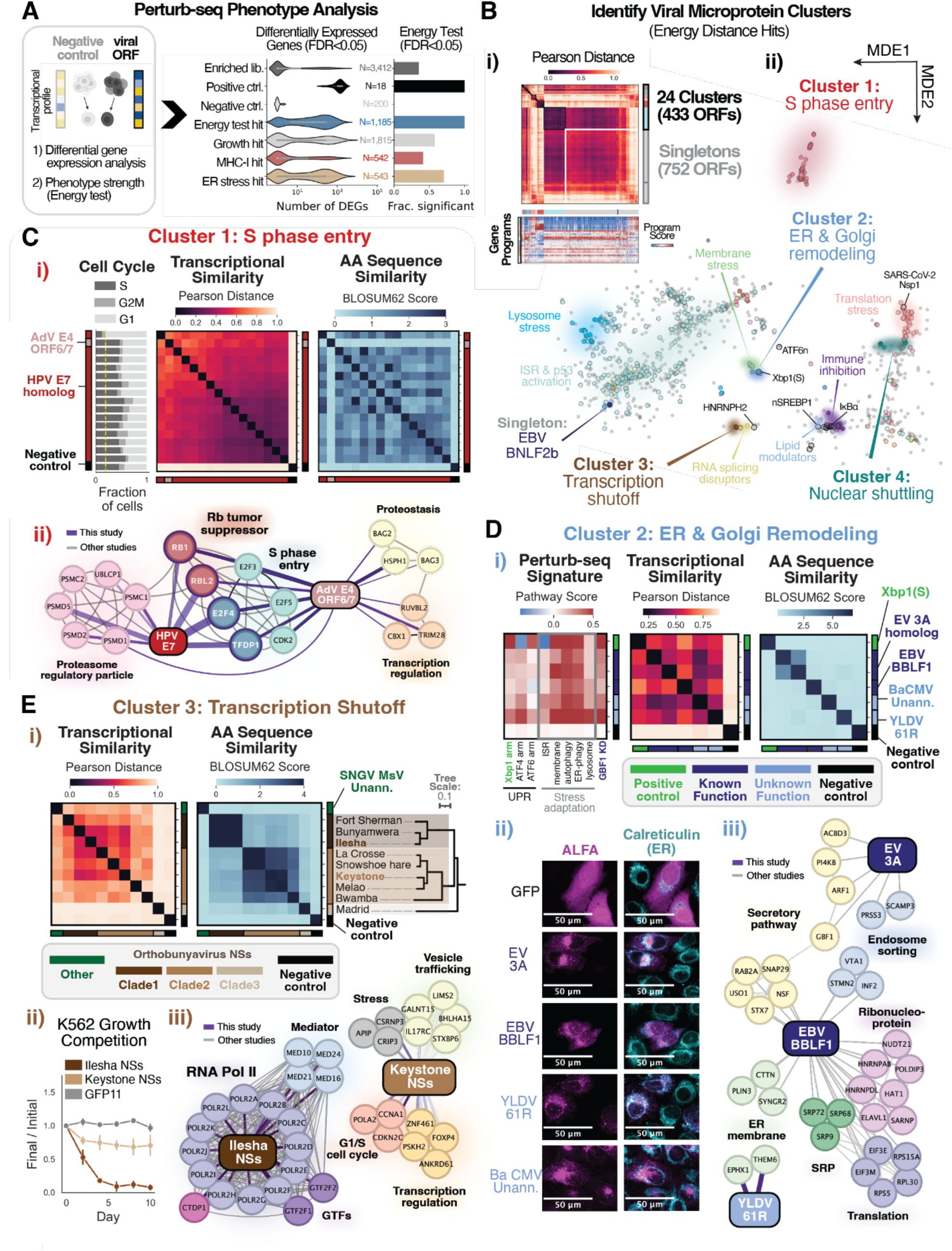
Multi-omic data integration reveals that viral small proteins have diverse cellular and molecular functions. (**A**) Perturb-seq phenotype strength evaluated by differential gene expression analysis and the energy test relative to negative controls. Violin plot of the number of differentially expressed genes induced by smORF overexpression, and the fraction of hits from each secondary screen that pass the energy test. (**B**) Identification of Perturb-seq clusters and singletons from all smORFs passing the energy test. (i) Pairwise Pearson distance matrix with a heatmap of gene-program scores encompassing 24 major biological processes. (ii) Minimal-distance embedding plot of pseudobulk smORF transcriptional profiles, colored by cluster assignment, with proposed functions annotated for a subset of clusters. (**C**) A cluster of smORFs that promote S-phase entry. (i) Cell-cycle fraction, transcriptional similarity, and global amino-acid sequence similarity. (ii) Host interactors determined by AE-MS in RPE-1 cells overexpressing the smORFs. (**D**) A cluster of smORFs involved in ER and Golgi remodeling. (i) Gene-expression pathway analysis, transcriptional similarity, and protein-sequence similarity matrix. (ii) Immunofluorescence confocal microscopy of ALFA-tagged elements co-stained with an anti-calreticulin antibody. (iii) Host interactors determined by AE-MS in 293F cells transfected with the smORF. (**E**) A cluster of homologous transcriptional-shutoff smORFs. (i) Transcriptional and protein-sequence similarity of the cluster, alongside the phylogenetic relationships of the orthobunyaviruses. (ii) K562 growth-competition experiments of cells transduced with Ilesha or Keystone virus NSs overexpression vectors. (iii) Host interactors determined by AE-MS in 293F cells transfected with the smORF. For all AE-MS networks, edges are colored as in Fig. 2.

To evaluate the accuracy of our method, we examined whether expected transcriptional signatures were captured in cells with positive-control ORF perturbations. At the single-cell level, cells clustered by ORF assignment rather than experimental batch (fig. S11A and B), and pathway-enrichment analysis of DEGs identified in ORF-overexpressing cells recovered the expected biological processes (fig. S11C). These benchmarks provided confidence in our new, more cost-effective Perturb-seq method, which we used to profile transcriptional phenotypes induced by 3,412 smORFs. We identified 1,185 viral smORFs that passed the energy test, 691 of which induced transcriptional phenotypes with >10 DEGs (Fig. 3A); of these, 75% were either unannotated (55%) or had little to no functional characterization (UniProt annotation score of 1).

### Perturb-seq classifies smORFs into functional clusters and singletons

Cellular phenotypes arise from coordinated changes across multiple host pathways rather than single genes. Perturb-seq captures this complexity with high resolution, but converting these high-dimensional profiles into biological meaning is nontrivial. As a first approach, we drew on prior CRISPRi Perturb-seq datasets in RPE-1 cells to curate gene sets and score 24 major gene programs important for RPE-1 function (*39*, *64*). Although positive-control perturbations showed the expected increases and decreases in this analysis (fig. S11D), this approach provided limited mechanistic insight in the absence of *a priori* hypotheses about the functions a smORF might encode (fig. S12).

To overcome this challenge, we leveraged a "guilt-by-association" annotation framework (*39*, *65*, *66*), which clusters genes based on a multidimensional feature space, enabling systematic inference of gene function by harnessing prior knowledge of well-characterized neighbors rather than transcriptional profiles alone. Specifically, we selected perturbations that passed the energy-distance test and clustered them based on two lower-dimensional representations: (i) a transcriptional-similarity matrix capturing the pairwise Pearson distance between all ORF pairs, and (ii) a low-dimensional space derived from a minimal-distance embedding of the transcriptional profiles (*39*). After clustering on each independently, we integrated the assignments and incorporated prior literature and gene-program scores to guide interpretation of viral smORF functions (Fig. 3B).

In total, our approach defined 24 clusters comprising 433 smORFs, alongside 752 singletons with phenotypes not closely shared with any other smORF in this dataset (Fig. 3B). Layering these results with protein-sequence analysis and viral phylogenies revealed three types of groupings: (i) clusters of homologous genes inducing similar transcriptional responses (Fig. 3C to D); (ii) clusters of non-homologous elements that independently converged on shared functions (Fig. 3C to D); and (iii) homologous sequences with diverging functions (Fig. 3E). By pairing transcriptional phenotypes and sequence homology, this organizational framework serves as a blueprint for exploring the function and evolution of viral microproteins.

### Perturb-seq cluster 1 (S-phase entry): convergent evolution of non-homologous viral proteins

One of the most distinctive clusters in our dataset consisted of viral microproteins that promote cell-cycle entry. This cluster was dominated by homologs of the well-characterized HPV E7 oncogene, but pairwise protein-sequence analysis revealed a non-homologous member, the adenovirus (AdV) E4 ORF6/7 microprotein. E4 ORF6/7 phenocopies the transcriptional changes from overexpressing E7, both driving the accumulation of cells in S and G2/M phases (Fig. 3C.i). Our AE-MS results confirmed this functional grouping: both microproteins physically associate with core members of the host Rb-E2F repressor complex (Fig. 3C.ii), a critical regulator of S-phase entry (*67*). This is consistent with the known functions of E7 and E4 ORF6/7, which target this complex through distinct mechanisms—E7 binds and degrades Rb (*49*), whereas E4 ORF6/7 stabilizes active E2F transcription factors (*68*). This case study demonstrates how our phenotype-first approach can link genes by shared phenotype, a connection that sequence- or structure-based homology pipelines would miss.

### Perturb-seq cluster 2 (Viral egress): guilt-by-association guides functional annotation of poorly characterized genes

Having established that our pipeline can group genes with shared functions, we asked whether it could infer functions for uncharacterized smORFs. We focused on a cluster containing the positive control XBP1(s), the master transcription factor of the IRE1 branch of the UPR (*69*). Alongside XBP1(s), this cluster includes several non-homologous viral elements: homologs of the enterovirus 3A ER-disruptor and MHC-I inhibitor, the EBV BBLF1 microprotein known to facilitate secretory transport of viral particles (*70*), and two poorly characterized elements: Yaba-like disease virus (YLDV) 61R and an unannotated baboon cytomegalovirus (BaCMV) smORF. Because 3A proteins activate the UPR to remodel the host ER-Golgi network and traffic viral replication organelles (*56*), we hypothesized that all cluster members act as ER/Golgi modulators. Consistent with this, they induced robust UPR activation (Fig. 3D.i), localized to the ER-Golgi (Fig. 3D.ii), scored as top hits in our ER-stress screens (fig. S8), and carried predicted TMDs. AE-MS further showed that YLDV 61R interacts with the ER-membrane components THEM6 and EPHX1 (Fig. 3D.iii). Together, these findings indicate that the cluster’s functional program centers on hijacking the host secretory pathway to support assembly of viral replication organelles and viral egress. This example demonstrates how our "guilt-by-association" framework can inform initial mechanistic hypotheses that specialized measurements can then confirm and refine.

### Perturb-seq cluster 3 (Transcription inhibition): divergent functions of homologous genes

Another notable cluster, marked by strong transcriptional responses, comprised highly toxic viral smORFs that ranked among the strongest hits in both K562 and RPE-1 growth screens (data S2). Cluster elements include multiple non-structural S-segment protein (NSs) homologs from negative-sense orthobunyaviruses, though some homologs diverged sharply. Madrid virus NSs, for instance, did not co-cluster in transcriptional space with the others (Fig. 3E.i). Divergence also extended to homologs that did co-cluster in transcriptional space, as illustrated by the differences in potency and host interactors of Ilesha and Keystone NSs. Ilesha NSs is markedly more toxic than Keystone NSs (Fig. 3E.ii) and interacts with host RNA polymerase (Fig. 3E.iii), consistent with the known function of the closely related La Crosse virus NSs as a transcriptional inhibitor (*71*). By contrast, Keystone NSs engaged cell-cycle factors and other transcriptional regulators (Fig. 3E.iii). This partitioning suggests that NSs homologs evolve rapidly, potentially under selection for functions tuned to distinct virus-specific pressures.

### Perturb-seq cluster 4 (Nuclear shuttling disruptors): independently evolved smORFs inhibit interferon signaling

With an analytical toolkit for mechanistic dissection in hand, we turned to a striking cluster of highly toxic viral smORFs grouped purely by transcriptional similarity (Fig. 4A and B), but drawn from phylogenetically distant viruses spanning multiple Baltimore classes. Its members were: 1) Vpr from the HIV-1 retrovirus (ssRNA-RT); 2) ORF6 from the SARS-CoV-2 coronavirus (ssRNA+); 3) MPXVgp154 homologs from the poxviruses (dsDNA) monkeypox (MPXV), cowpox (CPXV), and the more distant Yaba-like disease virus (YLDV) (fig. S13); and 4) an unannotated smORF from HMO astrovirus A (HMO AstV-A), a recently identified ssRNA+ virus primarily affecting immunocompromised individuals (*72*). Apart from the MPXVgp154 homologs, these microproteins shared little global sequence similarity (Fig. 4B), raising the question of what characteristics group them.

**Fig. 4.**
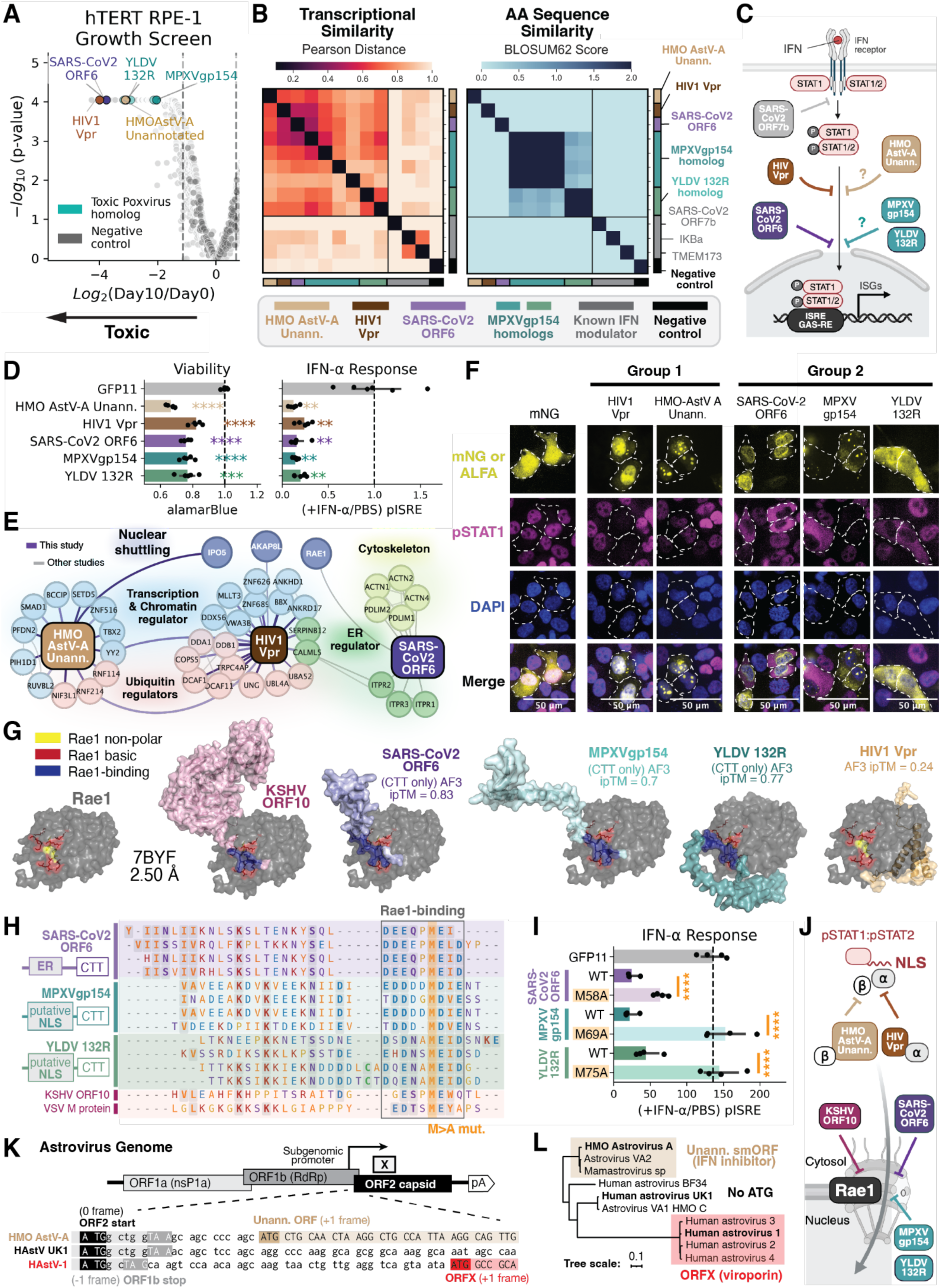
Convergence of smORFs that antagonize interferon signaling by disrupting nuclear shuttling. (**A**) Volcano plot of the 10-day growth screen in RPE-1 cells, with cluster elements highlighted. (**B**) Identification of a toxic cluster of homologous and non-homologous viral smORFs by Perturb-seq. (Left) Heatmap of the top differentially expressed genes for each smORF. (Middle) Transcriptional similarity matrix of pairwise Pearson distances between smORF-induced gene-expression profiles. (Right) Amino-acid sequence similarity matrix based on pairwise BLOSUM62 alignment scores. (**C**) Proposed mechanisms of IFN signaling inhibition. (**D**) Arrayed validation of cluster hits in transfected HEK293T cells. (Left) Cell viability measured by alamarBlue assay. (Right) Response to IFN-α (1,000 U/mL) measured using a pISRE-NanoLuc reporter assay. (**E**) AE-MS in 293F cells 28 h after transfection with C-terminally ALFA-tagged HMO AstV-A unannotated ORF, HIV-1 Vpr, and SARS-CoV-2 ORF6; interactors are integrated into the STRING host protein network. (**F**) Immunofluorescence confocal microscopy of A375 cells transfected with ALFA-tagged ORFs for 28 h and stimulated with IFNg for 45 min.(**G**) Structural comparison of Rae1 bound to KSHV ORF10 (PDB: 7BYF) and AF3 predictions of Rae1 in complex with indicated viral smORFs. (**H**) Identification of functional motifs. MSA of the putative Rae1-binding motif in the C-terminal tails of SARS-CoV-2 ORF6, MPXVgp154, and YLDV132R homologs, and of the putative NLS in HIV-1 Vpr, MPXVgp154, YLDV132R, and the HMO AstV-A unannotated ORF. (**I**) IFN-α response upon overexpression of Rae1-binding-motif mutants. (**J**) Proposed model for microprotein disruption of nuclear shuttling. (**K**) Astrovirus genomic locus encoding the unannotated smORF. (**L**) Phylogenetic tree of human astroviruses. All pISRE-NanoLuc data are shown as mean ± SEM; *P < 0.05, **P < 0.01, ***P < 0.001, ****P < 0.0001 (Student’s t-test relative to GFP11 control).

Examination of the top DEGs common to all members pointed to pathways consistent with its toxic impacts including p53-pathway activation, G1/S arrest, and apoptosis (fig. S14A). However, since toxicity and activation of these pathways can arise through many mechanisms, we turned to prior literature on well-characterized cluster members to narrow down potential functions. Although Vpr and ORF6 are not typically regarded as functional analogs, both have been shown to inhibit interferon signaling (*73*, *74*) (Fig. 4C), prompting us to test whether IFN-α/γ antagonism was shared across the cluster. Indeed, in arrayed luciferase reporter assays, overexpression of any cluster element significantly decreased response to IFN-α and IFN-γ alongside impaired cell viability (Fig. 4D, fig. S14B).

Applying the same logic we used to dissect the MHC-I inhibitors, we next sought to understand which step of the interferon signaling pathway is disrupted by this microprotein cluster. The known mechanisms by which Vpr and ORF6 inhibit the JAK-STAT pathway point to disruption of nucleocytoplasmic transport, through distinct routes: Vpr competes with host cargo for nuclear-import receptors (*73*), whereas ORF6 binds the nuclear-pore component Rae1 to block pSTAT1 translocation (*75*). We therefore hypothesized that hijacking of nucleocytoplasmic transport is the molecular signature defining this cluster (Fig. 4C).

To test this, we used three complementary assays: transcriptional-phenotype analysis, AE-MS, and microscopy. First, we reasoned that if cluster elements specifically disrupt nuclear shuttling, they should not co-cluster with interferon inhibitors that act through other mechanisms. Indeed, modulators such as SARS-CoV-2 ORF7b, which blocks interferon activation at the STAT1-phosphorylation step (*76*), showed transcriptional profiles distinct from cluster elements (Fig. 4B). The cellular localization and physical interactomes of ALFA-tagged cluster elements further supported our model, revealing two groups matching the known functions of Vpr and ORF6. The first comprises Vpr and an unannotated HMO AstV-A smORF, both of which interact with nuclear-import receptors: Vpr is known to interact with importin-α (*73*) and the unannotated HMO AstV-A smORF was identified to interact with the importin-β transport protein IPO5 in our AE-MS dataset (Fig. 4E, data S7). Consistent with this, both show nuclear or nucleolar localization (Fig. 4F) and contain basic residues that may be nuclear localization signals (NLSs) (fig. S15). The second group, SARS-CoV-2 ORF6, MPXVgp154, and YLDV132R, blocks nuclear import, shown by nuclear exclusion of pSTAT1 in IFN-γ-stimulated A375 cells expressing these microproteins (Fig. 4F).

These findings indicate that disruption of nucleocytoplasmic transport, an essential host process, is the defining function of this cluster, with the toxicity and impaired interferon response likely arising as indirect consequences.

### Identification of a shared Cluster 4 molecular key: independent emergence of a SLiM that mediates IFN inhibition

Having identified nuclear-shuttling disruption as a shared function of this cluster, we asked whether a common molecular key underlies the specialized phenotypes induced by SARS-CoV-2 ORF6, MPXVgp154, and YLDV132R. We again leveraged AF3 structure prediction to nominate functionally important residues for further analysis. Because the ORF6-Rae1 interaction is critical to the ability of ORF6 to block nuclear import (*75*), we examined predicted structures of Rae1 bound to each microprotein. These highlighted residues in the C-terminal tails (CTTs) of all three that contact the cargo-binding residues of Rae1 (Fig. 4G). For ORF6, the predicted interface matched residues previously shown to be important for its function (*75*). These structures also resemble the solved structure of Rae1 bound to KSHV ORF10, a known viral inhibitor of mRNA nucleocytoplasmic transport (PDB 7BYF) (*77*). By contrast, AF3 predictions of Rae1 with full-length and truncated versions of HIV-1 Vpr lacked a comparable interface and had poor confidence scores (ipTM < 0.4) (Fig. 4G).

Guided by these residues, we constructed a multiple-sequence alignment (MSA) of the CTTs of SARS-CoV-2 ORF6 and MPXVgp154 homologs alongside the N-terminal Rae1-binding region of KSHV ORF10, which revealed a shared DDXXMEID-like short linear motif (SLiM) across these sequences (Fig. 4H). To test the importance of this motif for MPXVgp154 and YLDV132R function, we mutated the critical methionine to alanine, a substitution shown to abolish nucleocytoplasmic-transport inhibition by KSHV ORF10 and SARS-CoV-2 ORF6 (*75*). Each mutant (ORF6 M58A, MPXVgp154 M69A, and YLDV132R M75A) lost the IFN-inhibition phenotype relative to wild type (Fig. 4I; fig. S14C), arguing that the independent emergence of SLiMs predicted to bind Rae1 underlies the shared function of SARS-CoV-2 ORF6, MPXVgp154, and YLDV132R.

This vignette illustrates how pairing Perturb-seq with decades of accumulated knowledge on well-studied genes can convert complex phenotypes into specific, testable mechanisms. Furthermore, it uncovers a case of convergence at the pathway and molecular level, illustrated by independent arrival to the same short sequence motif (Fig. 4H) or related host nodes (Fig. 4J). This cluster has intriguing parallels with prior observations of MHC-I presentation modulators: in both, multiple viral inhibitors from across the virome converge on a single host node, namely Rae1 here and TAP there, which may mark them as particularly vulnerable points in the host protein network.

### De novo emergence of an unannotated smORF from HMO Astrovirus via overprinting

With the mechanistic function of this cluster resolved, we next explored the evolution of its members, which drew our attention to the unannotated HMO astrovirus A smORF. Despite its potent toxicity and ability to inhibit interferon signaling (Fig. 4A to D), it had escaped conventional annotation, prompting us to ask what sequence features allowed it to go undetected. A BLAST search of its protein sequence returned no significant hits against the NCBI database, leading us to turn to its genomic origins.

The HMO AstV-A genome is compact and information-dense, spanning ∼7 kb across three overlapping ORFs (ORF1a, ORF1b, and ORF2) (Fig. 4K). ORF1a and ORF1b each encode a polyprotein and overlap one another, with ORF1b translated via programmed -1 ribosomal frameshifting. Downstream of and overlapping with ORF1b is ORF2, which encodes the viral capsid protein and is translated from a highly expressed subgenomic transcript bearing a short (∼20-bp) 5′ untranslated region (UTR) (*78*). The unannotated smORF is embedded entirely within ORF2 but read in the +1 frame, making it an overprinted gene, which is a protein-coding sequence that overlaps another functional gene but is read in a different frame (*79*). Its start codon is just 19 bp downstream of the ORF2 initiation site, a configuration resembling that of the SARS-CoV-2 genes ORF9b and ORF3b. Phylogenetic analysis showed that the smORF is lineage-restricted, with its start codon absent from other closely related astroviruses, pointing to a *de novo* evolutionary origin (Fig. 4K and L; fig. S16).

Unexpectedly, further analysis of more distantly related astrovirus lineages revealed additional smORFs arising from the same genomic locus, but initiating from different start codons (Fig. 4K; fig. S16). In agreement, an independent study based on conservation analysis suggested that there have been multiple parallel occurrences of the *de novo* emergence of smORFs within the +1 frame of the 5′ region of the ORF2 gene. That study characterized one such smORF, termed ORFX, in the Human Astrovirus 1 (HAstV1) clade, showing that it is highly translated during active infection, contains a predicted TMD, localizes to the host plasma membrane, and has viroporin-like activity (*78*). By contrast, the unannotated HMO AstV-A smORF identified here has a distinct sequence and function: a putative NLS (fig. S15), nucleolar localization (Fig. 4F), interaction with nuclear import receptors (Fig. 4E), and interferon-blocking activity (Fig. 4D). The independent emergence of structurally and functionally disparate microproteins from the same locus across distinct astrovirus clades demonstrates that the astrovirus genome harbors "hot spots" supporting *de novo* gene birth.

### EBV BNLF2b is a singleton that promotes epithelial cell proliferation

Although our guilt-by-association approach successfully predicted functions for clustered elements, singletons require a different strategy. We therefore looked to the strongest hits in our genetic screens and prioritized smORFs that accelerate growth, a phenotype relevant to virus-associated malignancies.

Growth-promoting smORFs were detected almost exclusively in immortalized RPE-1 cells and did not score in cancer-derived K562 cells (Fig. 5A; fig. S17A), underscoring the importance of cellular context for studying oncogenic phenotypes (*80*). Beyond the aforementioned HPV E7 oncoprotein, six other viral microproteins validated in orthogonal arrayed growth-competition assays (fig. S17C and D). All six are singletons in the Perturb-seq dataset, sharing no transcriptional phenotype, amino-acid sequence similarity, or host-protein interactors with E7 or with one another (Fig. 5B; fig. S18, S19).

**Fig. 5.**
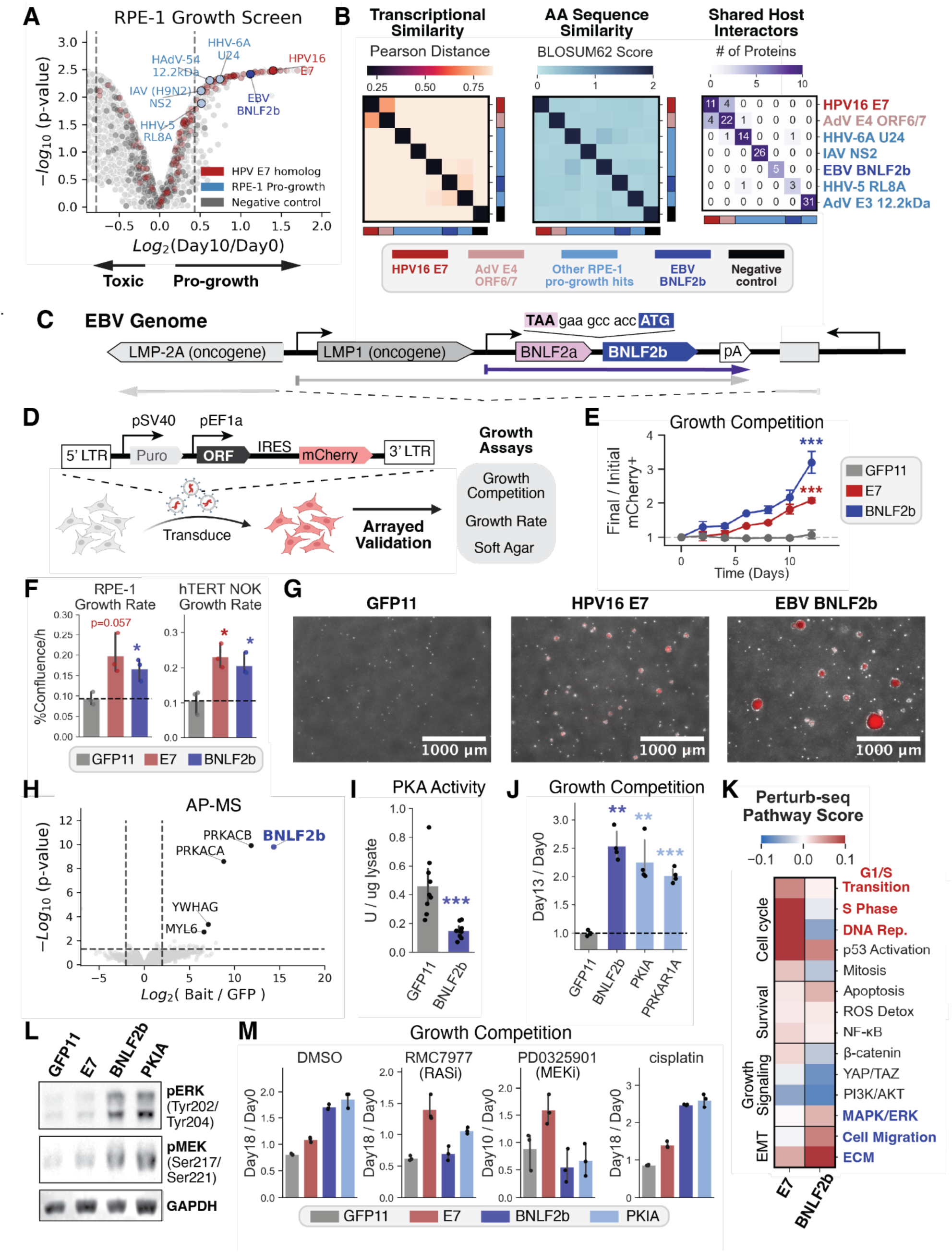
EBV BNLF2b is an oncogene that drives growth by inhibiting PKA to activate MAPK signaling. (**A**) Volcano plot of 10-day RPE-1 growth screens with pro-growth hits and HPV E7 homologs highlighted. (**B**) Pairwise transcriptional and protein sequence similarity as well as shared host interactor among pro-growth hits and HPV16 E7 and HAdV E4 ORF6/7. (**C**) EBV genome locus encoding BNLF2b. BNLF2b is located in a polycistronic transcript encoding BNLF2a upstream of BNLF2b. The neighboring genes LMP1 and LMP2A are known EBV oncogenes. (**D**) Experimental design of orthogonal growth assays. Viral smORFs were overexpressed in RPE-1 cells using a constitutive lentiviral vector with an IRES-mCherry on the same transcript as the ORF. (**E**) Growth competition assays tracking mCherry+ population over time. (**F**) Proliferation kinetics of puromycin-selected cells, measured by longitudinal phase-contrast confluence imaging in RPE-1 (left) and hTERT NOKs (right). (**G**) Soft agar colony formation assay 30 days post-seeding for assessment of anchorage-independent growth. Images show phase contrast overlaid with mCherry fluorescence. (**H**) Volcano plot of AE-MS for ALFA-tagged BNLF2b overexpressed in RPE-1 cells. (**I**) Biochemical PKA activity assay in RPE-1 lysates expressing GFP11 (control) or BNLF2b. (**J**) Growth competition assays in RPE-1 cells overexpressing the endogenous PKA inhibitors PKIA or PRKAR1A. (**K**) Transcriptional scoring of growth and survival pathways from RPE-1 Perturb-seq data. Pseudobulk gene expression profiles for HPV E7 and EBV BNLF2b were scored based on established oncogenic gene sets. (**L**) Western blot analysis of phosphorylated ERK (Thr202/Tyr204) and MEK (Ser217/Ser221) levels. (**M)** Growth competition of cells overexpressing HPV E7, BNLF2b, or PKIA treated with the RAS inhibitor RMC-7977 (1uM), the MEK inhibitor PD0325901 (1uM), or cisplatin (100nM). Data represent mean +/- SEM; *p < 0.05, **p < 0.01, ***p < 0.001, ****p < 0.0001 (Student’s t-test relative to GFP11 control).

Among the validated pro-growth smORFs, EBV BNLF2b emerged as an intriguing candidate. First, it was one of the most potent growth-promoting hits in both the primary screens and the arrayed growth-validation assays (Fig. 5A; fig. S17C and D). Second, it lies within an EBV genomic locus (Fig. 5C) that includes the well-established oncogenes LMP1 (*81*) and LMP2A (*82*), suggesting a coordinated functional role given the evolutionary tendency of viral genomes to group related genes. Third, although ribosome-profiling studies confirm its active translation (*10*), BNLF2b remains poorly characterized, with a UniProt annotation score of 2 (*44*). Although BNLF2b was included in large-scale affinity-purification mass spectrometry surveys of EBV proteins, its function or pro-growth phenotype had not been documented (*83*, *84*). Finally, BNLF2b is linked to EBV-associated epithelial malignancies. It is highly transcribed in EBV-positive epithelial tumors (*85*, *86*), and a recent landmark study identified serological anti- BNLF2b antibodies (P85-Ab) as a sensitive biomarker for mass screening of nasopharyngeal carcinoma (NPC), but not for infectious mononucleosis or EBV-associated lymphomas (*41*).

To further characterize the growth-promoting capabilities of BNLF2b, we performed three orthogonal assays in RPE-1 with the pConstitutive expression vector (Fig. 5D). In both competition assays (Fig. 5E) and confluency-based growth rate measurements (Fig. 5F), cells expressing BNLF2b or E7 exhibited a robust fitness advantage relative to GFP11 controls. This pro-proliferative phenotype was also validated via growth rate measurements in normal oral keratinocytes (NOKs) (Fig. 5F), a cell-line model commonly used to study viral oncogenes (*87*). Most remarkably, both BNLF2b and E7 promoted robust anchorage-independent growth in soft agar (Fig. 5G), a key signature of oncogenic transformation (*88*) that was not observed for any other tested pro-growth candidate. These findings motivated us to investigate the molecular mechanism of BNLF2b-driven growth.

### EBV BNLF2b inhibits PKA to drive proliferation via downstream MAPK activation

Because BNLF2b does not co-cluster with E7 or any other viral smORFs (Fig. 5B), we hypothesized that it operates through a pro-growth pathway distinct from Rb inhibition. AE-MS profiling of ALFA-tagged BNLF2b in RPE-1 cells identified PRKACA and PRKACB, the catalytic subunits of protein kinase A (PKA), as top interactors (Fig. 5H). Western blot analysis validated these interactions and confirmed that BNLF2b does not bind the regulatory subunits of the PKA holoenzyme (fig. S20A and B). To evaluate how this interaction impacts PKA function, we performed PKA activity assays and Western blotting for phosphorylated PKA substrate levels in RPE-1 lysates, which revealed that BNLF2b acts as an inhibitor (Fig. 5I; fig. S20B).

Furthermore, the native human PKA inhibitor PKIA phenocopied BNLF2b when ectopically expressed in RPE-1 growth-competition assays (Fig. 5J), demonstrating that PKA inhibition alone is sufficient to promote epithelial cell fitness.

How the biochemical function of BNLF2b translates to enhanced growth was not immediately obvious, as PKA regulates diverse cellular processes and can promote or suppress growth depending on context (*89–93*). To reconcile PKA inhibition with increased proliferation, we analyzed our Perturb-seq data and calculated pathway scores for key growth-promoting processes. Unlike E7, which directly activates cell-cycle programs, BNLF2b induced a transcriptomic state marked by MAPK/ERK signaling, cell migration, and extracellular-matrix (ECM) remodeling (Fig. 5K). Western blotting validated MAPK activation, showing increased pERK and pMEK in cells overexpressing BNLF2b or PKIA, but not E7 (Fig. 5L; fig. S20C).

Having established that BNLF2b activates MAPK signaling, we next asked whether this pathway is necessary for BNLF2b-driven growth and performed growth-competition experiments under pharmacological perturbation of the MAPK cascade. Treatment with the RAS inhibitor (RASi) RMC-7977 or the MEK inhibitor (MEKi) PD0325901 selectively abolished the fitness advantage conferred by BNLF2b or PKIA, while leaving E7-driven growth unaffected (Fig. 5M). These targeted inhibitors were more effective at abrogating BNLF2b-mediated growth than cisplatin, a current clinical standard of care for Stage II and above NPC tumors (*94–96*). Conversely, pharmacologically activating PKA with forskolin and IBMX reduced overall cell growth and amplified the fitness disparity between wild-type and BNLF2b-expressing cells (fig. S20D). These findings demonstrate that BNLF2b drives epithelial cell proliferation by co-opting the PKA–MAPK signaling axis.

### Identification of an EBV BNLF2b molecular key: mimicry of a human PKA inhibitor motif to promote growth

To resolve the molecular basis of the BNLF2b–PKA interaction, we modeled the complex using AF3. Truncating disordered regions of BNLF2b (residues 35–61) yielded a confident prediction (ipTM = 0.79) in which BNLF2b occupies the substrate-binding cleft of PKA. Structural alignment with a reference complex of PKA bound to an inhibitor peptide (PDB: 3FJQ) (*97*) revealed that BNLF2b is predicted to bind PKA through molecular mimicry of human PKA inhibitors such as PKIA and PRKAR1A (Fig. 6A).

**Fig. 6.**
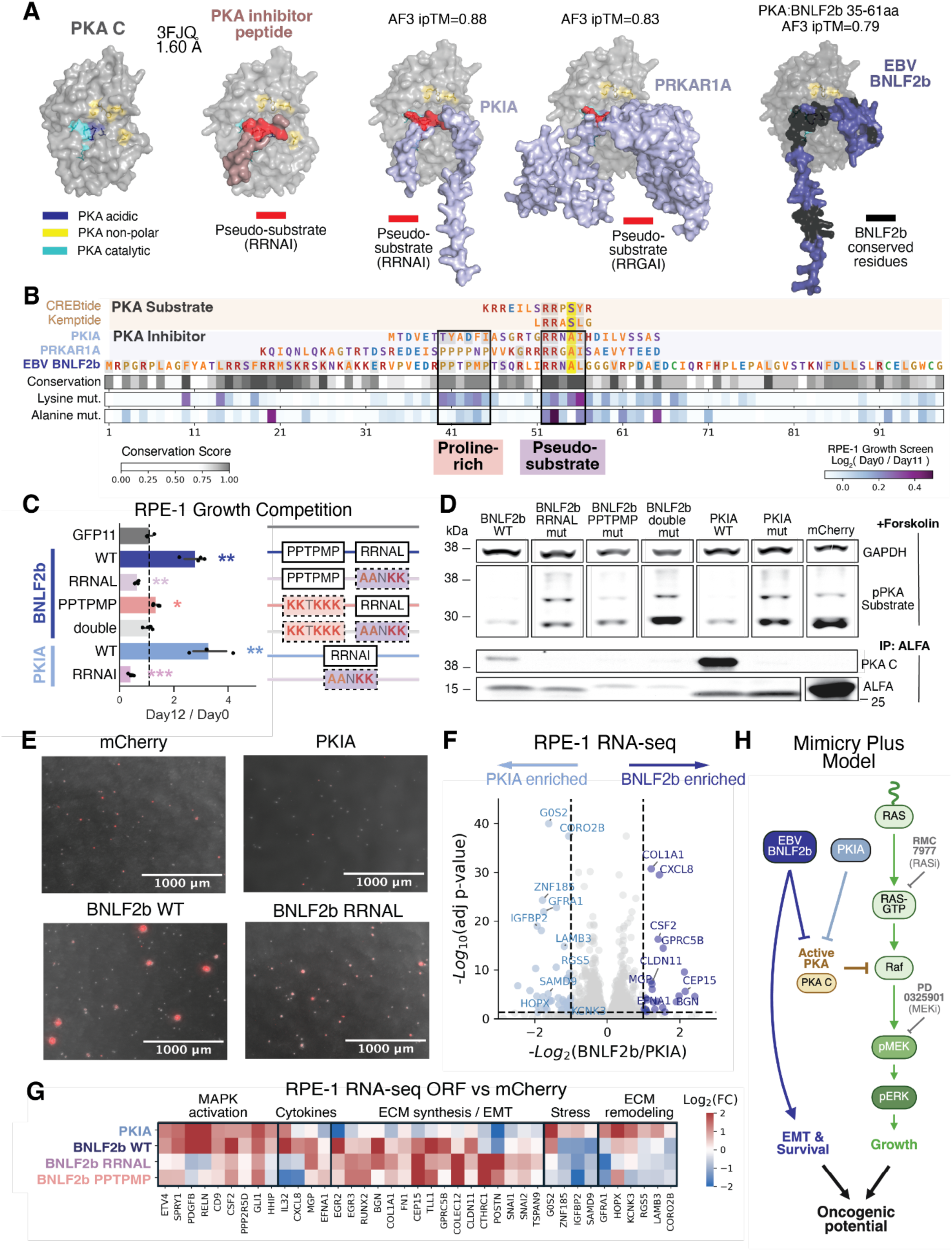
BNLF2b mimics host PKA inhibitors while driving divergent oncogenic programs. (**A**) Structural comparison of the PKA catalytic subunit co-crystallized with a canonical inhibitor peptide (PDB: 3FJQ) and AF3 predicted complexes with PRKAR1A, PKIA, or BNLF2b. (**B**) Mapping of BNLF2b functional motifs. (Top) MSA with host PKA substrates and inhibitors reveals a conserved proline-rich motif and a putative pseudosubstrate motif. (Bottom) High-resolution alanine and lysine scanning of BNLF2b for loss of pro-growth phenotype. Epitope mapping of anti-BNLF2b peptide reactivity from serum antibodies from NPC patients. (**C**) Functional validation of identified motifs via growth competition assays. (**D**) Biochemical characterization of PKA inhibition. Western blot analysis of total phospho-PKA substrate levels and corresponding anti-ALFA immunoprecipitations (IP) assessing bait stability and PKA recruitment. (**E**) Anchorage-independent growth. Soft agar colony formation assays shown by representative phase-contrast microscopy 40 days post-seeding (top) and quantification of colony size (bottom). (**F**) Comparative transcriptomics. RNA-seq analysis comparing DEGs between BNLF2b and PKIA overexpressing cells. (**G**) Heatmap of top DEGs grouped by pathway. (**H**) Proposed “mimicry plus” model for BNLF2b driven proliferation in epithelial cells. BNLF2b and PKIA inhibit the high basal PKA activity, resulting in activation of MAPK signaling. Simultaneously, BNLF2b activates additional EMT and survival programs that contribute to its oncogenic potential.

Sequence analysis and MSAs further supported this molecular-mimicry model. BNLF2b contains a conserved RRNAL motif that mirrors the RRXAI pseudosubstrate motifs of PKIA and PRKAR1A, in which the phosphorylatable serine/threonine of a typical PKA substrate is replaced by a non-phosphorylatable alanine. BNLF2b also has a conserved proline-rich motif adjacent to this pseudosubstrate sequence that resembles an auxiliary regulatory motif in PRKAR1A (Fig. 6B).

To validate the functional importance of these two motifs, we performed pooled growth screens using a library of alanine- and lysine-scanning BNLF2b variants, which showed that mutations within either the RRNAL pseudosubstrate motif or the proline-rich sequence decreased BNLF2b-driven growth (Fig. 6B; fig. S21). To validate these screening results, we designed targeted mutants disrupting each motif individually or in combination for further characterization. Indeed, these mutants failed to confer a fitness advantage (Fig. 6C) and exhibited impaired PKA binding and inhibition (Fig. 6D; fig. S22A and B). These results define two essential mimicry motifs in BNLF2b. On a methodological side, this approach illustrates how the compact size of microproteins enables rapid, systematic identification of critical residues.

### EBV BNLF2b activates an additional oncogenic program independent of PKA inhibition

Although endogenous PKA inhibitors, including PKIA, are associated with poor prognosis in certain clinical contexts (*92*, *93*), they are not classically regarded as autonomous oncogenes. This distinction led us to investigate whether BNLF2b induces phenotypes not explained by PKA inhibition alone. Soft-agar assays revealed a striking divergence between BNLF2b and PKIA: whereas BNLF2b robustly drives anchorage-independent growth of RPE-1 cells, PKIA fails to do so (Fig. 6E). Furthermore, PKA-inhibition-defective BNLF2b mutants retained partial capacity to induce soft-agar colony formation (fig. S22C and D), suggesting that BNLF2b harbors pro-growth capabilities independent of its function as a PKA inhibitor.

To define the molecular basis of these phenotypic differences, we performed RNA-seq in RPE-1 cells expressing PKIA, wild-type BNLF2b, PKA-inhibition-defective BNLF2b mutants, or mCherry control. Differential gene expression analysis revealed three distinct transcriptional signatures (Fig. 6F and G). First, supporting that PKA inhibition alone is sufficient to activate MAPK signaling, downstream MAPK targets *ETV4* and *SPRY1* were upregulated by PKIA and wild-type BNLF2b, but to a lesser extent by PKA-inhibition-defective BNLF2b mutants. Second, to identify pathways responsible for the BNLF2b-driven anchorage-independent growth, we examined DEGs shared by BNLF2b variants but absent in PKIA. This analysis revealed upregulation of genes involved in ECM synthesis and epithelial-to-mesenchymal transition (EMT), including *RUNX2*, *COL1A1*, *FN1*, *BGN*, and *CLDN11*. Third, PKIA overexpression upregulated stress-induced and anoikis-related genes (*G0S2*, *ZNF185*, *IGFBP2*, *SMAD9*) that were downregulated by wild-type and BNLF2b mutants (Fig. 6G). Together, these findings support a two-arm model in which BNLF2b enhances oncogenic potential through PKA-inhibitor mimicry that activates MAPK signaling, alongside PKA-independent induction of EMT and suppression of anoikis (Fig. 6H).

Overall, mechanistic dissection of BNLF2b illustrates how high-throughput, unbiased genetic screens can complement targeted biochemical assays to identify and annotate poorly understood microproteins. Pooled phenotypic screening provided an entry point to isolate biologically active smORFs from a library of tens of thousands, while high-resolution multi-omic assays resolved their specific activities and functions. Scaling this multilayered framework across the virome and dark proteome is a tractable, powerful route to systematically decoding host–pathogen interfaces.

## Discussion

Here, we present a pan-viral functional atlas that maps the diversity of viral microproteins, revealing the broader principles underlying their activities. Using a multi-assay funnel strategy, we scaled from a 52,979-element smORF library to high-resolution phenotypic profiling of an enriched 4,107-element subset, identifying more than 2,000 biologically active smORF-encoded microproteins from over 300 viruses spanning all Baltimore classifications. This profiling showed that viral microproteins target a wide range of essential host processes, including the cell cycle, stress responses, transcription, nuclear transport, innate and adaptive immunity, protein degradation, and ER processing, among others. These high-throughput assays nominated robust candidates for further investigation, which led to the identification and mechanistic dissection of the oncogenic properties of a previously uncharacterized EBV microprotein. Collectively, our findings allow us to explore three themes: (i) viral microproteins adopt diverse, often multifunctional activities at the cellular, molecular, and biochemical levels; (ii) convergence on shared functions is pervasive across the virome; and (iii) viral microproteins represent a rich reservoir of evolutionary innovation.

In reference to the first theme, despite their small size, viral microproteins occupy an expansive functional landscape arising from diversity at multiple scales. At the cellular level, they modulate a wide variety of host processes, ranging from cell proliferation (N = 2,028), antigen presentation (N = 748), survival under ER stress (N = 602), to global transcriptional state (N = 1,185), with many exhibiting potencies comparable to canonical proteins. This breadth of cellular phenotypes is mirrored by an expansive arsenal of molecular strategies for perturbing the host. Antigen presentation provides a striking example: viral inhibitors act at nearly every step of the pathway, from synthesis (AKMV OPG024) and peptide-loading complex assembly (the TAP inhibitors ICP47, BNLF2a, and CPXV012) to MHC-I maturation (HPV E5 and UL147A), trafficking (HPV E5 and CeHV US5), and degradation (STL polyomavirus). Likewise, the 24 functional programs identified by Perturb-seq coupled with AE-MS show that similar cellular outcomes can be achieved through distinct molecular mechanisms. For example, ER remodeling can arise from perturbing different ER-trafficking and membrane regulators, and nucleocytoplasmic transport can be disrupted by targeting either nuclear import receptors or nuclear pore components. Thus, viral microproteins can engage a broad spectrum of host proteins and pathways.

Underlying this broad array of molecular activities is a versatile biochemical toolkit of molecular keys serving as interaction interfaces for reprogramming host protein networks. In place of large scaffolds and multidomain architectures, microproteins utilize molecular keys that fall into two general classes: 1) longer structured elements (∼20aa), including transmembrane helices, beta-hairpins, and signal peptides and 2) SLiMs (∼5aa), which range from common sequences such as NLSs to more specialized motifs (e.g., Rae1-interacting and PKA pseudosubstrate motifs) (Fig. 4H, 6B). These components can be combined to enhance potency or confer additional functions. For example, MHC-I inhibitors pair TMDs with TAP-binding regions (BNLF2a and CPXV012) or ER-targeting sequences (CeHV UL147A and CeHV US5) to ensure proper localization that strengthens their inhibitory activity. Similarly, SARS-CoV-2 ORF6-like proteins combine Rae1-binding motifs with subcellular-localization domains that concentrate them at nuclear pores to enhance blockade of nucleocytoplasmic transport (*75*, *98*). Most notably, EBV BNLF2b mimics the pseudosubstrate motif of human PKA inhibitors to promote proliferation, while inducing additional oncogenic phenotypes independent of its PKA inhibitory residues (Fig. 6H). Together, these examples suggest that microproteins are composed of modular molecular keys that are combinatorially assembled to yield diverse, potent, and multifunctional activities.

Regarding the second theme, convergence on shared functions is widespread even among phylogenetically distant viruses. Although seemingly paradoxical given the broad diversity of viruses, this reflects the selection pressures driving viruses to repeatedly counteract host processes. This is well-illustrated at the cellular level, as hundreds of microproteins were found to impact growth, signaling, and stress responses. The MHC-I pathway illustrates this particularly well: recurrent antagonism by large dsDNA viruses such as herpesviruses (HSV ICP47, EBV BNLF2a, CeHV UL147A), poxviruses (CPXV012, AKMV OPG024), papillomaviruses (HPV E5), and polyomaviruses (STLPyV unannotated smORF), underscores that T-cell immunity is a major barrier to these viruses, which often establish persistent infections and carry large proteomes that generate immunogenic peptides (*50*). At the molecular and biochemical level, we observe multiple examples of convergence toward the same host nodes: TAP inhibition to inhibit antigen presentation (EBV BNLF2a, HSV ICP47, CPXV012) (*53*), Rb repressor complex (HPV E7 and HAdV E4 ORF6/7) inhibition to drive S-phase entry, and nuclear-import receptor targeting to suppress immune responses (HIV-1 Vpr, HMO AstV-A smORF) (*73*). Perhaps most surprising is convergence at the biochemical level. Despite the vast sequence space available to viral microproteins, two cases of molecular mimicry were identified: Rae1-binding SLiMs shared across poxviruses (MPXVgp154, YLDV132R, VACV protein 169), sarbecovirus (ORF6), and herpesvirus (KSHV ORF10) (Fig. 4H) and PKA pseudosubstrate motifs shared between human PKIA and EBV BNLF2b (Fig. 6B). These recurrent solutions mark these host nodes as especially effective points for viral hijacking, suggesting that systematically annotating the viral proteome can clarify the selection pressures different viruses face and, in turn, reveal key host vulnerabilities.

In reference to the third theme, pervasive convergence supports a model in which viruses continually generate new sequences that are selected for functions, positioning microproteins as an important reservoir of evolutionary innovation. We capture a case of the *de novo* gene birth of an unannotated HMO AstV-A smORF that hijacks host nuclear-transport machinery and arose through overprinting. Remarkably, the same genomic locus in other astrovirus clades has independently evolved smORFs that initiate from distinct start codons, have poor sequence similarity, and have entirely different functions. This repeated emergence of new smORFs marks this locus as a “gene nursery”, a site that supports new gene production (*99*). For this locus, features including high transcript-level expression, a short UTR enabling polycistronic translation via leaky scanning, and flexible sequence constraints enable *de novo* emergence (*78*). These characteristics have been found in SARS-CoV-2 genomic loci containing overprinted genes (*100*), raising questions about the prevalence, characteristics, and origins of gene nurseries across the virome. Beyond generating new functional peptides, viruses face strong, shifting selection pressures that drive rapid functional divergence. For example, the orthobunyavirus NSs homologs differ in potency and host interactions (Ilesha vs. Keystone NSs), with Madrid virus NSs the most distinct, clustering separately from the other homologs in the Perturb-seq dataset (Fig. 3E). These considerations support a model for how viral smORFs emerge and evolve complex functions: (i) random generation of new sequences; (ii) emergence of short molecular keys; (iii) selection of elements to more robustly impact the host protein network; and, ultimately, (iv) broad expansion and conservation (fig. S23).

Beyond advancing fundamental concepts in virus-host biology and gene evolution, characterizing the viral microproteome has therapeutic implications. This is illustrated by our mechanistic dissection of the oncogenic capabilities of EBV BNLF2b, which is particularly intriguing given its connection to the EBV-associated malignancy NPC. Although EBV infects over 90% of the global population, it causes certain lymphomas and epithelial malignancies in a geographically restricted manner. Recent clinical studies evaluating biomarkers to distinguish healthy EBV-positive individuals from NPC patients identified high anti-BNLF2b antibody titers as a robust diagnostic marker, though BNLF2b’s biological function was unknown (*41*). Our study offers a mechanistic explanation for this clinical observation: BNLF2b expression induces oncogenic properties in epithelial cells through a “mimicry-plus” strategy (Fig. 6H). BNLF2b promotes proliferation by activating MAPK signaling through PKA inhibition, while engaging additional oncogenic programs that enable anchorage-independent growth, a phenotype not observed from overexpressing the selective PKA inhibitor PKIA. Future efforts will be needed to explore its role in NPC, but our findings provide a rationale for exploring MAPK-targeted therapies in NPC and related EBV-associated epithelial malignancies.

Several considerations suggest our studies underestimate the full scope of the viral microproteome, both in number and functional diversity. First, our approach misses microproteins that require other viral co-factors. For example, the well-annotated EBV BDLF3.5 gene did not score in any screen because it acts within the viral pre-initiation complex (vPIC) to drive late lytic gene expression (data S2 and S3). Such co-dependencies could be reconciled using CRISPR-based screens or deep mutational scanning of intact viral genomes (*48*, *101*, *102*). Second, many functions are highly context-specific, so our efforts capture only a subset of true physiology. This limitation is particularly acute for oncogene identification, since pro-growth phenotypes depend heavily on cell type and media conditions (*80*, *103*). Furthermore, reliance on transformed or cancer-derived cell lines can mask the effects of exogenous oncogenic perturbations. Although BNLF2b is the first reported viral oncogene modulating the PKA-MAPK axis, others may act through the same mechanism but have been overlooked due to biases of experimental models. Finally, our funnel’s initial growth-phenotype criterion misses smORFs with growth-independent functions, as the MHC-I screens illustrate. Screens covering a broader range of phenotypes, physiological contexts, and cell types could serve as alternative entry points to the funnel, as used in other recent virome studies (*36–38*). Ultimately, resolving the complete viral landscape will require complementary strategies, encompassing host-centered overexpression alongside direct viral mutagenesis as well as broad unbiased screening paired with specialized reporters.

As obligate parasites, viruses have evolved proteomes deeply integrated with host protein networks. Systems-level profiling of these interfaces can offer valuable insight into both virology and cell biology. Enabled by the scalability of Perturb-seq (*52*, *104*), our approach could extend to the entire pan-viral proteome for function-based organization of viral classes. Such a resource could expose host vulnerabilities to inform pan-viral drug design for pandemic preparedness (*105*). Furthermore, because viruses have evolved to manipulate host machinery in profound ways, viral ORFs, both canonical and non-canonical, offer a class of perturbations distinct from widely used CRISPR-based tools for probing host biology. By driving cells into novel states, viral-ORF perturbations may reveal hidden processes informing both targeted discovery and virtual-cell modeling efforts. Ultimately, this work contributes toward a future of predictive virology (*106*) and cell biology, capable of anticipating the functional consequences of newly evolved viral sequences and novel cellular phenotypes.

## Acknowledgments

We thank members of the Weissman, Mann, Gewurz, and Elledge labs for helpful discussions and advice. From the Weissman Lab, we thank Zebulon Levine for insights on biochemistry experiments and Pu Zheng for advice on probe design for Perturb-seq experiments. From the Elledge Lab, we thank Caleb Glassman and Zack Mirman for input on pINDUCER cell line construction. We also acknowledge Bidisha Mitra from the Gewurz lab for helpful discussions on EBV biology. We also thank Xiaowei Zhuang and Jiaqi Lu for insightful conversations as part of the broader Virome Consortium. We thank the Whitehead Flow Cytometry Core, Whitehead Genome Technology Core, Whitehead Keck Innovation Center, Broad Institute Clinical Labs, Broad Institute Klarman Cell Observatory, and Koch Institute Nanowell Cytometry Platform for technical support. We thank Idan Frumkin and Andrew Murray for thoughtful feedback on the manuscript. This work was conducted as part of the ILLUMINE team, supported by the Cancer Grand Challenges partnership funded by the National Cancer Institute. The authors acknowledge the use of Claude (versions 5.0 and 4.8) to assist with editing and formatting. Some figures were created using elements from BioRender.com.

## Funding

Bill and Melinda Gates Foundation (SJE, MM, JSW)

Chan Zuckerberg Initiative 2024-346405 (5022) (JSW)

Howard Hughes Medical Institute (JSW, SJE)

National Cancer Institute OT2CA321158 (JSW)

NSF GRFP (YHC)

National Institutes of Health P01CA269043 (BEJ)

George and Sandra K. Schussel (BEJ)

DFG – German Research Foundation HO2489/1-1 (KPH)

## Author contributions

Conceptualization: YHC, JSW

Methodology: YHC, LG, ACM, TH, SJE, MM, JSW

Investigation: YHC, LG, ASZ, ACM, TH, EZ, PE, CP, KEY, SS, ENN

Visualization: YHC

Funding acquisition: SJE, KH, BEG, MM, JSW

Supervision: KPH, MM, JSW

Writing – original draft: YHC, JSW

Writing – review & editing: YHC, WS, LG, ACM, KEY, ENN, KPH, BEG, MM, JSW

## Competing interests

JSW declares outside interest in 5 AM Ventures, Amgen, Chroma Medicine, KSQ Therapeutics, Maze Therapeutics, Tenaya Therapeutics, Tessera Therapeutics, Ziada Therapeutics, and Third Rock Ventures. SJE is a founder of TSCAN Therapeutics, MAZE Therapeutics, Infinity Bio, and Mirimus; serves on the scientific advisory boards of Infinity Bio and TSCAN Therapeutics. M.M. is an indirect shareholder in Evosep. The remaining authors declare no competing interests.

## Data, code, and materials availability

The raw sequencing data generated from genetic screens in this study have been deposited in the NCBI Sequence Read Archive (SRA) under BioProject accession PRJNA1516588. The Perturb-seq datasets have been uploaded to the Gene Expression Omnibus (GEO; https://www.ncbi.nlm.nih.gov/geo/) under accession number GSE344535. The RNA-seq datasets have been deposited in GEO under accession GSE342977. The mass spectrometry proteomics data have been deposited to the ProteomeXchange Consortium via the PRIDE partner repository with the dataset identifier PXD08253. All analysis scripts are publicly available at the GitHub repository: https://github.com/cheny4/viral_microproteins

## Materials and Methods

### Cell culture, cell line construction, and lentivirus production

#### Cell lines and culture conditions

The hTERT-immortalized retinal pigment epithelial cell line RPE-1 was cultured in DMEM/F12 basal medium (Thermo Scientific, 11330057). The human chronic myelogenous leukemia line K562 was cultured in RPMI basal medium (Thermo Scientific, 22400-105). The human embryonic kidney line HEK293T (clone c17) and the human melanoma line A375 were cultured in DMEM basal medium (Thermo Scientific, 11965118). Unless otherwise noted, all of the above lines were cultured in basal medium supplemented with 1× penicillin-streptomycin-glutamine (Thermo Scientific, 10378016) and either 10% fetal bovine serum (VWR International, 97068-085, lot 039K24) or 10% Tet System Approved fetal bovine serum (Takara Bio, 631367).

HEK293F cells were maintained in suspension in FreeStyle 293 expression medium (Gibco, 12338-018) at 37 °C and 8% CO₂ on an orbital shaker at 120 rpm between 0.5 and 4 × 10⁶ cells/mL.

The hTERT-immortalized normal oral keratinocyte line NOK was a gift from Ben Gewurz (Brigham and Women’s Hospital, Harvard Medical School) and was cultured in keratinocyte serum-free medium with 0.2 ng/mL EGF and 25 µg/mL bovine pituitary extract (K-SFM Kit; Thermo Scientific, 17005042) supplemented with 5 µg/mL gentamicin (Thermo Scientific, 15710064). All cell cultures were maintained at 37 °C in humidified air with 5% CO₂.

#### Lentivirus production and transduction

Lentivirus for genetic screens and transgene expression was packaged as previously described (*107*). Briefly, lentivirus was produced by co-transfecting HEK293T cells with transfer plasmids and standard packaging vectors (psPAX2, Addgene #12260; pCMV-VSV-G, Addgene #8454) using FuGENE 6 transfection reagent (Promega, E2692). Medium was changed 20 h after transfection, and supernatant was collected 24 h after the medium change and flash frozen on dry ice. Lentiviral transduction was performed by combining single cell suspensions with virus in media supplemented with 8 µg/mL polybrene (Millipore Sigma, TR-1003-G). Adherent cells were seeded after mixing, while K562 suspension cells were spinfected at 33 °C for 75 min prior to seeding at 1 million cells/mL.

#### Flow cytometry and cell sorting

Aria I, Aria II and Aria Fusion cell sorters (BD Biosciences) and a Sony SH800 sorter (Sony) at the Whitehead Institute Flow Cytometry Core, together with a Sony SH900 sorter, were used for sorting. An Attune NxT flow cytometer (Thermo Fisher Scientific) was used for analytical flow cytometry. Antibodies used for sorting and analysis by flow cytometry are summarized in Table S1.

#### Doxycycline-inducible cell line construction

K562 and RPE-1 pINDUCER20 cells were constructed as previously described (*46*). Briefly, K562 and RPE-1 cells were transduced with pINDUCER20-BFP and pINDUCER20-mCD19 (gifts from Stephen J. Elledge, Harvard Medical School). Cells were then "ping-pong sorted" through three sequential rounds of fluorescence-activated cell sorting (FACS). First, cells were induced with 500 ng/mL doxycycline for 48 h, stained with an anti-mouse CD19 antibody (eBioscience, 11-0191-82), sorted to recover the top 20% most mCD19⁺ or BFP⁺ cells, and reseeded in medium without doxycycline. After one week, cells were sorted on the bottom 30% of mCD19⁻ cells and reseeded. After 2–3 days of recovery, cells were induced with 500 ng/mL doxycycline for 48 h and sorted for the top 70% most mCD19⁺ or BFP⁺ cells.

### Viral microprotein library design

#### Curation of viral smORFs for the 50K library

RefSeq accessions for all viruses documented to infect human cells were retrieved from the VirusHostDB database (*43*), yielding 502 GenBank files with complete genomes. Annotations were parsed to obtain all annotated protein-coding sequences, which amounted to 5,921 total ORFs encoding 7,141 proteins, including polyprotein cleavage products. Genome sequences were also analyzed de novo to retrieve all ORFs with ATG or CTG start codons, which recovered 5,456 (92%) of the annotated ORFs; the annotated ORFs that were missed were products of splicing, transcriptional slippage or ribosomal frameshifting.

Retrieved ORFs were then restricted to those encoding <121 aa. Of the 679 annotated products in this size range, 647 were recovered by the de novo search; of the 32 that were missed, 12 arose from splicing, 1 from frameshifting, 4 from start codons other than ATG or CTG, and the remainder from unknown mechanisms. A further 233 short products in this size range were cleavage products of polyproteins. ORFs identified by ribosome profiling of HCMV, HHV-6A/B, KSHV and SARS-CoV-2 (ORF6) were also included. To make the library scale tractable, unannotated ORFs encoding less than 25aa were filtered out, while shorter annotated smORFs and those detected by ribosome profiling were retained. Further information on viral genomes and library sequences can be found in Data S1.

#### Negative control sequences

Four classes of negative control were designed by (i) randomly selecting ORFs and shuffling their nucleotide sequences while maintaining the start codon and removing any stop codons that were generated, (ii) shuffling amino acid sequences, (iii) retrieving ORFs in the antisense direction for viral genomes annotated with directionality, and (iv) generating inert sequences comprising linkers and tags.

#### Oligonucleotide design for the 50K viral smORF library

Once all sequences were retrieved, recognition sites for the BamHI and I-CeuI restriction enzymes were removed by codon optimization. To reduce cost, sequences were sorted into three length tiers: <65 aa (230-nt oligonucleotides), <88 aa (300 nt) and <121 aa (400 nt). Because the smORFs vary in length, filler sequences were added 3′ of each ORF so that amplicon sizes were comparable across the library. 18-bp PCR handles were appended to the ORFs for pooled oligonucleotide synthesis (230 nt, Agilent Technologies; 300 nt and 400 nt, Twist Bioscience). The 5′ PCR handle was designed to have a strong Kozak context, while the 3′ handle was designed to minimize the probability of stop codon readthrough (*108*). The subpool-specific flanking handles are given below:

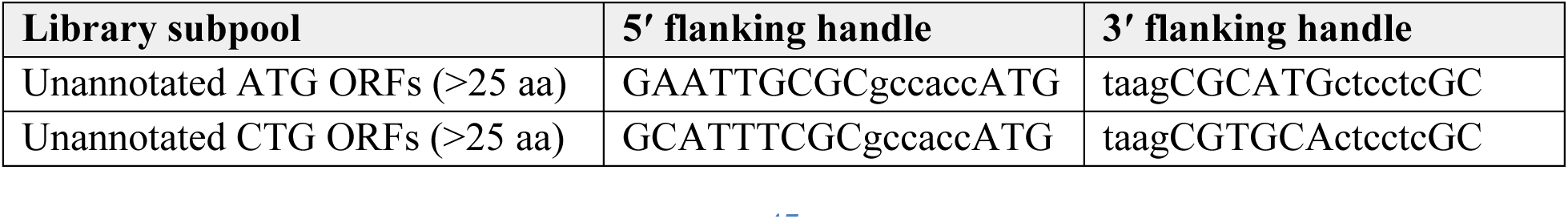
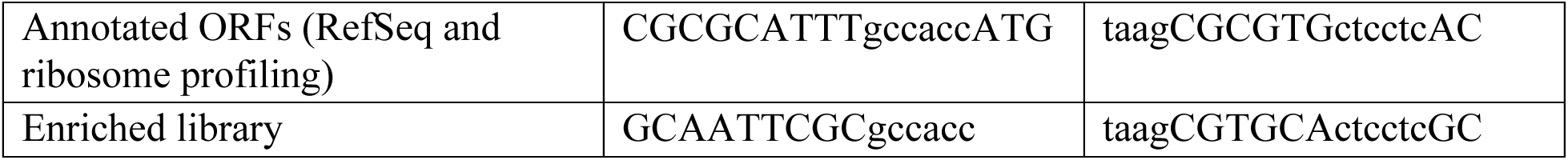

#### Enriched library design

Candidate functional smORFs were curated as described in the main text: (i) toxic hits in both the K562 and RPE-1 screens, (ii) pro-growth hits in either cell line, and (iii) all annotated sequences. Negative controls were regenerated as described above. Oligonucleotides were separated into <88 aa (300 nt) and <121 aa (400 nt) pools (Twist Bioscience). Where space allowed, a 16-bp barcode was added within the filler sequence to improve detection in Perturb-seq. Further information on viral genomes and library sequences can be found in Data S1.

#### Positive control constructs

Positive controls were designed by obtaining amino acid sequences from UniProt and codon-optimizing them with the GenScript codon optimization tool. To make them compatible with pooled genetic screens, the 5’ PCR handle used for smORF amplicon sequencing throughout this study (gatccgctaccttaagagag) was appended downstream of the stop codon, together with 16-bp indel-corrected barcode sequences (*109*). In this design the ORF stop codon and the barcode are separated by 39 bp, a distance within the range associated with minimal ORF–barcode uncoupling arising from PCR or lentiviral recombination, an important source of error in genetic screens (*110*). Downstream of the barcode, filler sequences were added so that the amplicon length matched that of the 300-nt oligonucleotide library. Regions flanking the BamHI and I-CeuI cut sites of pANDORA were added to the ORF–barcode pairs, which were ordered as gBlocks (Integrated DNA Technologies) and either PCR amplified or directly cloned into BamHI/I-CeuI digested pANDORA backbone with NEBuilder HiFi DNA Assembly (New England Biolabs, E2621X).

### Library cloning and amplicon sequencing

#### Viral microprotein library cloning

smORF libraries were cloned by restriction–ligation, similarly to pooled gRNA libraries as previously described (*107*). Briefly, the pANDORA vector was digested with BamHI-HF (New England Biolabs, R3136M) and I-CeuI (New England Biolabs, R0699L) overnight and gel purified (Zymo Research, D4001). Oligonucleotide pools were PCR amplified, with cycle numbers determined by qPCR where amplification was stopped 1-2 cycles before reaching the saturation phase in order to limit PCR bias. Amplicons were column purified (Zymo Research, D4034), digested with BamHI-HF and I-CeuI, gel purified, and then ligated into pANDORA with T4 ligase (New England Biolabs, M0202M) at 16 °C overnight. Ligation products were column purified (Zymo Research, D4034), electroporated into MegaX DH10B competent cells (Thermo Fisher Scientific, C640003) according to the manufacturer’s instructions and grown on LB plates or in liquid culture at 30°C. Plasmids were purified (Qiagen, 12945) and assessed for balance and representation. All libraries had skew ratios between 10 and 16.

#### Amplicon sequencing for library quality control and genetic screens

Genomic DNA extraction, library amplification, quality control and sequencing were performed similarly to previous work (*107*). Briefly, genomic DNA (gDNA) was extracted with a gDNA extraction kit (Macherey-Nagel, 740954.20 or 740954.100). Libraries were first amplified with a primers that amplify the open reading frame (Table S2) using NEBNext Ultra II Q5 PCR Master Mix (New England Biolabs, M0544X); these primers anneal upstream and downstream of the BamHI/I-CeuI restriction sites and carry partial Illumina adapters. For sequencing, 1 µg of plasmid for 12–16 cycles or 5–10 µg of gDNA for 21-24 cycles were used per 100 µL reaction following the manufacturer’s instructions. To add sample indexes, reactions were diluted 1:50 and a second round of PCR was performed with sample indexing primers for 6-8 cycles. Amplicons were isolated by gel purification or 0.8–1× SPRI bead selection (SPRIselect, Beckman Coulter, B23318), analyzed on a High Sensitivity DNA Bioanalyzer (Agilent, 5067-4626) and quantified with the Qubit 1× dsDNA HS Assay Kit (Thermo Fisher Scientific, Q33231). Sequencing was performed on a NextSeq 2000 P1, P2, P3, or P4 kit (Illumina) using a 215-bp read 1, an 85-bp read 2, and 8-bp dual index i7 and i5 index reads.

### Pooled genetic screens

#### Growth screens

Screening was conducted similarly to previous work (*107*). Briefly, positive control ORFs were spiked into each plasmid pool (50K viral smORF library: <65 aa, <88 aa and <121 aa; enriched viral smORF library: <88 aa and <121 aa) and separate lentivirus was made for each. All downstream steps were performed such that each library element was represented by a minimum coverage of ∼1,000× cells/perturbation. RPE-1 or K562 cells were transduced at low MOI (20–30% infection rate) to ensure that the large majority of infected cells received a single perturbation. 24 h after transduction, cells were replenished with complete medium. 48h post transduction, cells were reseeded in medium supplemented with puromycin (Gibco, A1113803) at 5 µg/mL for RPE-1 or 2 µg/mL for K562. After 2 days of selection, cells were recovered in complete medium for 24h. Upon recovery, cells were passaged, marking the initial time point of the screen; an aliquot representing at least 1,000× coverage was frozen down and the remaining cells were reseeded in medium containing 500 ng/mL doxycycline (Sigma Aldrich, D9891-5G). Cells were split every 2 days, and the cell numbers were maintained at a minimum of ∼1,000× cells/perturbation in media containing 500 ng/mL doxycycline. At day 10, cells were harvested, pelleted, and flash frozen on dry ice until gDNA extraction. Growth screens with the 50K Viral smORF library were performed with three replicates, while all screens with the Enriched Library were performed with four replicates.

#### ER stress screens

Screens for survival under ER stress were performed similarly to previous studies (*111*). Briefly, puromycin-selected K562 cells transduced with the enriched library at a minimum coverage of 1,000× cells/perturbation were induced with 500 ng/mL doxycycline and grown in medium supplemented with 100nM thapsigargin (Millipore Sigma, T9033-1MG), 150nM brefeldin A (Selleck Chemicals, S7046), or vehicle alone, for 4 days. Doxycycline, thapsigargin and brefeldin A were replenished every two days when cells were passaged.

*MHC-I screens*

For MHC-I screens, puromycin-selected RPE-1 and A375 cells were induced with doxycycline for 2 days and then stimulated with 200 ng/mL recombinant human interferon-γ (Cell Signaling Technology, 80385S) or PBS controls for 24 h. Cells were harvested and fixed in 4% paraformaldehyde (Thermo Scientific, J61899.AK). Surface MHC-I was stained at 10 million cells/mL with anti-β2-microglobulin antibody (BioLegend, 316318) diluted 1:100 in wash buffer (PBS + 1% BSA) and washed three times. Cell pellets were fixed in 4% PFA and washed three times. The top and bottom 20% of the stained population were sorted on a BD FACSAria (Flow Cytometry Core, Whitehead Institute). To reverse crosslink the samples, fixed cell pellets were resuspended in ChIP lysis buffer (1% SDS, 10 mM EDTA, 50 mM Tris-HCl pH 7.5) and incubated at 65 °C for 10 min. Ambion RNase cocktail (Thermo Scientific, AM2286) was added to a final concentration of 2% (v/v) after samples had cooled to 37 °C, and samples were incubated at 37 °C for 30 min. Proteinase K (New England Biolabs, P8107S) was then added to a final concentration of 10% (v/v) and samples were incubated at 37 °C for 2 h before boiling at 95 °C for 20 min. Samples were cooled on ice, and genomic DNA extraction and sequencing were performed as described above.

#### BNLF2b mutation scanning growth screen

The mutational scanning library of BNLF2b was constructed by swapping individual codons for either GCG (alanine) or AAG (lysine) and ordering the variants as arrayed DNA fragments (Integrated DNA Technologies). The DNA fragments were pooled, amplified by PCR, gel purified, and cloned into the pConstitutive expression vector digested with BamHI and I-CeuI with the NEBuilder HiFi DNA Assembly (New England Biolabs, E2621X). After assembly, plasmids were column purified and amplified by transforming NEB Stable Competent E. coli cells (New England Biolabs, C3040H) following the manufacturer’s instructions. Lentivirus production, 10-day growth screens, and analysis were performed as described above with four replicates.

### Genetic screen analysis

Read quality was assessed with FastQC (*112*). ORF-containing reads were trimmed with a custom Python script (see Data and materials availability) that locates the ORF using the constant sequences flanking the start codon (GCCACCATG) and the 3′ filler, allowing up to one mismatch, and removes the upstream constant region. Trimmed and untrimmed reads were aligned in paired-end mode with bowtie2 (version 2.4.2) (*113*) using default alignment parameters against an index built from the designed library sequences. Alignments were converted to BAM with SAMtools (version 1.11) (*114*), and uniquely mapping reads were retained by filtering on mapping quality (MAPQ ≥ 10) before sorting and indexing. Read counts per library element were obtained from uniquely mapping reads and normalized to the total read count per sample. Log₂ fold changes were computed between the final and initial time points and P values were calculated by Mann-Whitney U test against the distribution of negative control ORFs. P values were corrected for multiple hypothesis testing by the Benjamini-Hochberg procedure. Per-element counts, log₂ fold changes and adjusted P values for every screen are provided in Data S2 to S4.

### Perturb-seq

#### Experimental design

The selection of time points was based on a combination of previously published Perturb-seq experiments and the time scale of doxycycline induction. To ensure that toxic genes were sampled before they depleted from the library, we opted to assay cells at day 1 and day 2, as we observe saturated signals within 16–24 h of doxycycline induction.

RPE-1 cells were transduced with lentivirus made from the two oligonucleotide pools of the enriched library, split after 48 h, and doxycycline induced 4 days after transduction. Day 1 and day 2 doxycycline-induced samples were fixed using the 10x Genomics Fixed Sample Preparation Kit (10x Genomics, PN-1000781) following the manufacturer’s instructions. Cells were then FACS sorted for mCherry+ cells and stored at −80 °C until further processing.

#### Probe design

Probes for detection of library elements were obtained from Integrated DNA Technologies as oPools. Four probe designs for ORF detection were tested: (i) annealing to the 5′ end of the ORF, with right-hand-side probes annealing to the constant 5′ UTR; (ii) annealing to the 3′ end of the ORF, with left-hand-side probes annealing to the constant 3′ UTR; (iii) annealing to the 5′ end of the barcode, with constant right-hand-side probes; and (iv) annealing to the 3′ end of the barcode, with constant left-hand-side probes (fig. S6). The barcode spike-in probes contained the appropriate TruSeq sequences so that they could be amplified separately from the transcriptome probes and sequenced independently. A variable number of N bases was included to ensure base diversity during sequencing. Probe sequences are listed in Data S7.

### Transcriptome and ORF feature library construction and sequencing

Single-cell RNA sequencing was performed with the 10X Genomics GEMX Single Cell Gene Expression Flex platform (10x Genomics, PN-1000794). Spike-in probes for the library elements were included at a final concentration of 2 nM. Cells were counted on a Countess II (Thermo Fisher Scientific). We used four sample barcodes and recovered cells across one pilot experiment, then proceeded with eight sample barcodes for the remaining, totaling to 8 microfluidic channels. mRNA gene expression libraries were prepared according to the manufacturer’s instructions. ORF feature libraries were prepared with the Fixed RNA Feature Barcode Kit according to the manufacturer’s instructions. Libraries were sequenced by Psomagen on an Ultima Genomics platform (UG100).

#### ORF perturbation calling

CRAM files returned by the sequencing provider were converted to FASTQ with samtools fastq. Transcriptome libraries were processed with Cell Ranger (version 9.0.1) (*115*) using the 10x Flex human probe set (Chromium Human Transcriptome Probe Set v1.1.0). Feature libraries carrying ORF–cell barcode assignments were processed with a custom pipeline (see Data and materials availability): reads were first partitioned by probe type and split into ORF-derived and cell barcode/UMI-derived reads, then aligned with STARsolo (version 2.7.1a) (--soloType CB_UMI_Simple, 16-nt cell barcode, 12-nt UMI, --alignEndsType Local, --outFilterMultimapNmax 20, --outFilterMatchNmin 20) (*116*) against an index of the designed library sequences, using the Cell Ranger cell barcode whitelist. Per-element counts were tabulated with featureCounts (version 2.0.1) (*117*), and probe barcodes were resolved from the paired reads of uniquely assigned alignments. Perturbations were then called using a mixture model that jointly fits background and true ORF assignments, as previously described (*39*). A total of 882,225 out of the 1,231,800 cells recovered had at least one ORF assignment, among which 780,731 cells were singlets, and 101,494 had more than one ORF assigned.

#### Energy test and Pearson distance calculation

Pseudobulk matrices were calculated by taking the mean z-normalized counts for each perturbation. Pairwise Pearson distances between perturbations were calculated from this pseudobulk matrix with pertpy (version 1.0.3) (*118*), and energy tests were performed as previously described (*63*), using the first 20 principal components of the z-scored count matrix. The test statistic was the energy distance between the cells carrying a given ORF and the negative control reference group, and P values were obtained by random permutation of perturbation labels (500 permutations). The Benjamini-Hochberg procedure was used for multiple testing correction of the permutation-test P values. Energy distances, FDRs and differentially expressed gene counts for every perturbation are reported in Data S5.

#### Differential gene expression analysis

Differential expression was computed with scanpy (version 1.11.5) (*119*) using the scanpy.tl.rank_genes_groups function on the log1p-normalized layer, using a Wilcoxon rank-sum test with the negative control population as the reference group. Differentially expressed genes were filtered based on adjusted p-value <0.05.

#### Minimum distortion embedding of strong perturbations

The visualization in Fig. 3B is a minimum distortion embedding computed as previously described (*39*). Following that approach, the embedding was restricted to perturbations with a strong transcriptional phenotype (energy test FDR<0.05), and perturbations were compared using the correlation of pseudobulk expression profiles across highly variable genes, a scale-invariant measure of similarity that is robust to differences in the magnitude of effect between related perturbations. For each retained perturbation, a pseudobulk profile was computed as the mean normalized expression of cells transduced with a given ORF, restricted to the top 2,500 most highly variable genes.

Two embeddings were computed from this representation using a shared k-nearest-neighbor graph with k = 3. First, a 20-dimensional spectral embedding was computed with sklearn.manifold.SpectralEmbedding (affinity=’nearest_neighbors’, n_neighbors=3, eigen_solver=’arpack’). Second, the two-dimensional minimum distortion embedding shown in Fig. 3B was computed with the pymde (version 0.1.18) function pymde.preserve_neighbors (embedding_dim=2, n_neighbors=3, repulsive_fraction=3, attractive penalty pymde.penalties.Log1p, random initialization).

#### Clustering, gene program analysis, and manual annotations

To identify clusters of related perturbations, we combined two clustering methods. First, we applied HDBSCAN (metric=’precomputed’, min_cluster_size=3, min_samples=1, cluster_selection_method=’eom’) to the pairwise Pearson distance matrix. This procedure is intrinsically conservative given the choice of metric and clustering algorithm, so many perturbations are not assigned to any cluster; our emphasis was on identifying the strongest signals rather than the most comprehensive set. Second, Leiden clustering was performed on the 20-dimensional embedding described above: a k-nearest-neighbor graph was constructed with scanpy.pp.neighbors using Euclidean distance and k = 3 neighbors, and communities were detected with scanpy.tl.leiden at resolution 1.0.

For interpretation of cluster phenotypes, gene sets for transcriptional programs relevant to RPE-1 biology were obtained from genome-scale Perturb-seq datasets (*39*, *64*). Scores were computed with scanpy.tl.score_genes with default parameters. Using the two cluster assignments, gene program score matrix, and taking into account existing knowledge of the individual viral ORFs, the clusters were manually assigned and putative functions were annotated for 11 clusters.

### Arrayed plasmid cloning for mechanistic follow-up

All smORF screen hits and other control ORFs were codon optimized using the GenScript codon optimization tool and ordered as gBlocks (Integrated DNA Technologies). Plasmids were cloned as described for the BNLF2b mutagenesis library above. Toxic ORFs were cloned into the pANDORA vector backbone digested with BamHI/I-CeuI, while pro-growth ORFs were cloned into the pConstitutive backbone digested with BamHI/I-CeuI. Plasmid sequences were verified by nanopore sequencing performed by Quintara Biosciences. ALFA-tagged smORFs for high-throughput AE-MS were ordered from Twist Biosciences in the pTwist Lenti EF1a EGFP Puro backbone.

### High-throughput affinity-enrichment mass spectrometry

Affinity-enrichment mass spectrometry (AE-MS) for identification of viral microprotein interactors was performed as previously described (*40*). The pipeline is briefly outlined below.

#### Cell culture and expression of ALFA-tagged viral microproteins

HEK293T cells were maintained in DMEM (high glucose; Capricorn Scientific, DMEM-HPSTA) supplemented with 10% FBS and 1% penicillin–streptomycin (Gibco, 15140-122) at 37 °C and 5% CO₂, grown in 10- or 15-cm dishes (Sarstedt, 83.3902/83.3903) and passaged at 80–90% confluency with 0.05% trypsin–EDTA. Cells were seeded into 96-well plates (Sarstedt, 83.3924.005) at 40,000 cells per well in 90 µL and grown for 24 h to ∼50% confluency. Per well, 1 µL of plasmid DNA (100 ng µL⁻¹) in 4µL Opti-MEM (Gibco, 31985-062) was incubated for 5 min and combined with 0.2 µL Lipofectamine 2000 (Invitrogen, 11668019) diluted in 5 µL Opti-MEM; complexes were formed for 30 min and 10 µL was dispensed per well with an INTEGRA Mini 96 (Integra Biosciences, 4801). Transfections were performed in four replicates per construct and cells were harvested 16 h post-transfection. The arrayed library comprised 380 distinct viral baits including GFP11 and empty (‘nonè) constructs as negative controls, plus a set of control proteins. For the RPE1 experiment, stable RPE1 lines expressing nine selected ALFA-tagged viral proteins were generated by lentiviral transduction rather than transient transfection.

#### ALFA-tag affinity enrichment

Wash buffer was 50 mM Tris-HCl pH 7.5 (at 4 °C), 150 mM NaCl, 5% glycerol. IP lysis buffer was wash buffer plus 0.5% IGEPAL CA-630.Lysis buffers were freshly supplemented with cOmplete EDTA-free protease inhibitor (one tablet per 50 mL; Roche, 05056489001), 1 mM MgCl₂ and 0.1% (v/v) in-house benzonase.

Cells were lysed on-plate in 100 µL per well of ice-cold lysis buffer with the INTEGRA Mini 96, mixed by pipetting 30 cycles with 60 µL, shaken at 950 r.p.m. for 30 min at 4 °C, transferred to Eppendorf Twintec 96-well plates (0030 129.512), and cleared at 4,000 × g for 15 min at 4 °C. Streptavidin-coated 384-well plates (Thermo Scientific, 15405) were washed three times with 100 µL per well of wash buffer on an automated plate washer (Cytena C.Wash Plus).

Biotinylated anti-ALFA nanobody (NanoTag, N1505-Biotin, or produced in house) was diluted to 2.5 ng µL⁻¹ and dispensed at 50 µL per well (125 ng per well), incubated for 2 h at 25 °C and 1,200 r.p.m., washed three times with wash buffer, sealed and stored at 4 °C. Cleared lysate was transferred to the nanobody-coated plate at 50 µL per well, sealed and incubated 12–17 h at 4 °C and 1,200 r.p.m., briefly centrifuged, and washed three times with 100 µL per well of ice-cold wash buffer on the automated plate washer.

#### On-plate proteolytic digestion

Digestion buffers were prepared fresh. Buffer 1 (6 M urea, 10 mM Tris-HCl pH 8.5, 3 mM DTT 1 ng µL⁻¹ LysC) was dispensed at 20 µL per well and incubated for 3 h at 30 °C and 1,200 r.p.m. Buffer 2 (10 mM Tris-HCl pH 8.5, **0.4 ng µL⁻¹. Promega Trypsin Gold MS-grade (V5111), 7.5 mM 2-chloroacetamide) was added at 40 µL per well and digestion continued overnight at 37 °C** and 1,200 r.p.m. Reactions were quenched with 6.6 µL per well of 10% TFA. Evotips (Evosep Biosystems, EV2011) were activated with 50 µL of buffer B (acetonitrile, 0.1% FA) at 700 × g for 1 min, soaked in 2-propanol for 1 min, and washed with 50 µL of buffer A (water, 0.1% FA) at 700 × g for 1 min. Acidified peptides were loaded at 500 × g for 2–3 min and washed with 250 µL of buffer A at 400 × g for 1 min. Tips were stored at 4 °C in buffer A until LC-MS.

#### Liquid chromatography and mass spectrometry

Peptides were separated on an Evosep Eno using the 500 samples-per-day method for the smORF library (2.3 min total method duration) and on the Evosep One with the 200 samples-per-day method for the RPE1 experiment (5.6 min total method duration), as recorded in the instrument methods. Data were acquired in data-independent acquisition mode on an Orbitrap Astral Zoom (smORF library;2,304 runs) or an Orbitrap Astral (RPE1; 40 runs) mass spectrometer, operated in positive ion mode with label-free, unfractionated single-shot injections (one fraction, one technical replicate per sample).

Ionization was performed by nano-electrospray (NSI) with a static spray voltage of 1.9 kV (smORF library) or 2.0 kV (RPE1) and an ion transfer tube temperature of 280 °C. A FAIMS Pro interface was used in all experiments, operated in standard-resolution mode at a single compensation voltage of −40 V, with a total carrier gas flow of 3.5 L min⁻¹. The RF lens was set to 40% and the expected peak width to 10 s. smORF library (Orbitrap Astral Zoom, 500 SPD, 2.3 min method). MS1 survey scans were acquired in the Orbitrap at a resolution of 240,000 over m/z 380–980 in profile mode, with a normalised AGC target of 500% (absolute 5 × 10⁶) and a maximum injection time of 3 ms. DIA MS2 scans were acquired in the Astral analyser using 113 variable-width isolation windows spanning m/z 380.5–980.0 (width 4.03–12.99 Th, median 5.44 Th), with HCD at a normalised collision energy of 25%, a scan range of m/z 150–2000, a maximum injection time of 5 ms, a normalised AGC target of 500% (absolute 5 × 10⁴), and centroid data type.

RPE1 experiment (Orbitrap Astral, 200 SPD, 5.6 min method). MS1 survey scans were acquired in the Orbitrap at a resolution of 240,000 over m/z 380–1280 in profile mode, normalised AGC target 500% (absolute 5 × 10⁶), maximum injection time 3 ms. DIA MS2 scans were acquired in the Astral analyser using 150 fixed 4 Th isolation windows spanning m/z 380.4–980.7 with no window overlap and window placement optimisation enabled, HCD normalised collision energy 25%, scan range m/z 150–2000, maximum injection time 5 ms, normalised AGC target 500% (absolute 5 × 10⁴), centroid data type.

#### Protein abundance quantification and differential enrichment analysis

Raw files were processed with DIA-NN (*120*). The 1,536 runs of smORF plates 1–4 were searched individually with DIA-NN 2.1.0 against an in silico predicted spectral library generated with DIA-NN 2.3.1 from a viral smORF FASTA combined with the human SwissProt reference (2026-03-30). The 768 iteration-2 runs were searched together with DIA-NN 2.1.0 with match-between-runs enabled, against a predicted library built with DIA-NN 2.3.1 from the viral microprotein FASTAs and human SwissProt (2024-07-22). The 40 RPE1 runs were searched with DIA-NN 1.9.1 using a library-free FASTA digest against human SwissProt (2024-07-22) supplemented with the bait sequences, with match-between-runs enabled.

In all searches, in silico digestion allowed cleavage C-terminal to lysine and arginine with up to one missed cleavage and N-terminal methionine excision. Peptide length was restricted to 6–30 residues (7–30 for iteration 2; 5–31 for RPE1), precursor charge to 1–4, precursor m/z to 300–1,800 and fragment m/z to 200–1,800. Carbamidomethylation of cysteine was set as a fixed modification. Methionine oxidation was variable for the smORF searches, with protein N-terminal acetylation additionally variable for plates 1–4; the RPE1 search used no variable modifications. Precursor and protein identifications were filtered at 1% FDR, with heuristic protein inference and retention-time profiling enabled.

Protein abundances were quantified per condition by both MaxLFQ from the DIA-NN protein-group table and directLFQ (version 0.3.3) (*121*) from proteotypic precursors (Q value < 0.01) as previously described. Briefly, differential enrichment was performed on imputed directLFQ values and each bait gene was contrasted against all other samples, including the control wells, using an empirical-Bayes moderated t-test (alphapepttools, version 0.2.0) (*122*). Multiple hypothesis correction was performed by Benjamini–Hochberg, and significant protein interactors were called with FDR < 0.05 and log₂ fold-change > 1.

#### Protein network visualization

Significant protein interactions were visualized with Cytoscape (version 3.10.4) (*123*) and merged with the host protein network using the STRING application filtering edges with confidence scores >0.7. Edge weights between the viral smORF bait and host protein interactor were determined by multiplying the log₂ fold change by −log₁₀(FDR) and color intensity of edges corresponded to log₂ fold change.

### Bulk RNA sequencing

To define the transcriptional programs driven by individual microproteins, hTERT-RPE-1 cells were stably transduced with lentiviral constructs expressing GFP11 (negative control), the HPV16 E7 oncoprotein, EBV BNLF2b WT and mutants, or PKIA. After puromycin selection, cells were pelleted and lysed in Zymo DNA/RNA Shield (Zymo Research, R1100-50) and processed by Plasmidsaurus. Total RNA was profiled by 3′-end counting RNA sequencing in four biological replicates per condition. DESeq2 (*124*) was used for differential gene expression analysis of all ORFs against the GFP11 control cells as well as BNLF2b versus PKIA overexpression samples.

### Immunoprecipitation-mass spectrometry and western blotting

For immunoprecipitation experiments, transduced RPE-1or A375 cells were seeded in 6-well plates and allowed to grow to 70% confluence before harvesting with TrypLE Express (Thermo Scientific, 12604013). TrypLE was neutralized with an equal volume of complete growth medium. Cells were washed once with DPBS (Thermo Scientific, 14190144) and flash frozen on dry ice. Thawed cells were lysed by resuspension in Pierce IP Lysis Buffer (Thermo Scientific, 87787) supplemented with 1× cOmplete EDTA-free Protease Inhibitor Cocktail (Millipore Sigma, 4693159001) and incubation for 30 min at 4 °C on a revolver rotator (Thermo Scientific, 88881001). Anti-ALFA immunoprecipitation was performed using ALFA Selector ST (NanoTag Biotechnologies, N1516-L) following the manufacturer’s instructions with slight modifications: for western blots, samples were eluted by direct boiling in 1× SDS sample buffer; for mass spectrometry, samples were washed four additional times with 10 mM Tris-HCl pH 8.0 (Millipore Sigma) and submitted to the Taplin Mass Spectrometry Facility (Harvard Medical School). Antibodies used for western blotting are summarized in Table S1.

### Arrayed Growth competition assays

Growth competition assays were performed under conditions similar to the original screens. RPE-1 cells were transduced at 70% confluence at approximately 30% infection rate and seeded in 24-well plates. Every two days, three quarters of the cells were collected and analyzed on an Attune NxT flow cytometer (Thermo Fisher Scientific) and the remaining cells were reseeded.

Time points were collected up to 12 days after seeding. K562 growth competition assays were performed following similar procedures with the following modifications: cells were transduced at 250,000 cells/mL, and 500 ng/mL doxycycline was added the day before the initial time point for inducible expression vectors. Where the initial titer exceeded 50%, cells were selected with 7.5 µg/mL puromycin and pooled with untransduced cells to reach a ratio of approximately 30%. For growth competition assays with drug treatment, drugs were added fresh each time cells were split.

### Arrayed Growth rate measurements

Incucyte growth assays were performed following the instrument manufacturer’s instructions. Cells were seeded at approximately 1% confluence the day before the initial time point. Medium was exchanged for fresh medium before loading onto the Incucyte platform (S3 system). A plate scan was performed every 4 h for a total of 96 h (25 time points) and confluence was calculated for each time point using the Incucyte AI Confluence Analysis software (Sartorius). Growth rate was calculated as the mean of the steepest increase in confluence between two consecutive 4h measurements.

### RPE-1 soft agar colony formation assays

Soft agar colony formation assays were performed as previously described (*125*). Briefly, a bottom layer of 1% noble agar (Thermo Scientific Chemicals, J10907.22) mixed with an equal volume of 2× DMEM/F12 medium (Gibco, 12500062) was distributed into 6-well plates. Once this layer had solidified, an upper layer was added, consisting of a single-cell suspension in complete DMEM/F12 medium (3,000, 6,000 or 9,000 cells per well) mixed with 0.6% noble agar. Plates were then incubated in a humidified 37 °C cell culture incubator for 40–50 days, with 100 µL of medium added twice weekly. Plates were imaged on an EVOS M5000 imaging system (Thermo Fisher Scientific) using phase contrast and red fluorescence.

### ISRE luciferase reporter assay

HEK293T cells were seeded in 24-well plates at approximately 40% confluence (40,000 cells per well) and transfected the following day with a total of 0.4–0.5 µg DNA per well, comprising equimolar amounts of the ISRE reporter plasmid pNlucP-ISRE-Hygro (Promega, E4141), the firefly luciferase normalization plasmid pYC701, and the ORF expression plasmid, complexed with FuGENE HD (Promega, PAE2311) at a 3:1 reagent-to-DNA ratio in Opti-MEM. Three biological replicates were transfected per condition, with GFP (pYC544) and GFP11 (pYC488) constructs as negative controls. Two days after transfection, cells were split into a 96-well flat-bottom plate and stimulated with 75 U/mL recombinant human interferon-α A (Millipore Sigma, IF007) or left unstimulated. Approximately 16 h after stimulation, NanoLuc and firefly luciferase activities were measured with the Nano-Glo Dual-Luciferase Reporter Assay System (Promega, N1610) following the manufacturer’s instructions, and ISRE reporter activity was expressed as the ratio of NanoLuc to firefly luminescence normalized to the unstimulated control.

### Forskolin treatment and phospho-PKA substrate blotting

Cells were treated with either 20 µM forskolin (R & D Systems, 109910) in DMSO or a DMSO-only vehicle control for 30 min. Cells were lysed in Pierce IP Lysis Buffer (Thermo Scientific, 87787) supplemented with 1× cOmplete EDTA-free Protease Inhibitor Cocktail (Millipore Sigma, 4693159001) and 1× phosphatase inhibitor cocktail (Thermo Scientific, A32957). Blots were probed with an antibody against phosphorylated PKA substrates (Cell Signaling Technology, 9624).

### PKA activity assay

The PKA activity assay (Thermo Scientific, EIAPKA) was performed according to the manufacturer’s instructions with small modifications. Briefly, transduced hTERT-RPE-1 cells were selected with 7.5 µg/mL puromycin for 2 days and seeded the day before collection so as to reach 70% confluence at the time of harvest. Cells were lysed with the lysis buffer supplied with the kit, supplemented with 1× cOmplete EDTA-free Protease Inhibitor Cocktail (Millipore Sigma, 4693159001) and 1× phosphatase inhibitor cocktail (Thermo Scientific, A32957). The reaction, washes and readout were performed following the manufacturer’s instructions.

### Immunofluorescence microscopy

Samples for immunofluorescence were prepared according to the instructions of the manufacturers of the corresponding antibodies. Briefly, for phospho-STAT1 staining, 10,000 cells were seeded in 24-well coverslip-bottom plates (Ibidi, 82426) the day before transfection with 250 ng of target plasmid per well. 48 h after transfection, cells were treated with 200 ng/mL recombinant interferon-γ (Cell Signaling Technology, 80385) in DPBS (Thermo Scientific, 14190144) for 45 min. Cells were briefly washed with DPBS and fixed in 4% paraformaldehyde at room temperature for 10 min. After washing with DPBS, cells were permeabilized with 100% ice-cold methanol at −20 °C for 10 min, washed with DPBS, and further permeabilized and blocked in DPBS supplemented with 3% normal goat serum (Abcam, ab7481-50ml) and 0.1% Triton X-100 (Life Technologies, 85111) at room temperature for 30 min. The antibody against phospho-STAT1 (Cell Signaling Technology, 9167) was preconjugated with an anti-rabbit IgG secondary nanobody (NanoTag Biotechnologies, N2402-AF568-S) according to the manufacturer’s instructions. The conjugated antibody and an Atto643-conjugated anti-ALFA nanobody (NanoTag Biotechnologies, N1502-At643-L) were diluted in DPBS supplemented with 3% normal goat serum and 0.1% Triton X-100 to final dilutions of 1:500 and 1:1000, respectively. Cells were incubated with the diluted antibodies overnight at 4 °C, protected from light. After washing with DPBS and staining with 1 µg/mL DAPI (Thermo Scientific, 62248) in DPBS, cells were imaged on a Nikon CrestOptics X-Light spinning disk confocal microscope with a 40× water-immersion objective.

### Phylogenetic analysis

The poxvirus phylogenetic tree was obtained from the International Committee on Taxonomy of Viruses (*126*), with selected representative nodes shown. The astrovirus phylogenetic tree was constructed from genome sequences of the most similar astrovirus genome sequences to HMO Astrovirus A NC_013443.1 and the classical human astrovirus clade based on another study (*78*). ORF1b (RdRp) coding sequences were identified by locating the ribosomal frameshift motif and translating in the −1 frame; ORF2 (capsid) sequences were identified by the presence of the subgenomic promoter motif. Amino acid sequences were aligned with MUSCLE (version 5.1) (*127*) and alignments were converted to NEXUS format with a custom python script. Trees were inferred with MrBayes (version v3.2.6) (*128*) using the following block:

begin mrbayes;
set autoclose=yes nowarn=yes;
prset aamodelpr=fixed(wag);
lset rates=invgamma;
mcmc ngen=1000000 samplefreq=500 printfreq=1000 diagnfreq=5000 nruns=2
nchains=4;
sump burnin=250;
sumt burnin=250;
end;

Trees were visualized with iTOL (*129*). Scripts for sequence extraction, alignment and NEXUS file generation are provided in the code repository (see Data and materials availability).

### Structure prediction

Structures and complexes were predicted with the AlphaFold3 Server implementation of AF3 with default parameters (*60*) and visualized in PyMOL (version 2.1) (*130*). All interpretations were made from predicted complex structures with ipTM > 0.75.

### Amino acid sequence similarity analysis

Pairwise amino acid sequence similarity was computed with the Biopython PairwiseAligner in local alignment mode using the BLOSUM62 substitution matrix, with a gap-open penalty of −11 and a gap-extension penalty of −1.

---

**Fig. S1.**
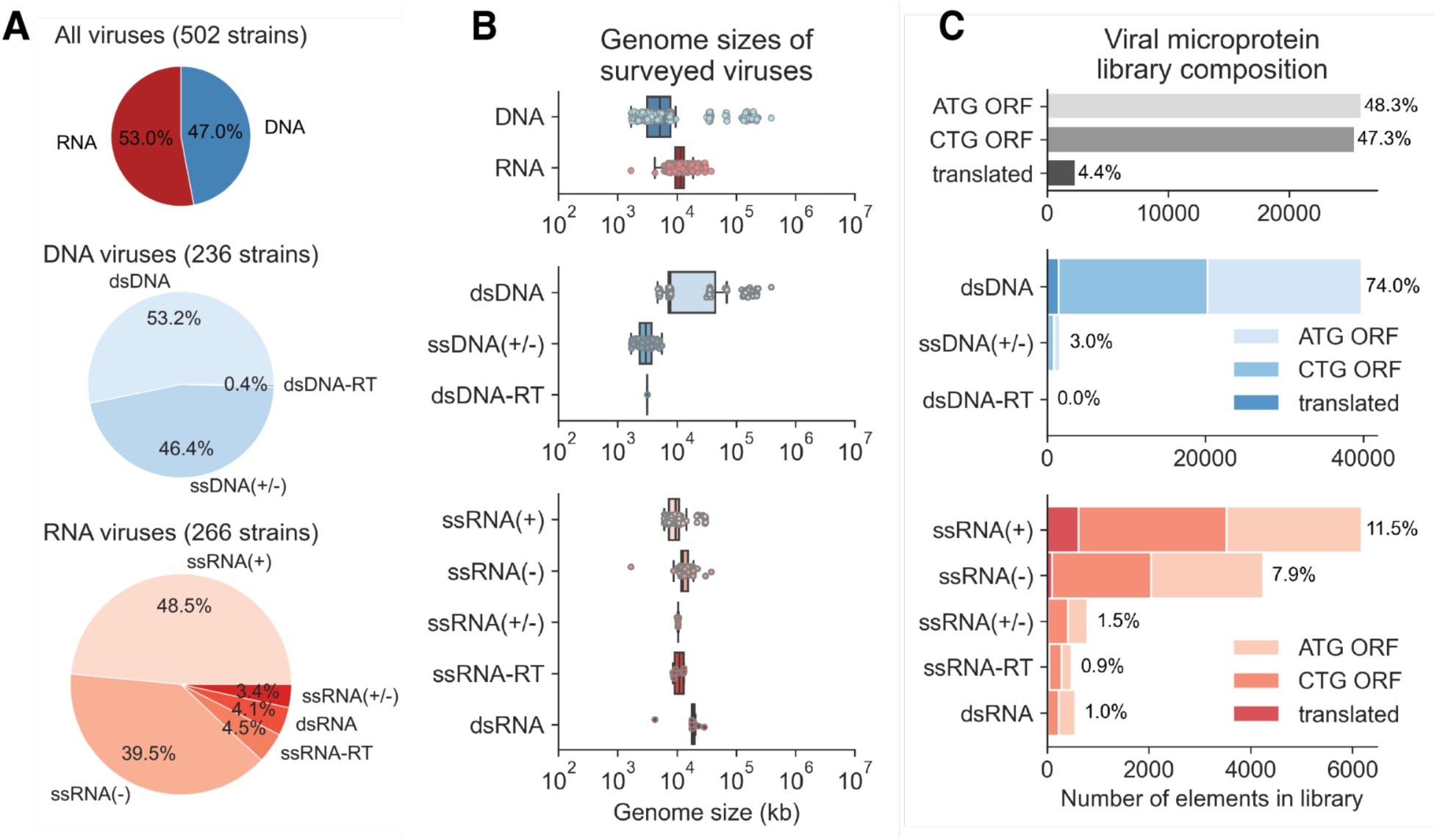
Survey of viral genomes for smORFeome curation. **(A)** Baltimore classifications of the viral genomes. (**B**) Genome sizes. (**C**) Fraction of ORFs derived from each classification.

**Fig. S2.**
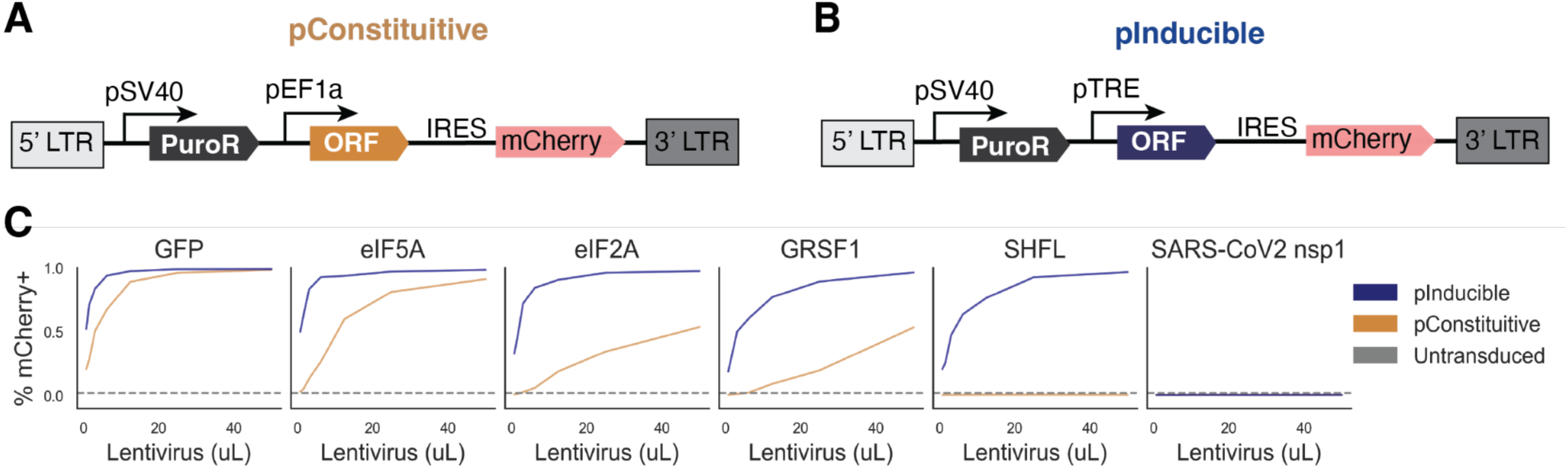
Toxic ORFs do not produce viable lentivirus. Lentiviral vector designs for (**A**) constitutive and (**B**) inducible ORF expression. ORFs are encoded on a polycistronic transcript with an IRES-mCherry. (**C**) Fraction of mCherry-positive cells 4 days after transduction and 1 day of dox induction with indicated ORFs titrated with different volumes of lentivirus.

**Fig. S3.**
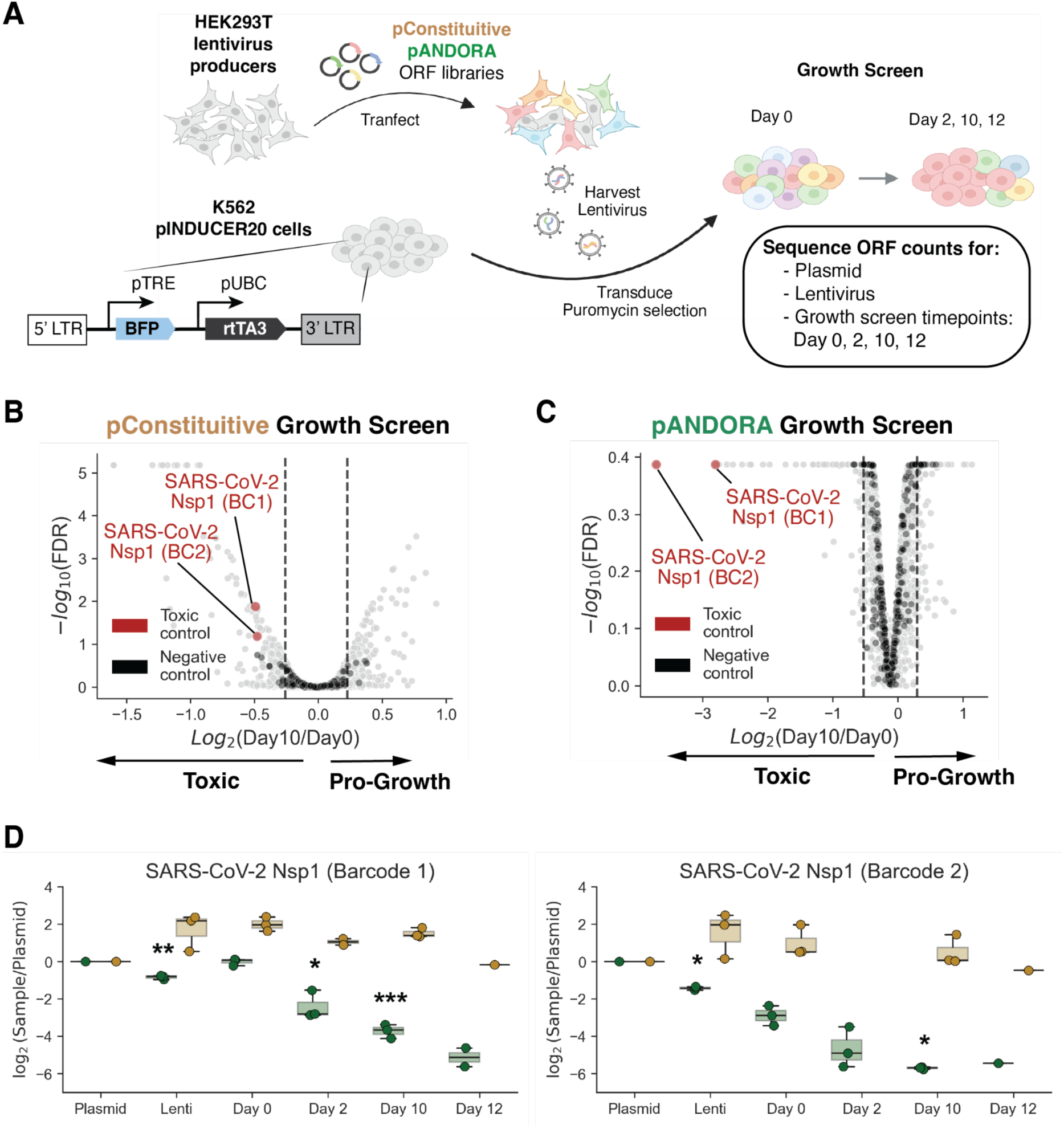
Leakiness of inducible ORF vectors impacts sensitivity of toxic ORF detection in pooled ORF screens. (**A**) Schematic of the screen design for evaluating the sensitivity of the pConstitutive and pANDORA vectors cloned with the 66-88aa smORF library. K562 growth screen results for the (**B**) pConstitutive and (**C**) pANDORA libraries. (**D**) SARS-CoV-2 Nsp1 ORF counts at the plasmid, lentivirus, day 0, day 2, day 10, and day 12 stages.

**Fig. S4.**
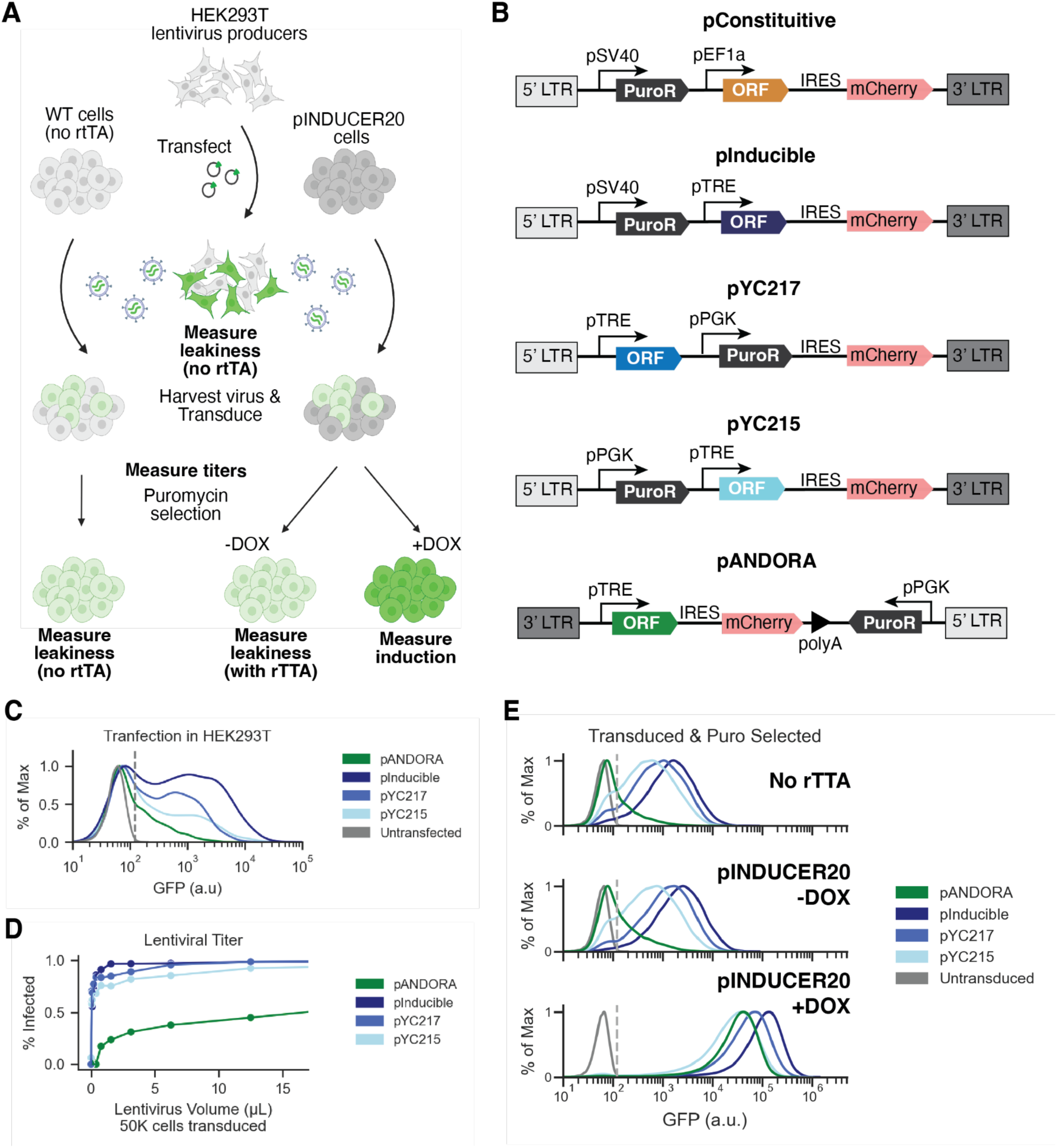
Optimization of a minimally leaky inducible vector for toxic ORF expression. (**A**) Schematic for examining titers, induction, and leakiness at different stages of the genetic screening pipeline. (**B**) Vector designs for inducible ORF overexpression; GFP was cloned into all ORF vectors for testing. (**C**) HEK293T producer cells that do not express rtTA were transfected, and GFP expression was examined by flow cytometry 48 h after transfection. (**D**) K562 cells were transduced with the indicated volumes of lentivirus produced from GFP ORF vectors, grown in doxycycline media for 72 h, and assessed for the fraction of GFP-positive cells. (**E**) GFP levels 72 h after transduction.

**Fig. S5.**
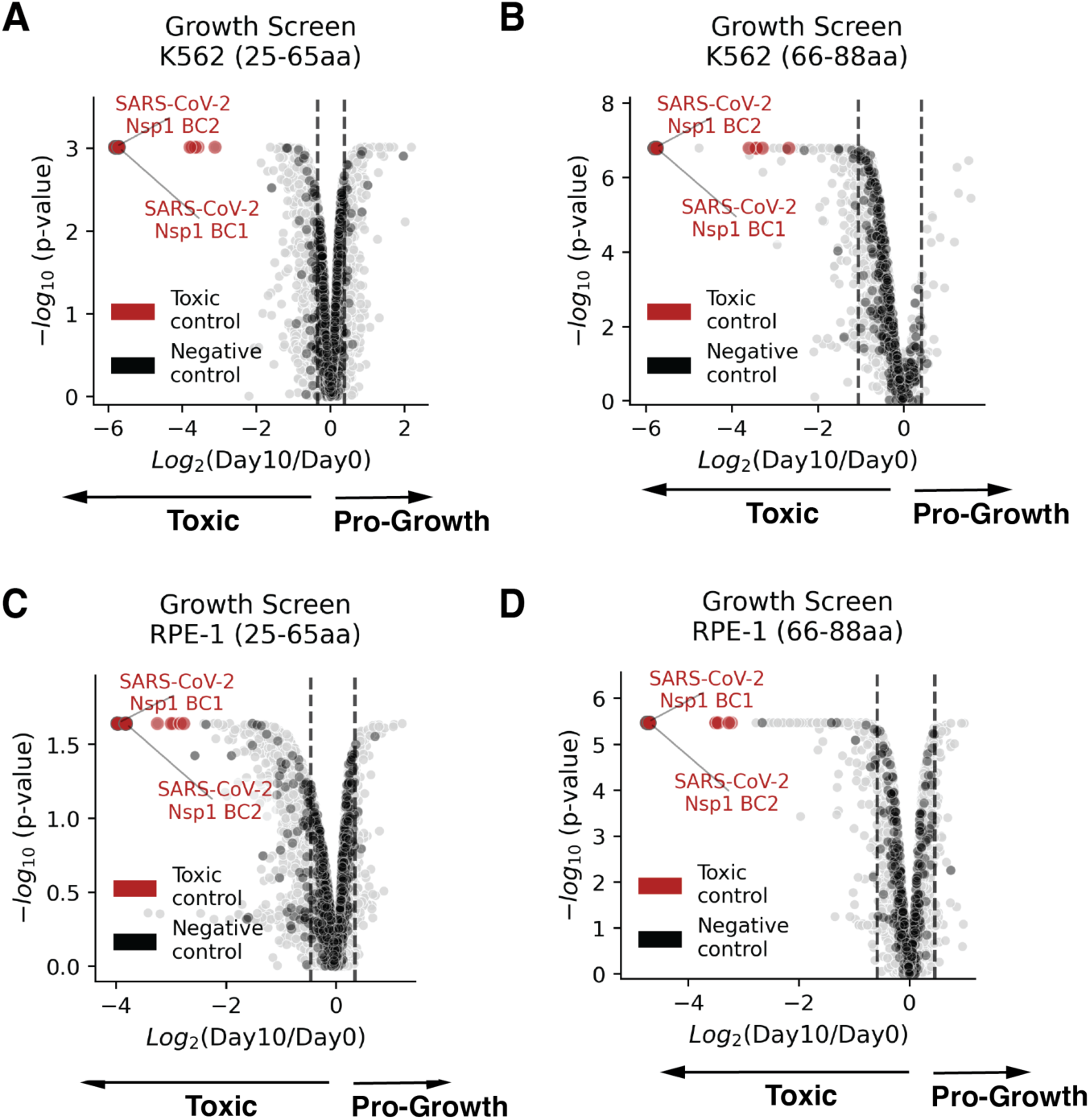
Volcano plots of growth screens for viral smORFeome sublibraries 25-88aa in length. 10-day K562 screens with the (**A**) 25-65aa and (**B**) 66-88aa libraries, and RPE-1 screens with the (**C**) 25-65aa and (**D**) 66-88aa libraries.

**Fig. S6.**
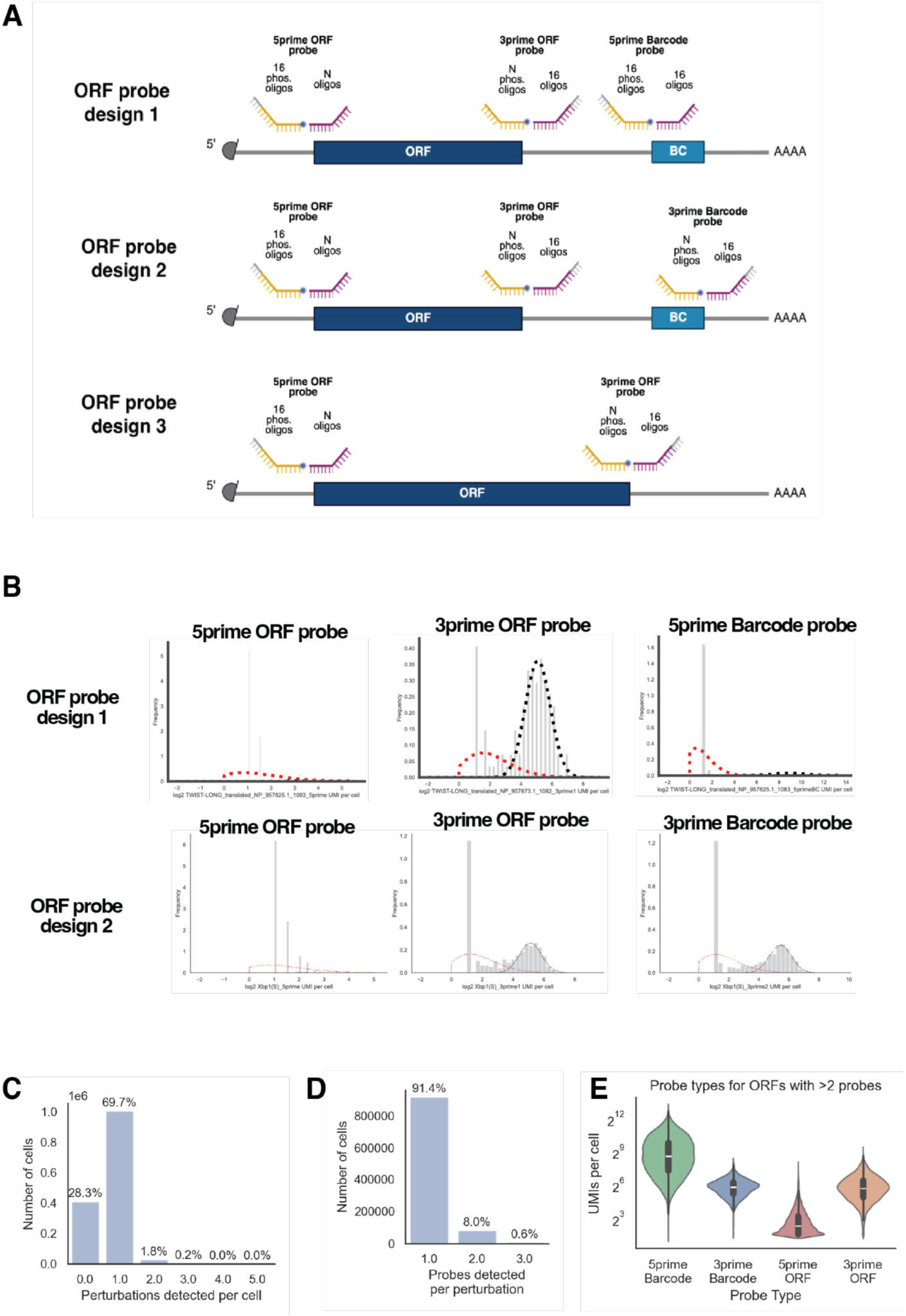
Perturb-seq custom probe design and ORF calling. (**A**) Schematic of the barcoded ORF vector and probe design. (**B**) Example of mixed-model fitting of probe UMI counts for ORF assignment. Shown are two ORFs with three possible probe designs. (**C**) Number of smORF perturbations detected per cell. (**D**) Number of probes detected for cells with one perturbation. (**E**) Violin plots of UMI counts for the different probe types.

**Fig. S7.**
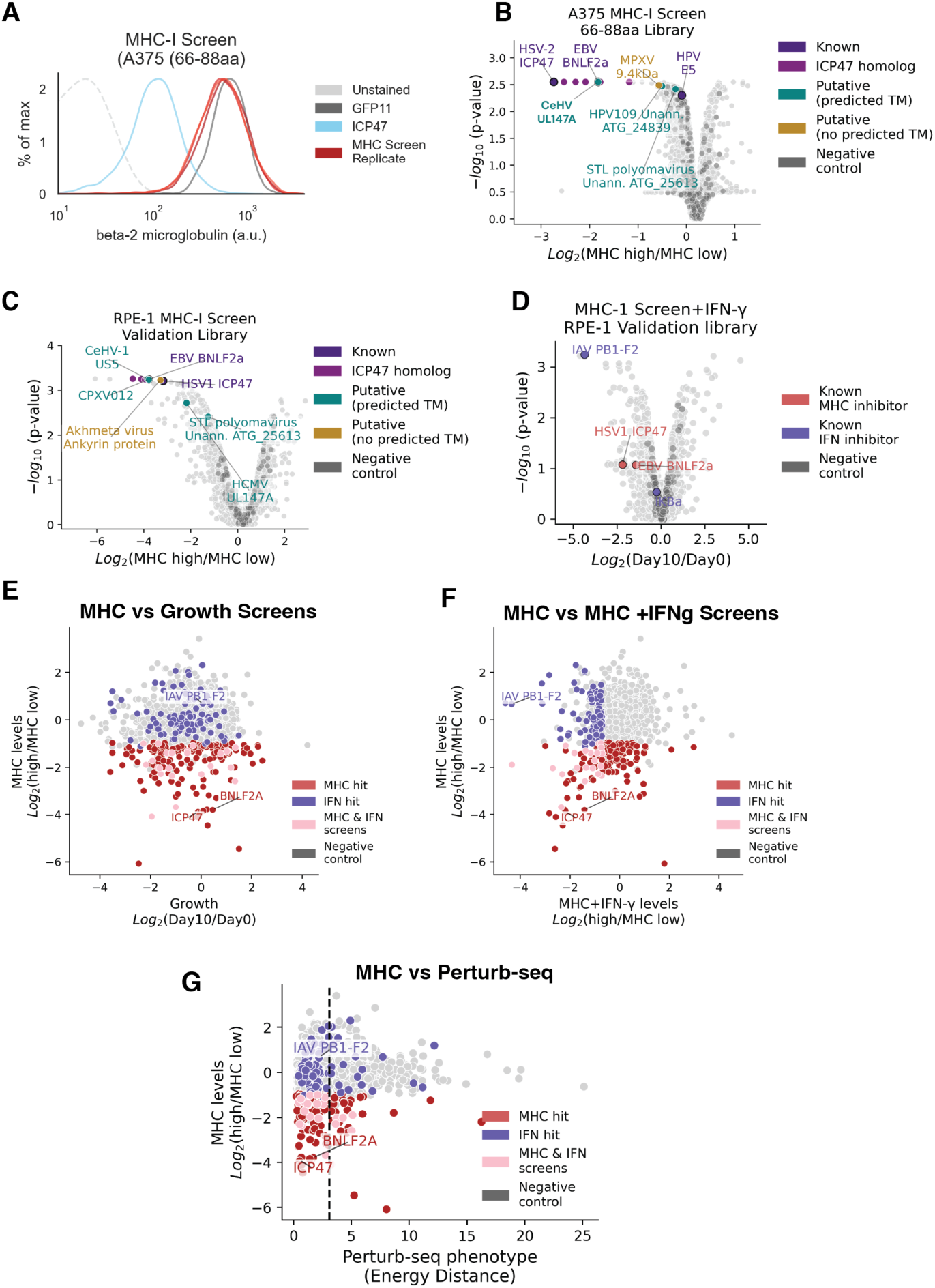
MHC screen hits do not have growth or Perturb-seq phenotypes. (**A**) MHC-I levels measured by beta-2-microglobulin staining on A375 cells transduced with the 66-88aa viral smORF library, compared with positive (ICP47) and negative (GFP11) controls. Volcano plots of screen results for surface MHC-I levels in (**B**) A375 cells transduced with the 66-88aa viral smORF library and (**C**) RPE-1 cells transduced with the enriched library under basal conditions and (**D**) after 24 h of 200 ng/mL IFN-γ stimulation. Scatter plots of phenotypes from (**E**) the RPE-1 surface MHC-I screen versus the 10-day growth screen, (**F**) the RPE-1 surface MHC-I screen without versus with IFN-γ stimulation, and (**G**) the RPE-1 surface MHC-I screen versus the Perturb-seq phenotype score, calculated as the energy distance from negative controls.

**Fig. S8.**
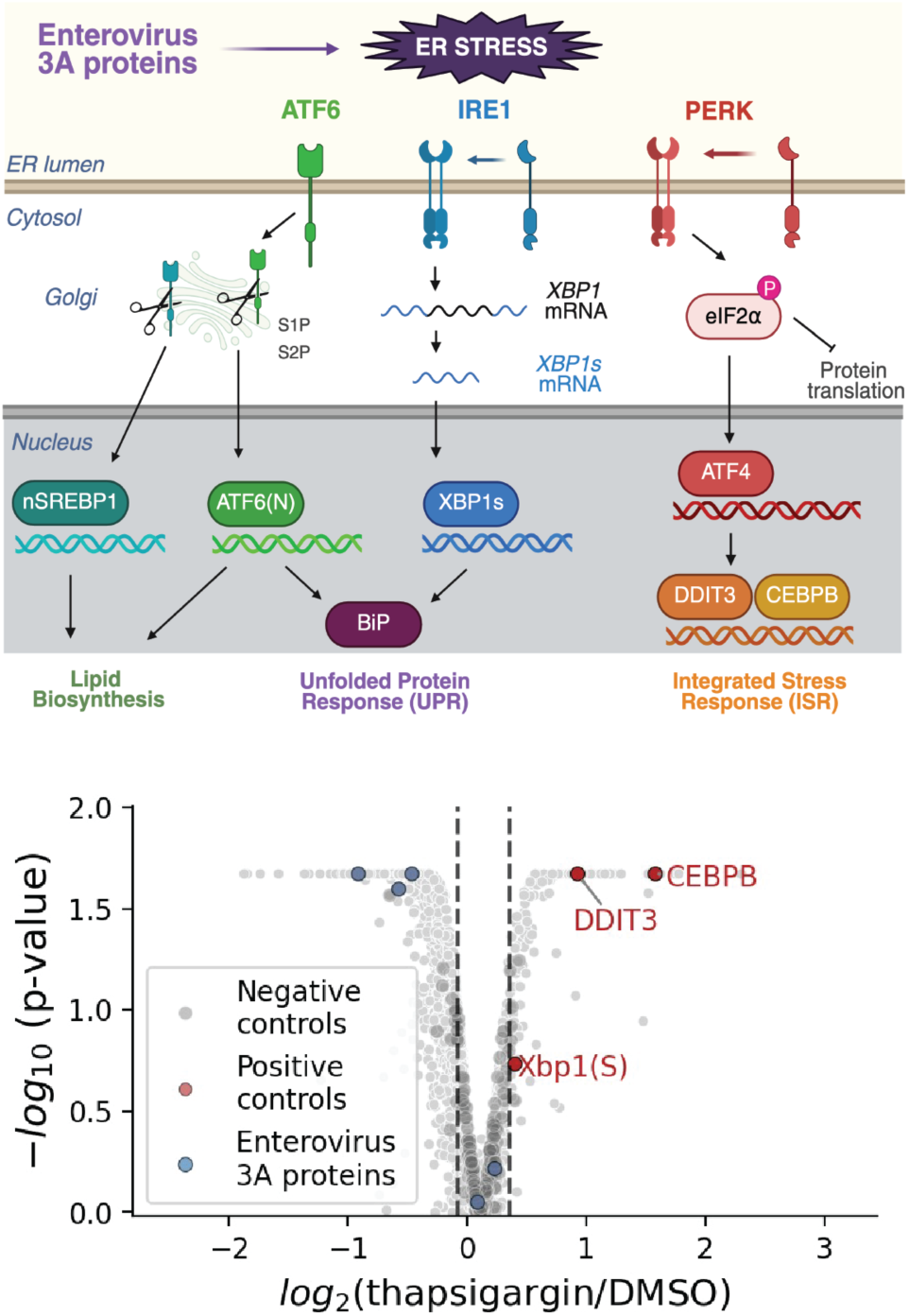
Stress screens identify known and novel viral modulators of protein stress. (**A**) The unfolded protein response (UPR) pathway. (**B**) Volcano plot of the 4-day protein-stress growth screen under thapsigargin treatment compared with DMSO control. Positive controls relevant to the UPR are highlighted.

**Fig. S9.**
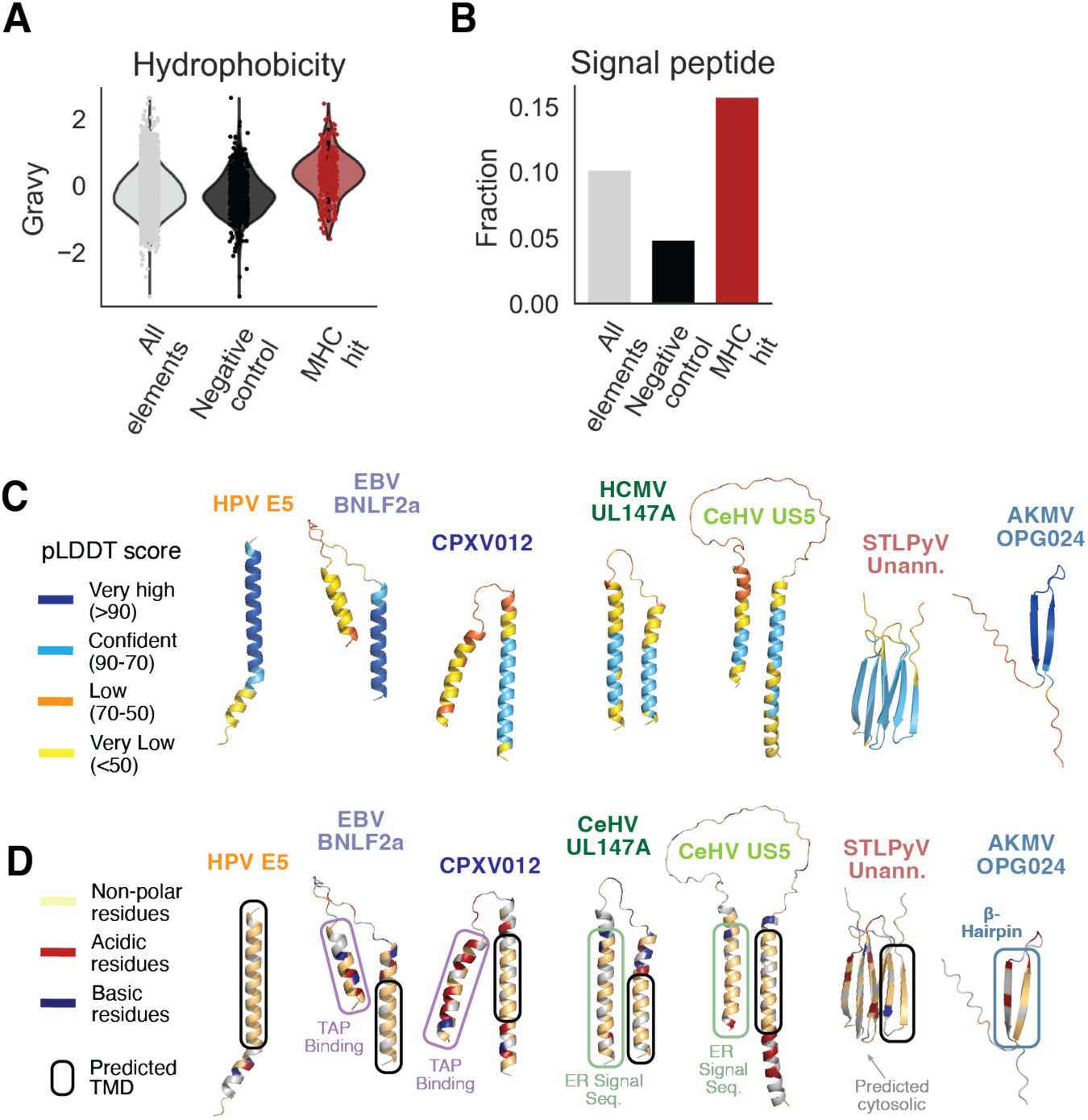
Sequence and structures of MHC screen hits. (**A**) Violin plot of GRAVY hydrophobicity scores. (**B**) Fraction of elements with an ER signal-peptide sequence predicted by PHOBIUS. AF3-predicted structures of MHC-I inhibitors colored by (**C**) pLDDT score and (**D**) amino-acid property.

**Fig. S10.**
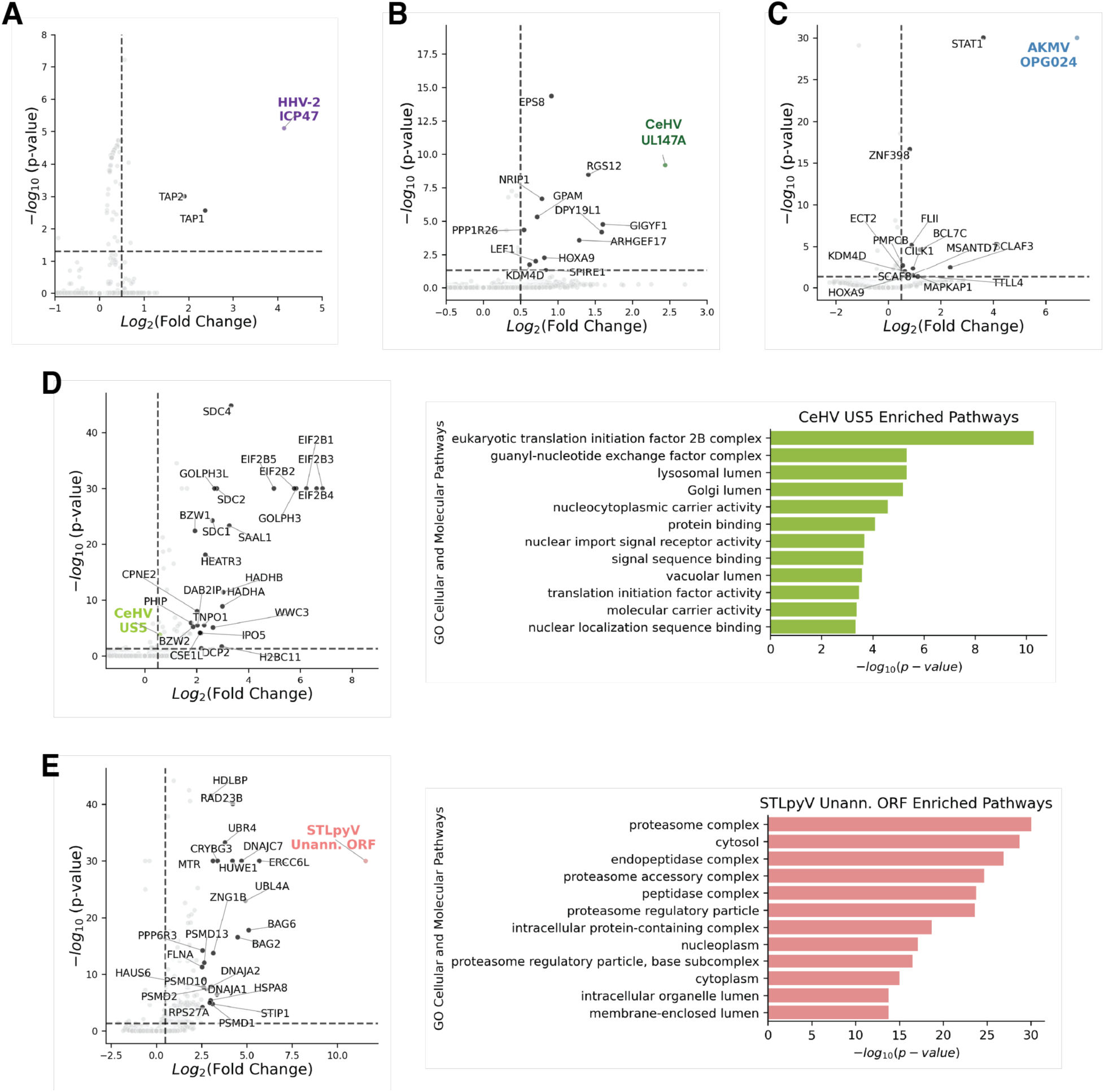
Identification of host interactors of MHC inhibitors. Volcano plots of AE-MS performed on 293F cells transfected with C-term ALFA-tagged MHC inhibitors compared with ALFA-tagged mCherry or GFP controls for (**A**) HHV-2 ICP47, (**B**) CMV UL147A, (**C**) AKMV OPG024, (**D**) CeHV US5, and (**E**) an unannotated STLpyV smORF. GO pathway enrichment was also performed on all significant interactors of CeHV US5 and STLpyV smORF.

**Fig. S11.**
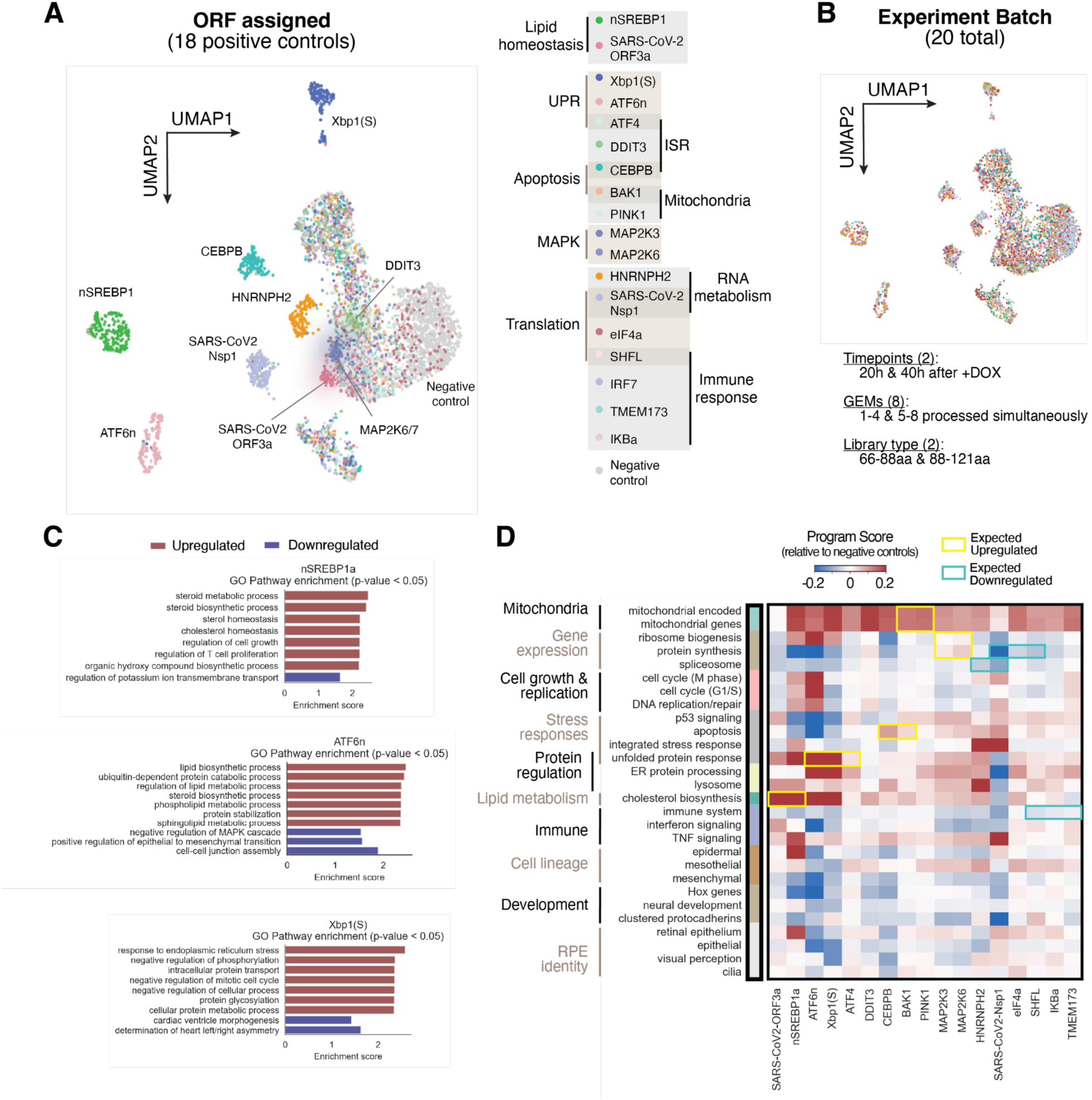
Perturb-seq positive controls have expected phenotypes. UMAP of single cells assigned to positive and negative control ORFs, colored by (**A**) ORF assignment and (**B**) experiment batch, which includes batches divided up by timepoint, GEM group, and library type. (**C**) Gene set enrichment analysis of all DEGs in example positive controls. (**D**) Heatmap of program scores derived from RPE-1 essential-gene Perturb-seq relative to negative controls.

**Fig. S12.**
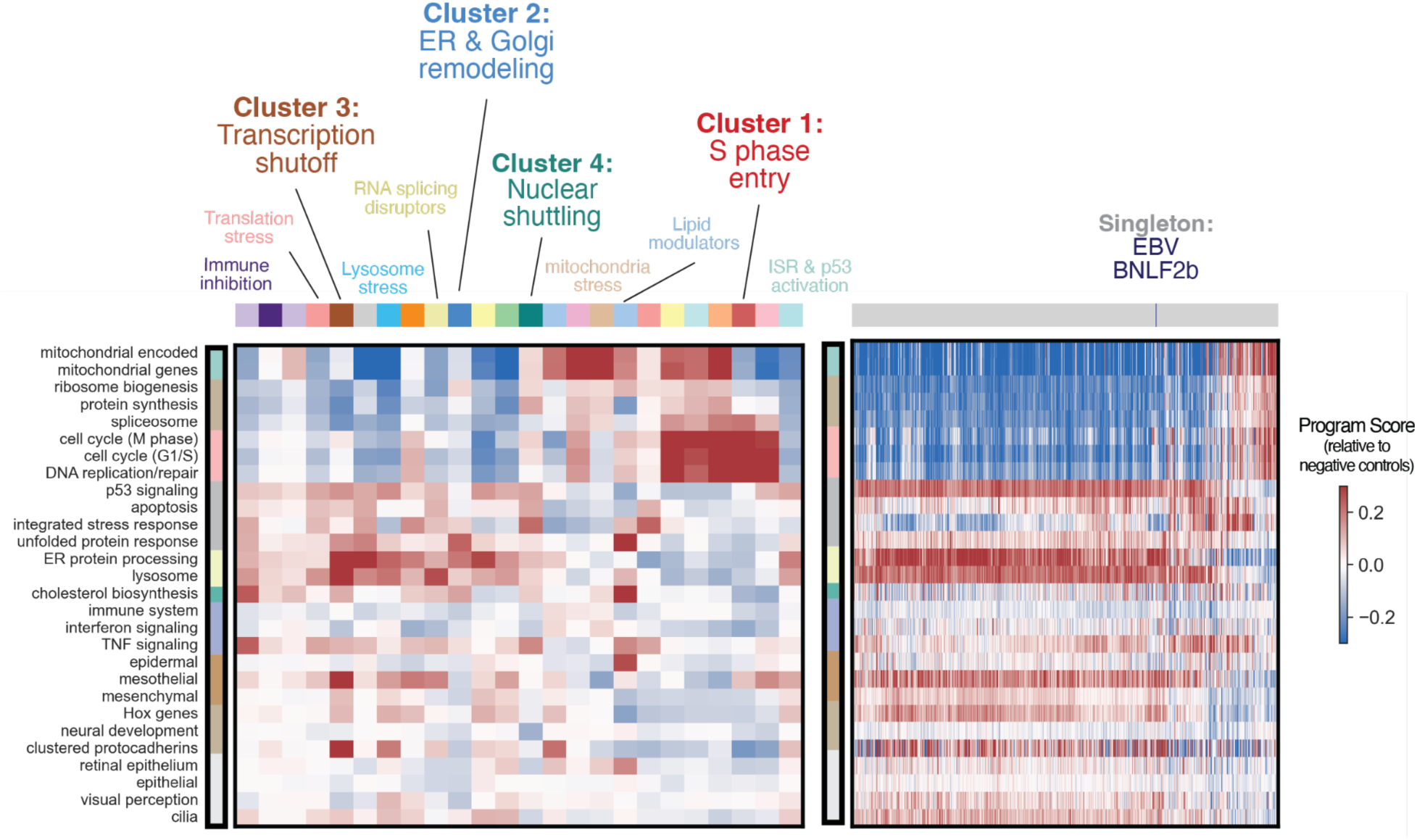
Gene pathway scoring from transcriptional profiles. Heatmap of program scores for (Left) Perturb-seq clusters and (Right) singletons that pass the energy-distance test, with EBV BNLF2b marked.

**Fig. S13.**
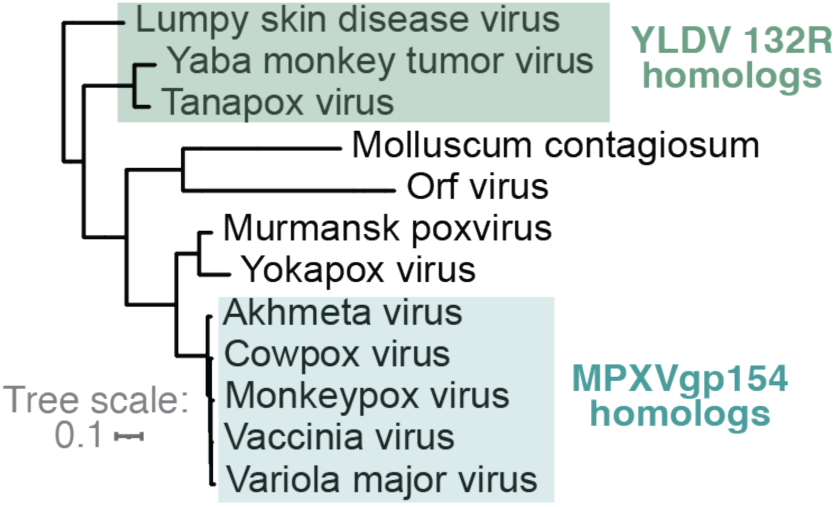
Phylogenetic tree of poxvirus lineage. Trees were obtained from ICTV with presence of MPXVgp154 and YLDV 132R homologs highlighted.

**Fig. S14.**
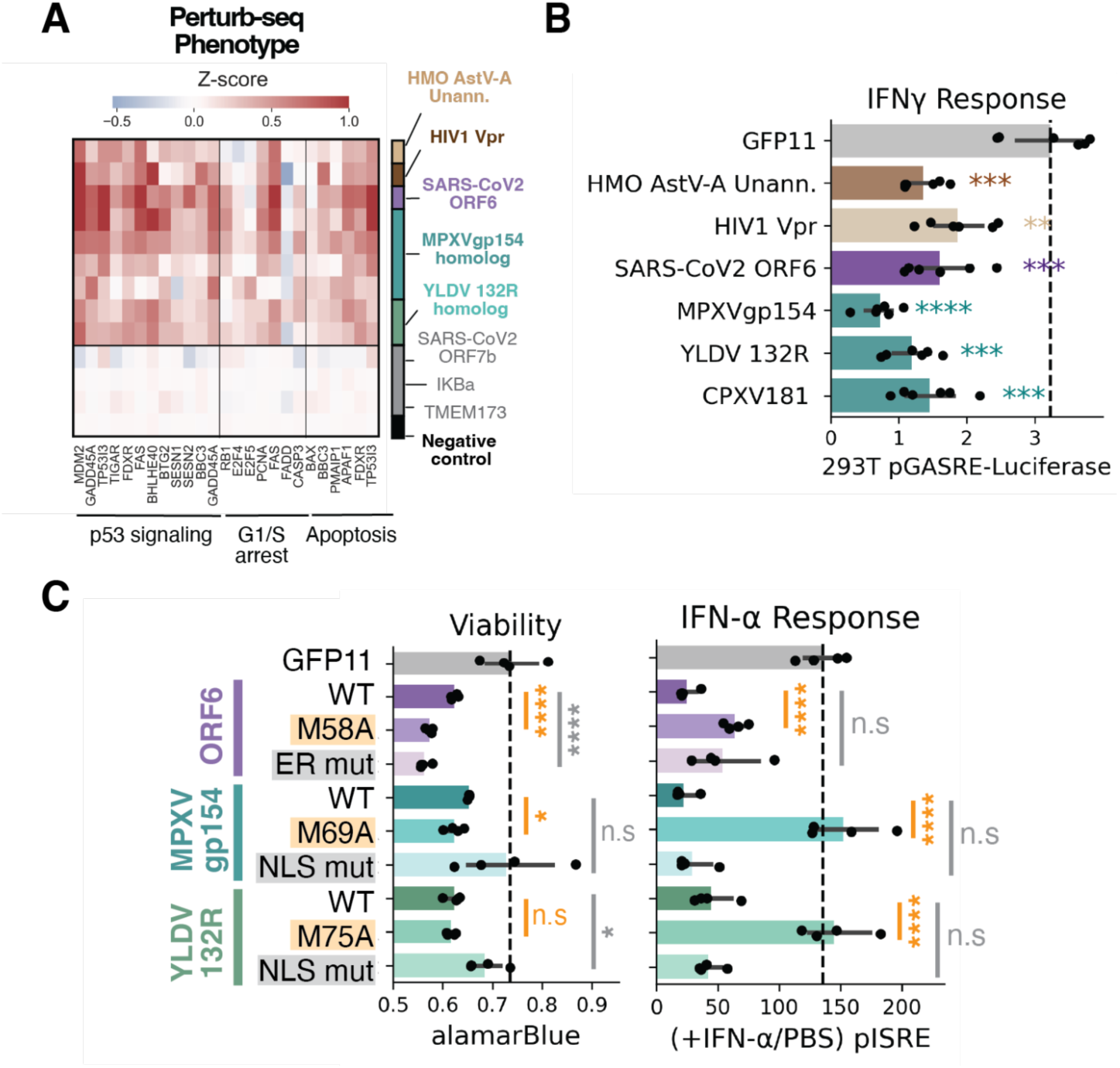
Phenotypic characterization of cluster 4, the nuclear shuttling inhibitors. (**A**) Heatmap of DEGs grouped by pathway. (**B**) IFN-γ response upon overexpression in HEK293T cells, measured with a pGAS-RE-NanoLuc assay after 16 h of stimulation with 200 ng/mL IFN-γ. (**C**) HEK293T cells transfected with the indicated WT and mutant ORFs were examined for viability (left) by alamarBlue assay and for IFN-α response (right) by pISRE-NanoLuc assay after 16 h of stimulation with 75 ng/mL IFN-α.

**Fig. S15.**
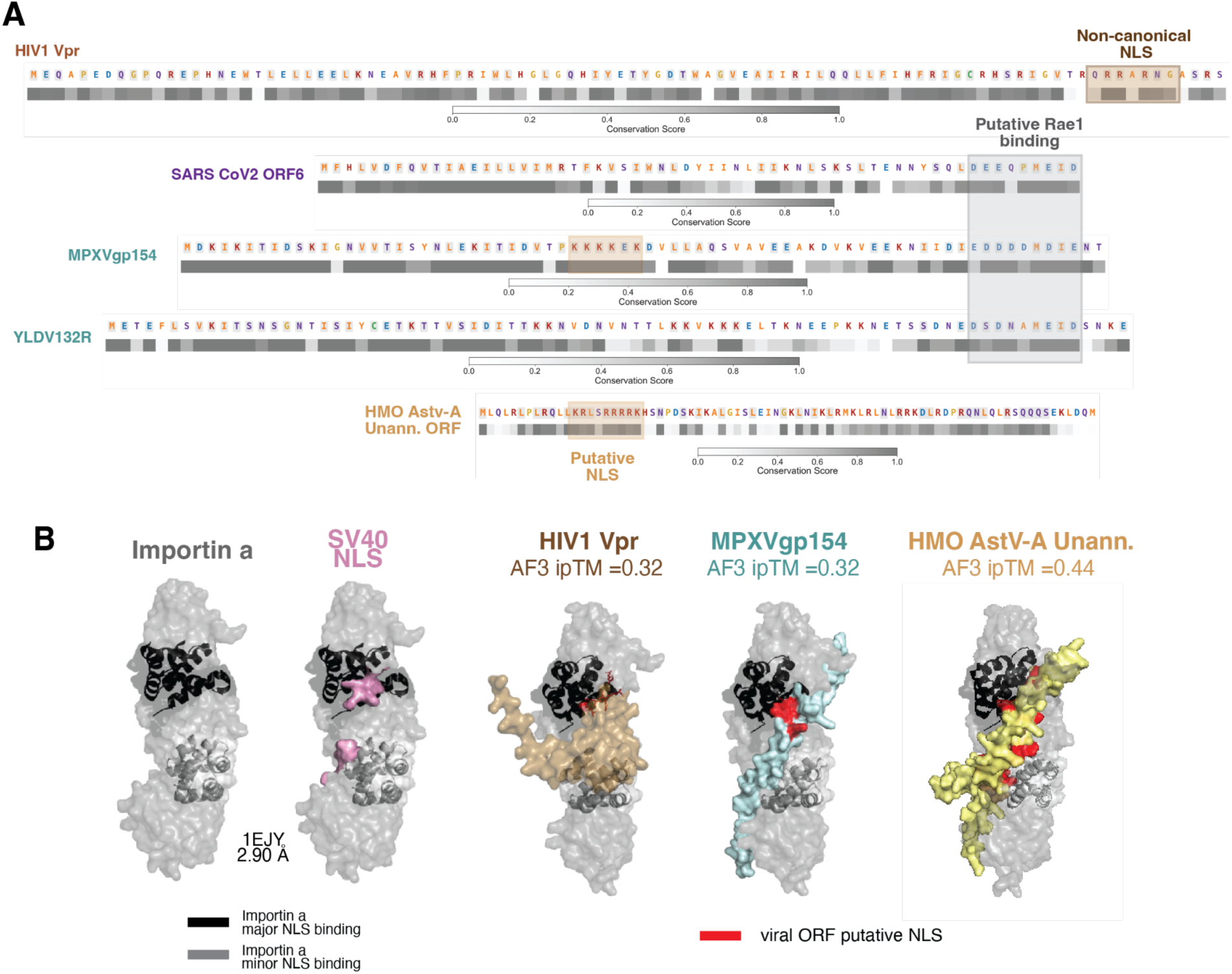
Identification of putative NLSs in nuclear shuttling inhibitor cluster elements. (**A**) Sequence analysis and conservation scores of cluster elements, with putative NLS and putative Rae1-binding motifs highlighted. (**B**) Comparison of importin-α bound to the SV40 NLS (PDB: 1EJY) with AF3-predicted structures of importin-α bound to indicated ORFs.

**Fig. S16.**
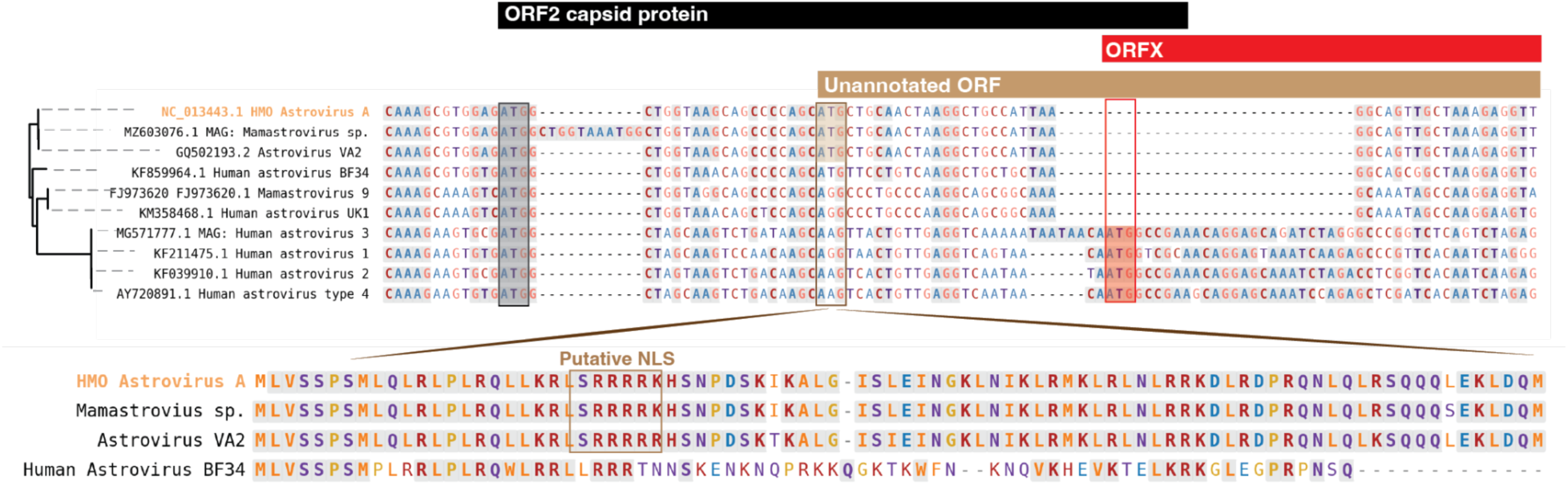
Astrovirus genome locus and protein sequences of the HMO AstV-A unannotated ORF across lineages. MSA of astrovirus genomic loci containing HMO AstV-A Unann. ORF with its +1 start codon highlighted, and comparison of the amino-acid sequences of smORFs sharing this +1 start codon.

**Fig. S17.**
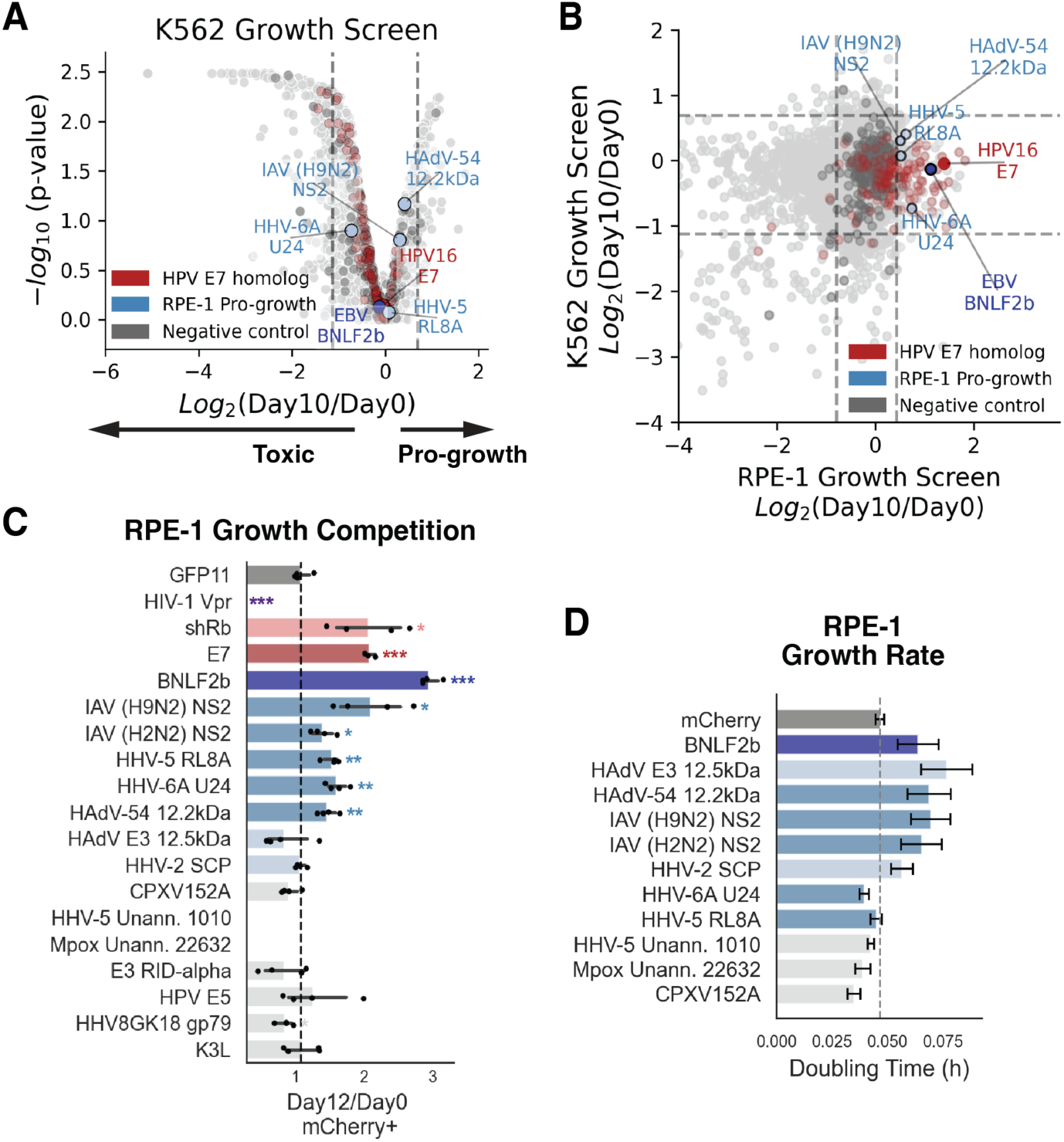
RPE-1 pro-growth hits do not promote growth in K562 cells but validate in RPE-1 cells. (**A**) Volcano plot of the 10-day K562 growth screen with pro-growth hits highlighted. (**B**) Scatter plot of K562 versus RPE-1 growth screen phenotypes with pro-growth hits highlighted. (**C**) Arrayed growth-competition and (**D**) arrayed growth-rate measurements by confluency tracking of RPE-1 cells overexpressing the indicated ORFs.

**Fig. S18.**
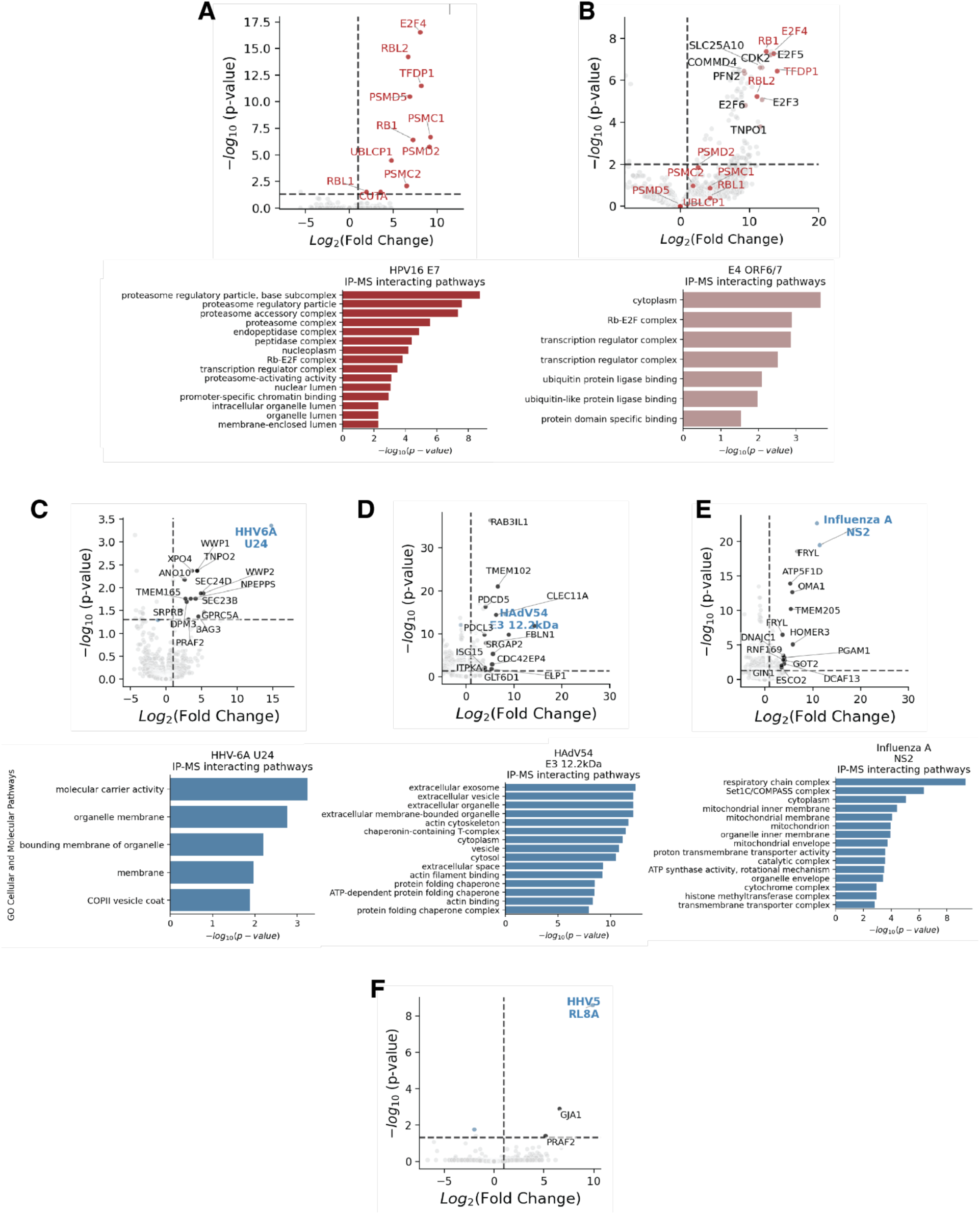
Identification of host interactors of validated RPE-1 pro-growth hits. Volcano plots of AE-MS performed on RPE-1 cells overexpressing ALFA-tagged pro-growth hits compared with ALFA-tagged mCherry or GFP controls, and GO pathway enrichment of significant interactors, for (**A**) HPV16 E7, (**B**) HAdV E4 ORF6/7, (**C**) HHV-6A U24, (**D**) HAdV54 E3 12.2 kDa, (**E**) IAV NS2, and (F) HHV-5 RL8A.

**Fig. S19.**
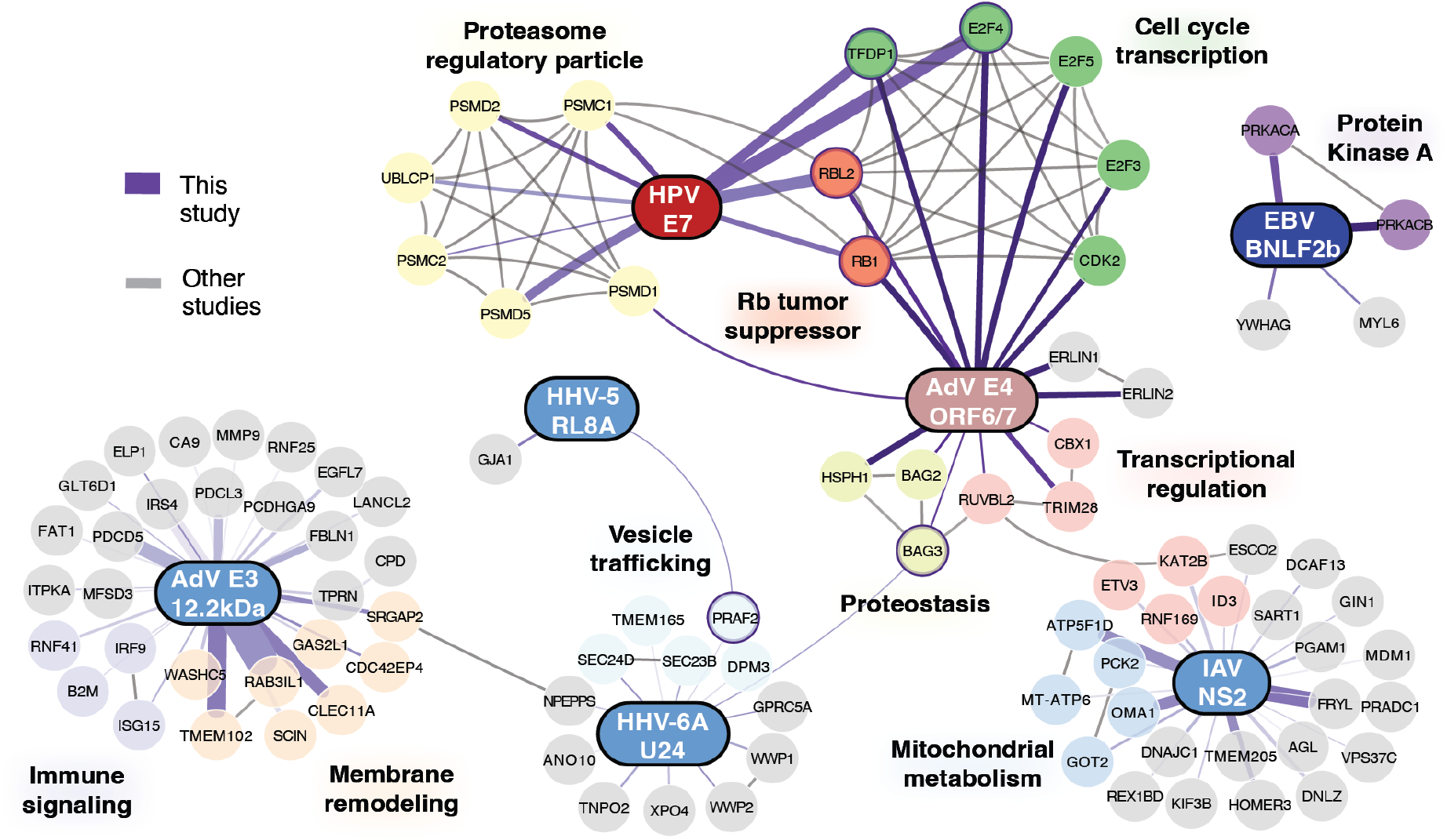
Protein interaction networks of pro-growth hits align with Perturb-seq cluster and singleton assignments. Networks were generated by integrating our AE-MS interactors with host protein networks derived from STRING, with edges drawn if the interaction confidence score is >0.75. Functions of host-interactor groups were manually annotated.

**Fig. S20.**
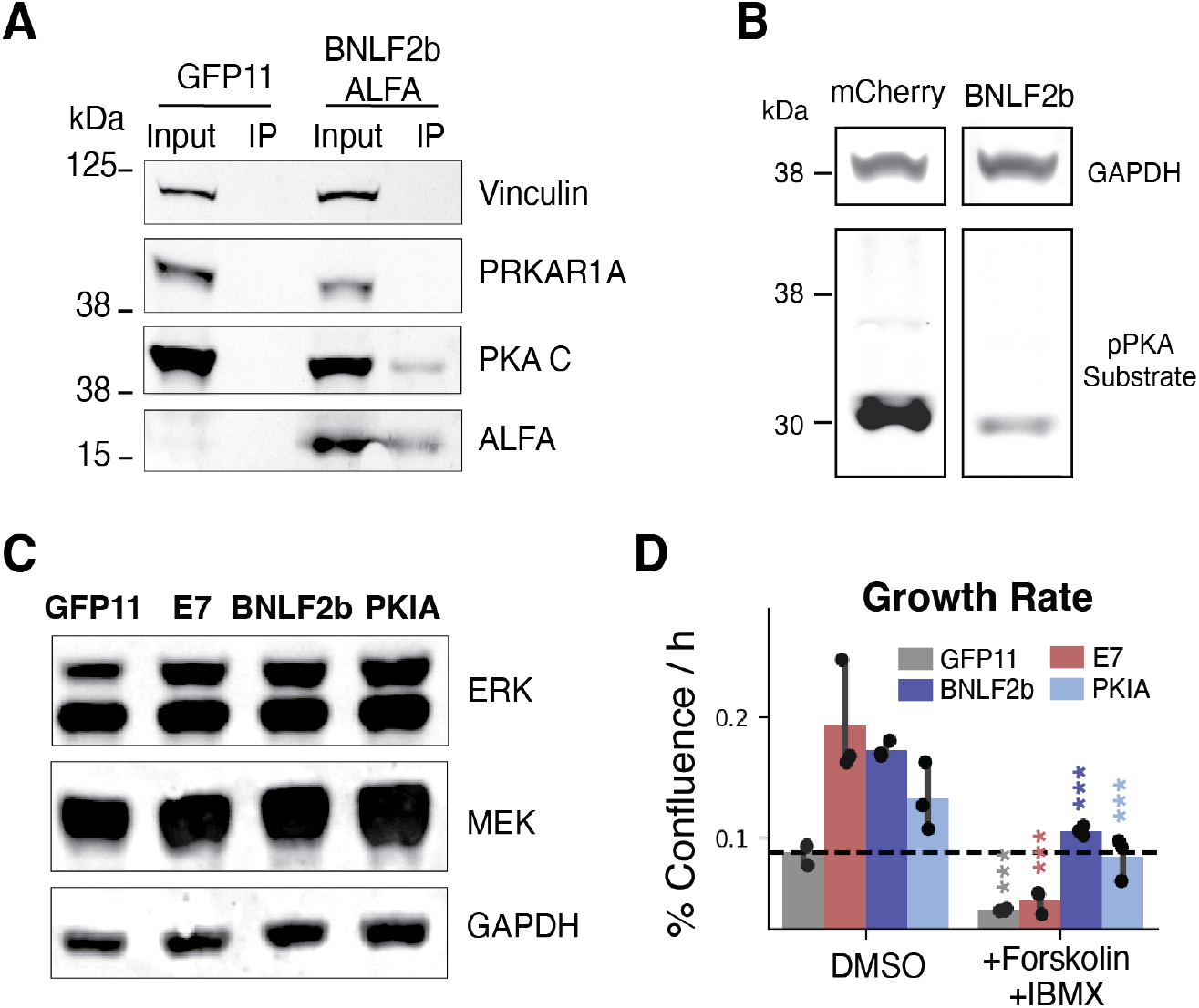
Further characterization of BNLF2b modulation of PKA and MAPK. (**A**) IP-Western of A375 cells overexpressing ALFA-tagged BNLF2b for examining interaction with PRKAR1A and PKA Cα (PRKACA). (**B**) Western blot of pPKA substrate levels in mCherry and BNLF2b overexpressing RPE-1 cells. (**C**) Total ERK and MEK levels of indicated ORFs overexpressed in RPE-1. (**D**) Growth rates of RPE-1 cells overexpressing the indicated ORFs, grown either in DMSO vehicle-control medium or in medium containing forskolin and IBMX.

**Fig. S21.**
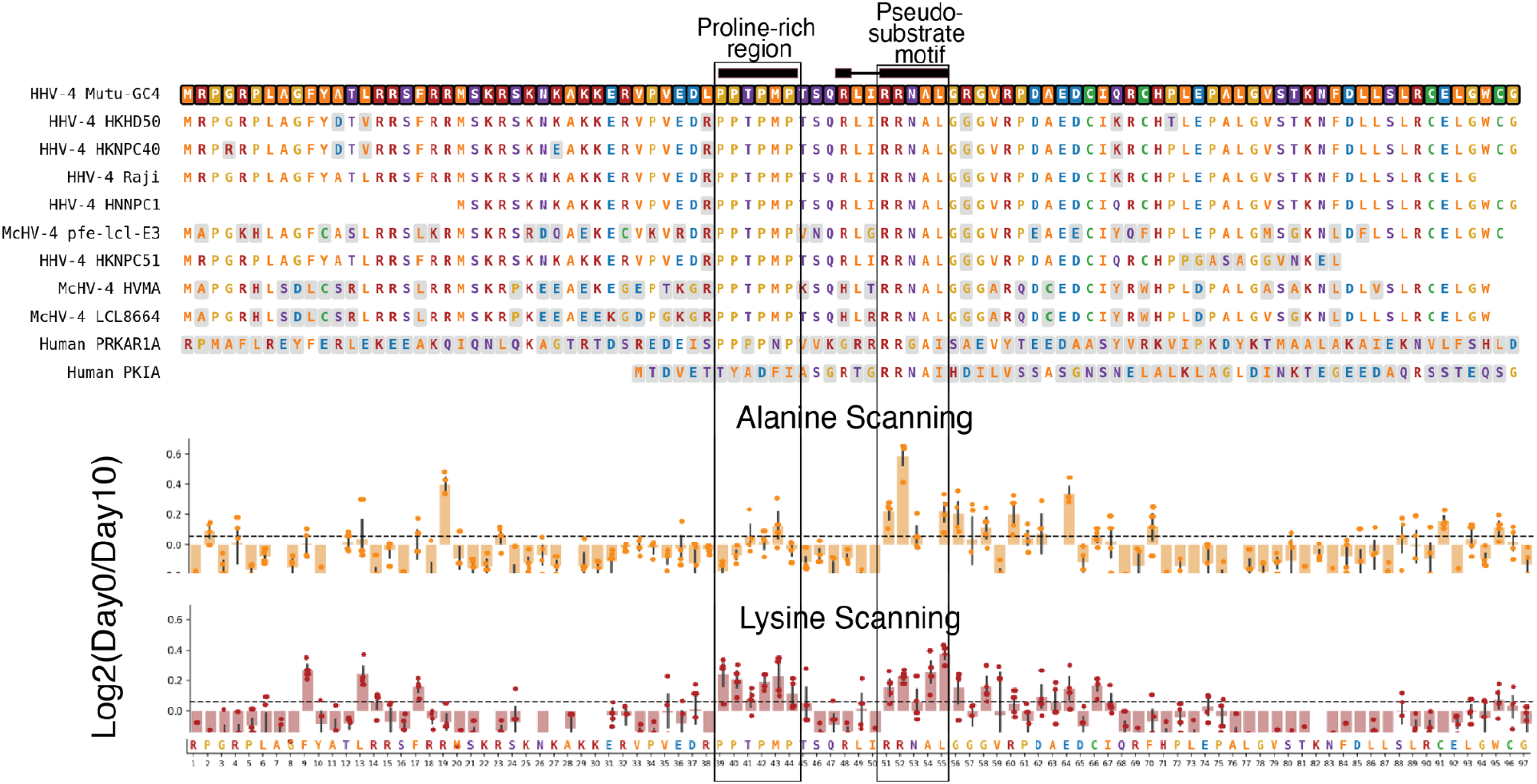
**BNLF2b mutagenesis screens identify key BNLF2b residues that impact growth. (**Top) MSA of BNLF2b homologs. Residues are aligned with the results of (Middle) alanine and (Bottom) lysine scanning from 10-day RPE-1 growth screens.

**Fig. S22.**
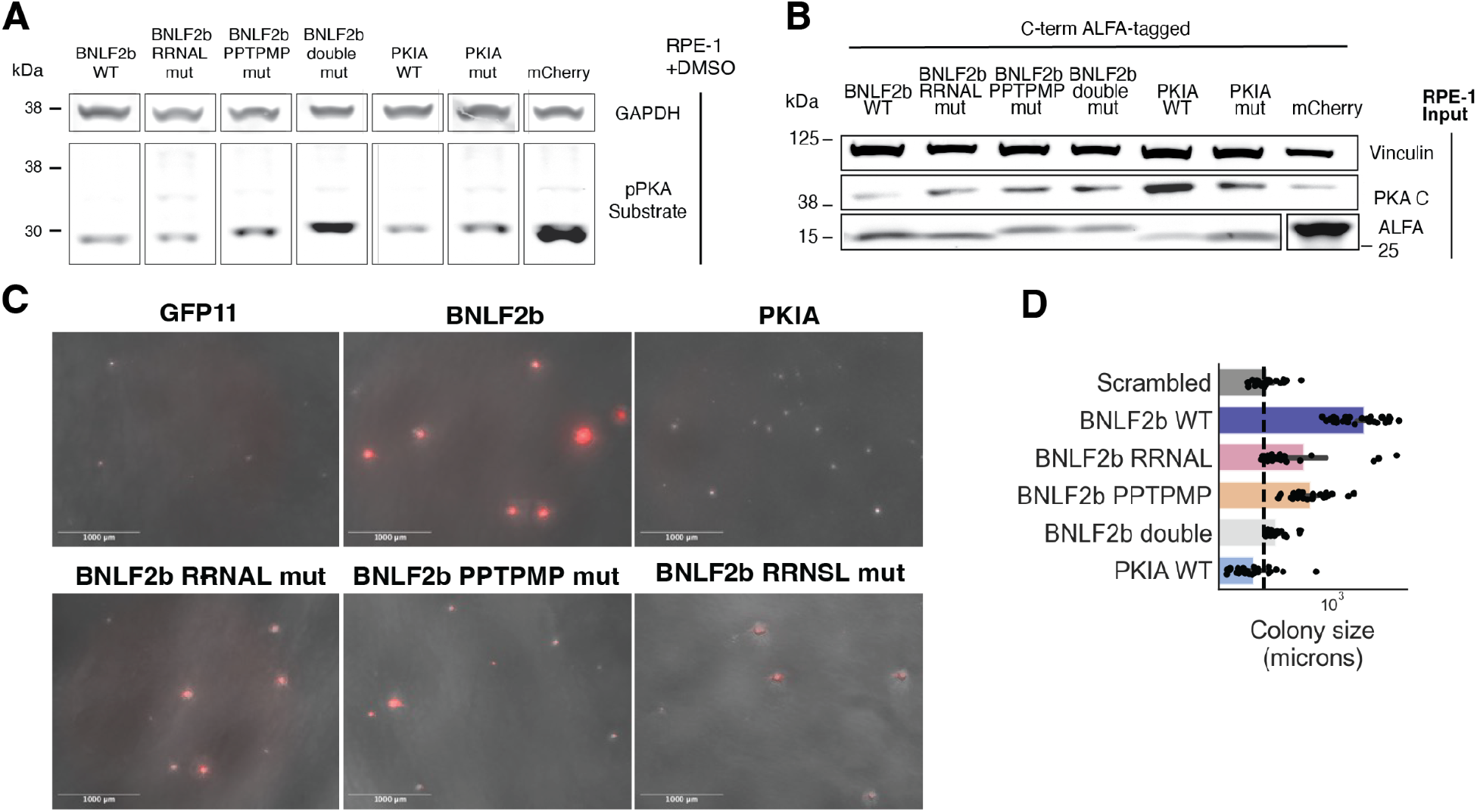
Further characterization of BNLF2b mutants. (**A**) pPKA substrate levels in RPE-1 cells overexpressing the indicated ORFs. (**B**) Input for the IP-MS of ALFA-tagged BNLF2b and PKIA (WT and mutants). (**C**) Soft-agar transformation assay 45 days after seeding of RPE-1 cells lentivirally transduced with constitutive overexpression constructs for the indicated ORFs. Images show phase contrast overlaid with mCherry fluorescence. (**D**) Quantification of colony sizes.

**Fig. S23.**
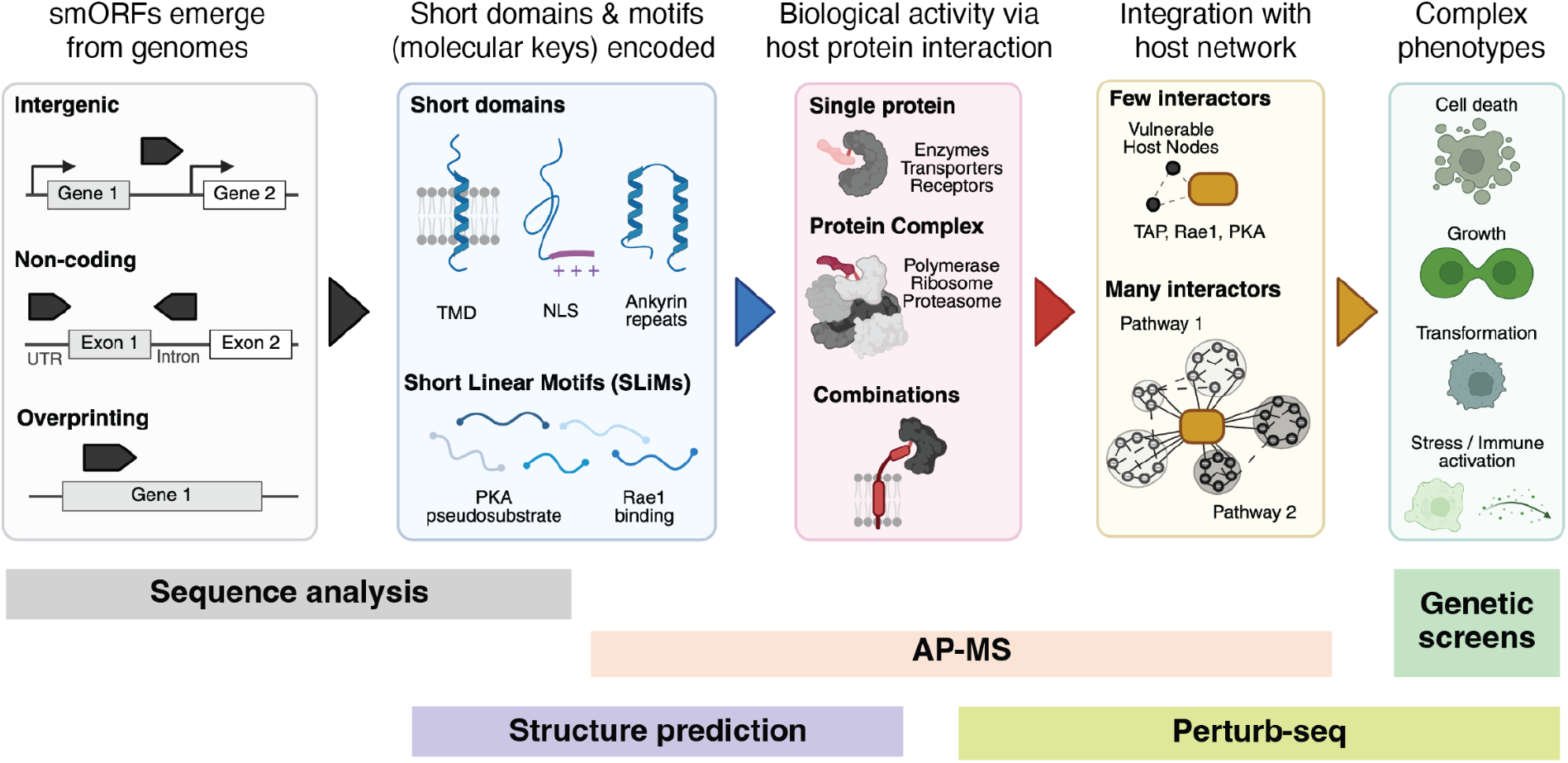
A model for emergence of functional smORFs. smORFs emerge in viral genomes in intergenic or non-coding regions as well as overlapping in alternative frames from other coding sequences by overprinting. smORFs generate microproteins containing short domains or SLiMs. Molecular keys enable interactions with host proteins and give rise to biological activities. Combinations of keys further enhance potency or give rise to multifunctionality. Integration with the host protein network modulates host pathways that result in emergence of complex cellular phenotypes.

**Table S1.**
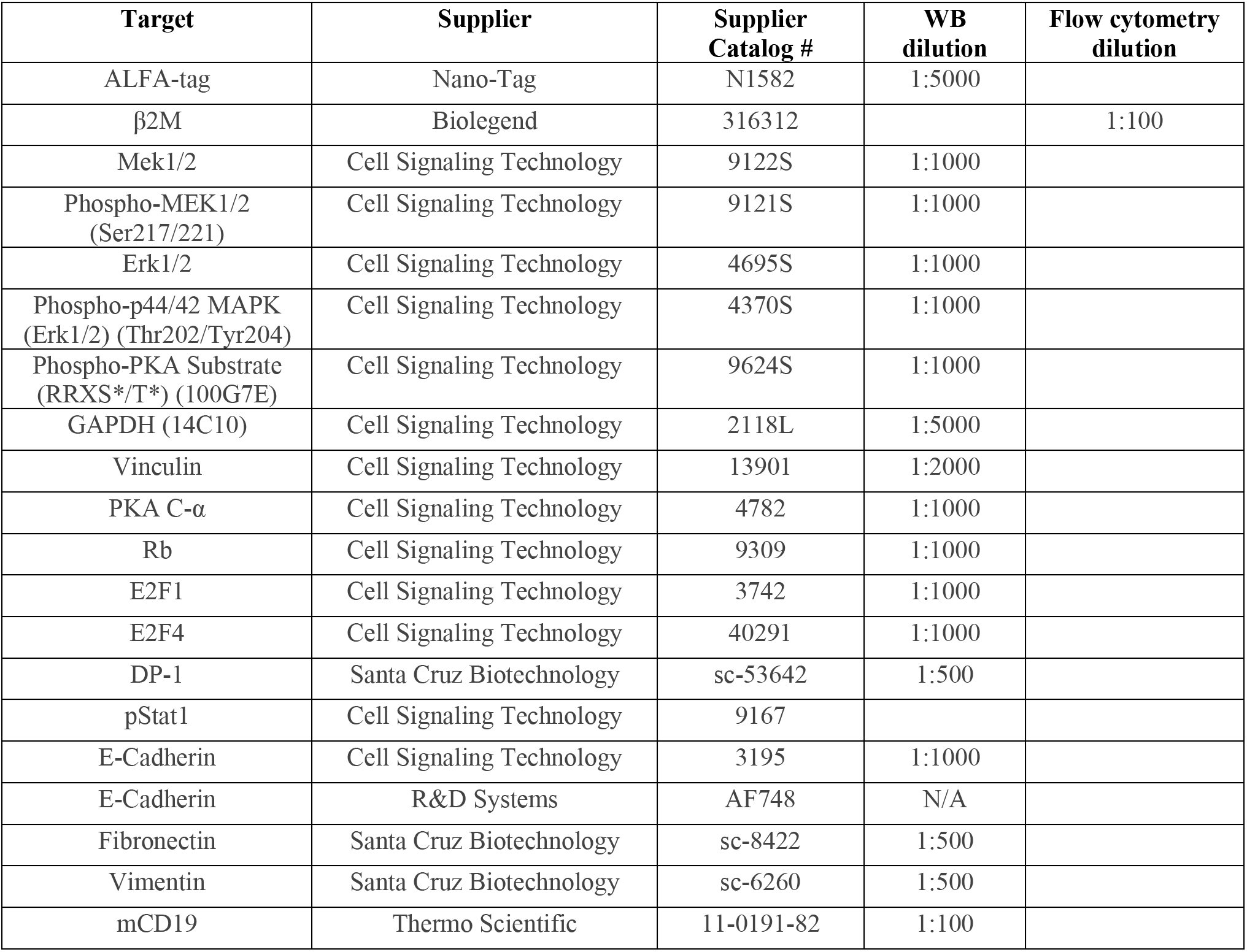
Antibodies used for western blotting and flow cytometry.

**Table S2.**
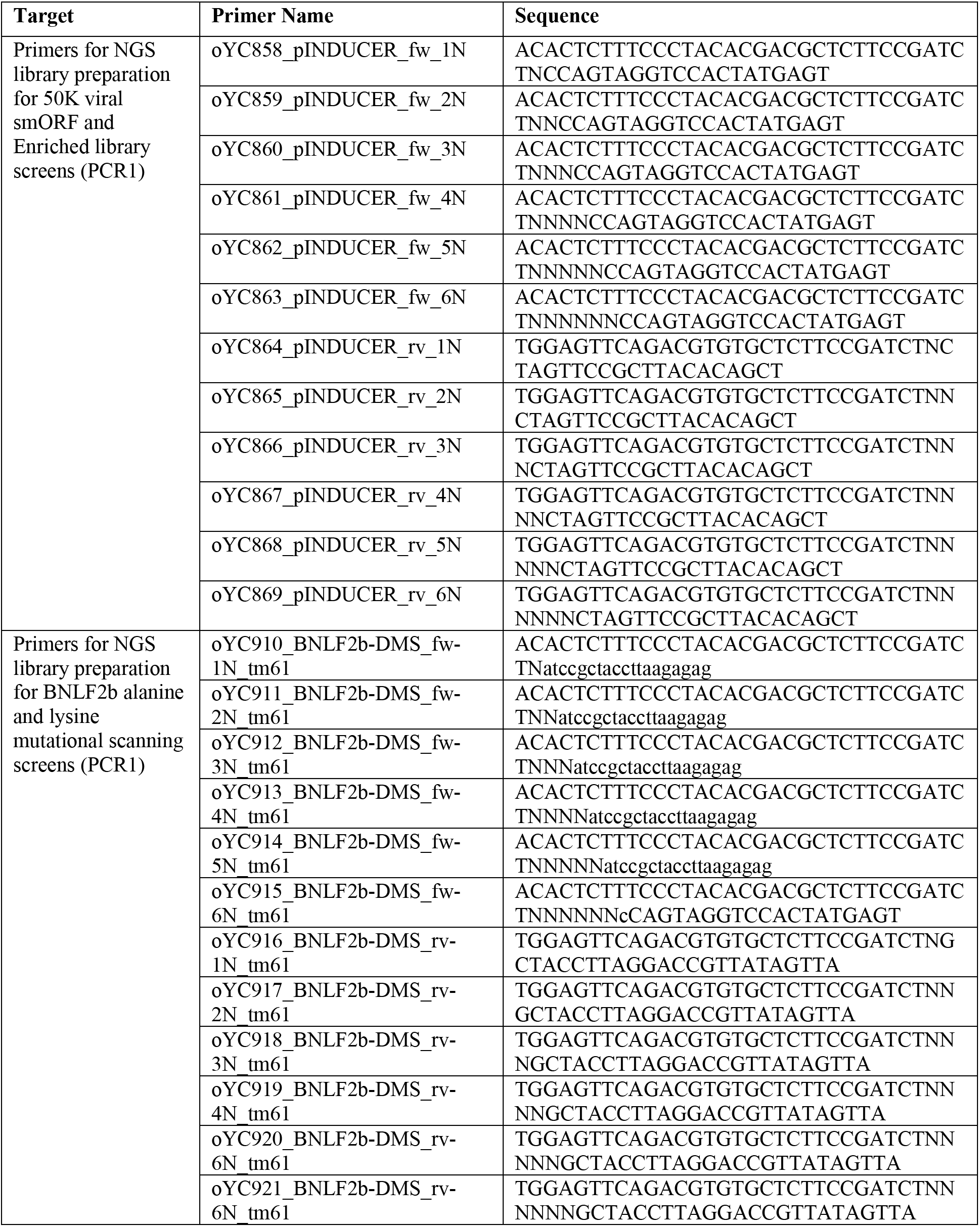
Primers for genetic screens.

## Data S1. Design and annotation of the viral microprotein libraries

Sheet S1a lists all 52,979 elements of the 50K viral smORF library and sheet S1b all 4,107 elements of the enriched library, with source virus, accession, genomic coordinates, start codon, annotation status, encoded amino acid sequence, subpool assignment and control class.

## Data S2. Pooled 10-day growth screens with the pANDORA 50K and the Enriched library of viral smORFs

Sheets S2a and S2b give per-element read counts, normalized abundances, log₂ fold changes, Mann-Whitney P values and Benjamini-Hochberg-adjusted P values for the K562 and RPE-1 screens, respectively. Sheets S2c and S2b give the same quantities as S2a and S2b for the K562 and RPE-1 enriched-library screens.

## Data S3. Pooled screens for all targeted secondary screens

Sheet S3a, MHC-I surface expression screen in A375; sheets S3b and S3c, MHC-I screens in RPE-1 without and with interferon-γ stimulation; sheets S3d and S3e, survival screens in K562 under ER stress induced by thapsigargin and brefeldin A, respectively.

## Data S4. Perturb-seq summary statistics

Energy distance to the negative controls, energy-test FDR, number of differentially expressed genes and cluster assignment for 3,412 perturbations passing the post-QC criteria for perturb-seq processing.

## Data S5. Affinity-enrichment and immunoprecipitation mass spectrometry

Sheets S5a to S5d report log₂ fold changes and Benjamini-Hochberg-adjusted P values for every bait–prey pair underlying Figs. 2, 3, 4 and 5, respectively.

## Data S6. Fixed Perturb-seq custom probe set for ORF calling

Custom probe sequences for fixed Perturb-seq with 10X Flex Gene Expression technology.

